# Pathogen-driven coevolution across CBP60 plant immune regulator subfamilies confers resilience on the regulator module

**DOI:** 10.1101/2020.07.16.207134

**Authors:** Qi Zheng, Kristina Majsec, Fumiaki Katagiri

## Abstract

Among eight Arabidopsis CaM-Binding Protein (CBP) 60 family members, AtCBP60g and AtSARD1 are partially functionally redundant, major positive immune regulators while AtCBP60a is a negative immune regulator. Phylogenetic analysis of CBP60 protein sequences of 247 diverse land plant species indicated that the CBP60a, CBP60g, and SARD1 immune regulator subfamilies diversified around the time of Angiosperm divergence. The immune regulator subfamilies, especially the CBP60g subfamily, have been evolving very fast, suggesting strong selection pressure from CBP60-targeting pathogen effectors. We closely examined this fast evolution of the immune regulator subfamilies using the subfamily protein sequences from 12 diverse Core Eudicot species. The fast evolution was caused by both high proportions of polymorphic sites and high evolution rates per polymorphic site, compared to the genomic norm. We developed an analytical platform for physical-chemical characteristics of amino acids, called Protein Evolution Analysis in Euclidean Space (PEAES). Using the pairwise distance rank on PEAES (PEAES-PDR), we detected signatures of significant coevolutionary interactions across the immune regulator subfamilies within the CBP60-conserved domain. The patterns of detected coevolutionary interactions were consistent with hypothetical coevolutionary mechanisms that protect the positive immune regulator function from targeting by pathogen effectors. The coevolutionary interactions across the subfamilies were largely specific to each species lineage, suggesting that the coevolution occurred through species lineage-specific histories of variable pathogen pressure. Thus, fast coevolution of the subfamilies with overlapping or opposing functions appears crucial to maintain resilience of the CBP60 immune regulator module against fast-evolving pathogen effectors.

## INTRODUCTION

Pathogen effectors, typically proteinaceous, are delivered from pathogens into plant cells and manipulate plant functions to improve pathogen environments *in planta*, including compromising plant immune signaling (Toruño et al. 2016). The plant immune signaling network characterized in Arabidopsis has a remarkable level of resilience as different parts of the network have compensating functions that buffer the disabling impacts of mutations or pathogen effectors (Tsuda et al. 2009; Hillmer et al. 2017; Katagiri 2018). We are interested in how such network resilience evolved in biological systems. To gain insights into this topic, we have been studying evolution of important components of the immune signaling network. Members of the CaM-Binding Protein (CBP) 60 family are such network components (Wang et al. 2009; Zhang et al. 2010; Wang et al. 2011; Truman et al. 2013).

Arabidopsis has eight CBP60 family members (AtCBP60a-g and AtSARD1) (Reddy et al. 2002; Zhang et al. 2010; Wang et al. 2011). The CBP60 protein family was defined by the domain conserved among known CBP60 proteins (Pfam: “calmodulin_bind”, PF07887). Despite the Pfam domain name, the actual CaM-binding domains are outside the conserved domain (Fig 1) (Reddy et al. 2002; Wang et al. 2009; Wang et al. 2011). Thus, we call the conserved domain the CBP60-conserved domain. AtCBP60g and AtSARD1 are major positive regulators of immunity controlling activation of many immune responses, including synthesis of the important immune hormone salicylic acid (SA) (Wang et al. 2009; Zhang et al. 2010; Wang et al. 2011). They are transcriptionally induced during immune responses. Their functions as positive immune regulators are partially redundant since an *atcbp60g atsard1* double mutant has a more severe immune deficiency than either of the single mutants (Zhang et al. 2010; Wang et al. 2011). In contrast, AtCBP60a is a negative regulator of SA signaling and immunity (Truman et al. 2013). In an *atcbp60a* mutant the basal level of SA is higher (Truman et al. 2013) and decrease of the post-induction level of AtCBP60g mRNA is slowed (Lu et al. 2018). The functions of the other AtCBP60 members (AtCBP60b-f) are unknown as mutants lacking any one of them did not show substantial changes in immunity or other obvious phenotypes (Truman et al. 2013). AtCBP60g and AtSARD1 are DNA-binding proteins (Zhang et al. 2010; Qin et al. 2018). Since the DNA-binding activity resides within the CBP60-conserved domain, it is likely that all CBP60 proteins are DNA-binding proteins. Phylogenetic analysis of AtCBP60 members with a CBP60 from the moss *Physcomitrella patens* as the outgroup suggested that the immune regulators AtCBP60a, AtCBP60g, and AtSARD1 form one clade and AtCBP60b-f form a separate clade (Wang et al. 2011).

**Fig 1.**
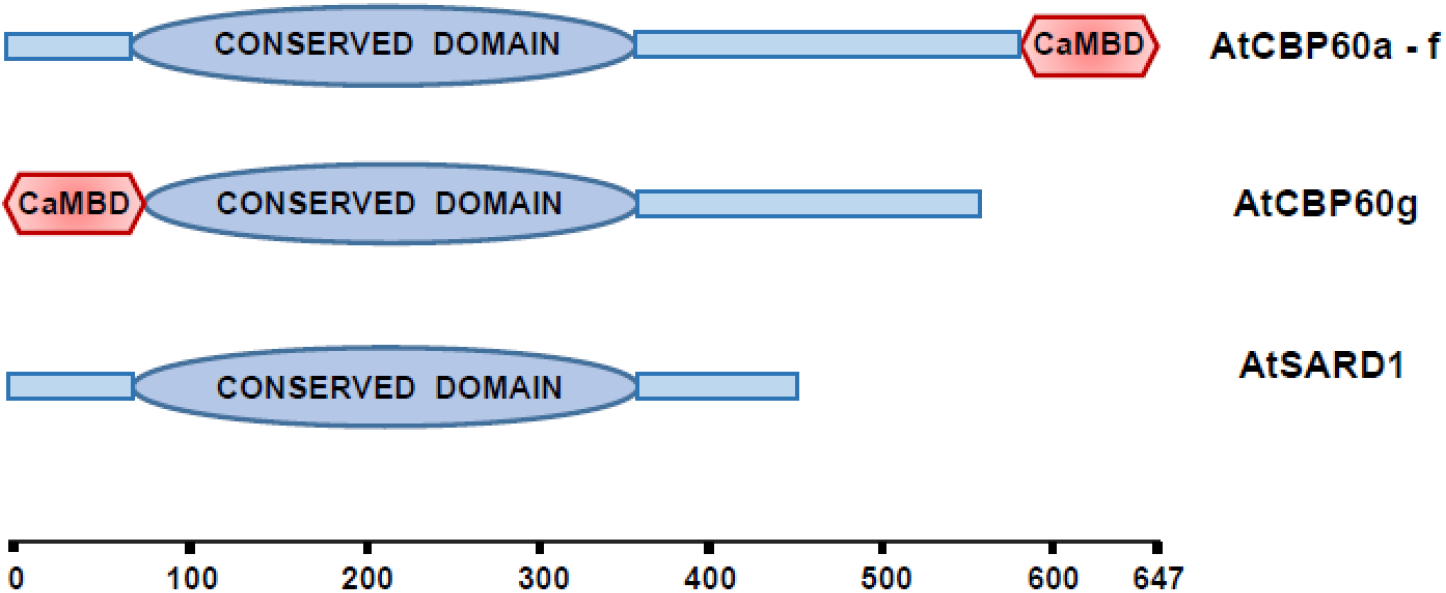
Domain structures of AtCBP60 proteins. The amino acid lengths between the CBP60-conserved domain and the C-CaMBD in AtCBP60a-f vary from 133 to 261: in this figure, the length for AtCBP60b was used. CaMBD, calmodulin-binding domain.

We applied phylogenetic analysis to the CBP60 protein family members in land plants and found that the immune-related clade diversified from the highly conserved, prototypical group around the time of Seed Plant divergence. The CBP60a, CBP60g, and SARD1 immune regulator subfamilies diversified within the immune-related clade around the time of Angiosperm divergence. The immune regulator subfamilies have been evolving very fast, which suggests strong selection imposed by fast-evolving pathogen effectors. Among 12 diverse Core Eudicot species, the fast evolution of the immune regulator subfamilies was caused by both higher proportions of polymorphic amino acid sites and fast evolution rates per polymorphic site, compared to the genomic norm represented by benchmark protein sequences.

We further investigated possible coevolutionary interactions across these fast-evolving CBP60 immune regulator subfamilies. We developed a geometric analysis platform for the physical-chemical characteristics of amino acids, Protein Evolution Analysis in Euclidean Space (PEAES). The pairwise distance rank on PEAES (PEAES-PDR) was used to examine possible coevolutionary interactions among the three immune regulator subfamilies in the CBP60-conserved domain in the 12 Core Eudicot species linages. We detected significant coevolutionary interactions, which were consistent with two coevolutionary mechanism hypotheses. First, two positive regulators evolve to be as different as possible, which makes it difficult for a single pathogen effector to target both at once. Second, the negative regulator evolves to retain similarity to the positive regulators at the sites where the positive regulators cannot be very different. If a pathogen effector targets both positive and negative regulators, the negative impact of the effector on immunity due to targeting the positive regulators is moderated by its simultaneous targeting of the negative regulator. Our discoveries strongly suggest that functionally related protein subfamilies influence the evolution of each other so that high levels of functional resilience can be achieved under strong and rapidly varying selection imposed by fast-evolving pathogen effectors.

## NEW APPROACHES

We adapted a description of physical-chemical characteristics of amino acids in an Euclidean space by Venkatarajan and Braun (Venkatarajan and Braun 2001) to evaluate the effect of selection. We call this geometric analysis platform PEAES. To detect coevolution of protein subfamilies after diversification of the subfamilies, we used the rank of the pairwise PEAES distance between the members of two subfamilies of a particular species among all species permutations between the two subfamilies (PEAES-PDR). The distributions of PEAES-PDR between two pairs of the protein subfamilies should be uncorrelated if evolution of each subfamily is independent. Detection of a significant correlation indicates coevolution of the subfamilies in the particular species lineage.

## RESULTS

### The immune-related CBP60 clade neofunctionalized from the prototypical CBP60 group

We inferred phylogenetic relationships among 1024 CBP60 protein sequences from 247 diverse land plants: one Liverwort, 35 Moss, 9 Lycophyte, 44 Fern, 69 Gymnosperm, and 89 Angiosperm species (Figs 2 and S1; the latter includes sequence names). Most plant species had at least one member of the highly conserved, “prototypical” CBP60 group that includes AtCBP60b-f (salmon color arc of the outermost ring in Fig 2). The tree topology indicates that a major clade diversified from the prototypical group around the time of Gymnosperm divergence (blue arc of the outermost ring in Fig 2). This clade further diversified into the three immune regulator CBP60a, CBP60g, and SARD1 subfamilies (pink, cyan, and green arcs in the second outermost ring in Fig 2) around the time of Angiosperm divergence. Hence, we call this clade the immune-related clade. It is evident from the branch lengths that the immune regulator subfamilies have been evolving much faster than the prototypical group.

**Fig 2.**
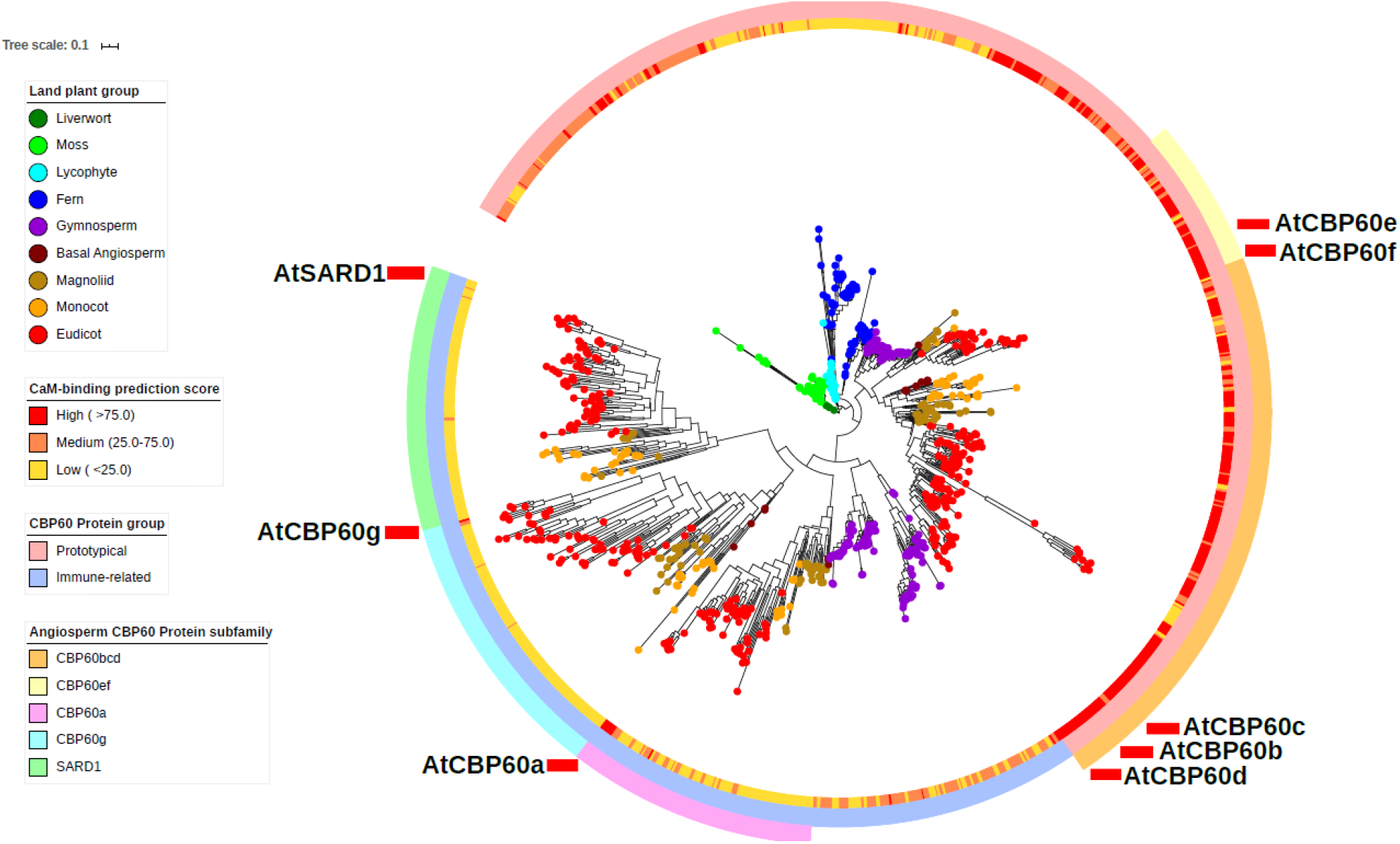
The immune-related clade diversified at or immediately before the Gymnosperm divergence and has been evolving very fast. The phylogenetic tree of 1024 CBP60 protein sequences from 247 land plant species was inferred using RaXML (Stamatakis 2014) based on ClustalW alignment (Thompson et al. 1994) and visualized using iTOL (Letunic and Bork 2019). Colored dots at the leaves of the tree represent the land plant group (from Liverwort to Angiosperm) the CBP60 member originated from according to the color code for the phylogenetic clade. The inner ring with the yellow (low) to red (high) color range shows the CaM-binding score predicted by CaMELS (Abbasi et al. 2017). The middle ring shows the CBP60 subfamilies according to AtCBP60 names. The outer ring shows the prototypical group (salmon) and the immune-related clade (blue). The AtCBP60 leaf positions are indicated in the outermost layer. The branch length scale of 0.1 (substitution per site) is shown at the top left.

None of the six Basal Angiosperm species had SARD1 subfamily members. This could be due to SARD1 deletion in Basal Angiosperms or to SARD1 subfamily diversification from the CBP60g subfamily after Basal Angiosperm divergence. A likelihood-ratio test of the tree topologies for these two possible cases favored the tree structure with SARD1 deletion in Basal Angiosperms (*p* < 0.01; Figs S2A and S2B). We identified multiple SARD1 subfamily members in recently-published genome sequences of a Basal Angiosperm water lily order (Zhang et al. 2020) (Fig S2C). This new information strongly supports our conclusion that diversification of all three immune regulator subfamilies occurred around the time of Angiosperm divergence.

### The common ancestor of the immune regulator subfamilies appears to have lost CaM-binding ability

AtCBP60a-f have CaM-binding domains (CaMBD) in their C-terminal regions (C-CaMBD), AtCBP60g has one in its N-terminal region (N-CaMBD), but AtSARD1 lacks CaM-binding ability (Reddy et al. 2002; Wang et al. 2009; Wang et al. 2011) (Fig 1). We investigated when the C-CaMBD domain was lost during evolution of the immune regulator subfamilies using CaMELS (Abbasi et al. 2017), which was the only algorithm among those we tested that correctly predicted C-CaMBD and N-CaMBD among AtCBP60 members. Generally, C-CaMBD was conserved among CBP60 proteins, except for some specific clades, including the immune regulator subfamilies (Fig 2, innermost ring). C-CaMBD in AtCBP60a and N-CaMBD in AtCBP60g appear to have been reacquired relatively recently. These observations suggest that CaM binding activity may not be essential for the CBP60 immune regulator functions, or alternatively, loss of C-CaMBD may have coincided with evolution of a CaM-binding adaptor protein that works together with the immune regulators. The latter could explain reacquisition of CaMBD in some of the immune regulators.

There were two forms of C-CaMBD loss (Fig 3). One form is deletion of the C-terminal region corresponding to C-CaMBD, such as in the SARD1 subfamily members (Fig 3A). The other form is mutations in C-CaMBD with weakly to moderately alignable sequences remaining in the C-terminal region. (Wang et al. 2009). A secondary structural property of amphipathic helix may be important for CaMBDs (Degrado et al. 1987). When the CaM-binding prediction scores at the C-CaMBD region were compared to the α-helix length and the hydrophobic moment (as an indication of a potential amphipathic α-helix), a long α-helix appeared important while its amphipathicity did not (Fig 3B). Although we cannot exclude a possible high false negative rate in CaMELS predictions, mutations that led to loss of CaMBD might have disrupted the CaMBDs through disruption of alpha-helices.

**Fig 3.**
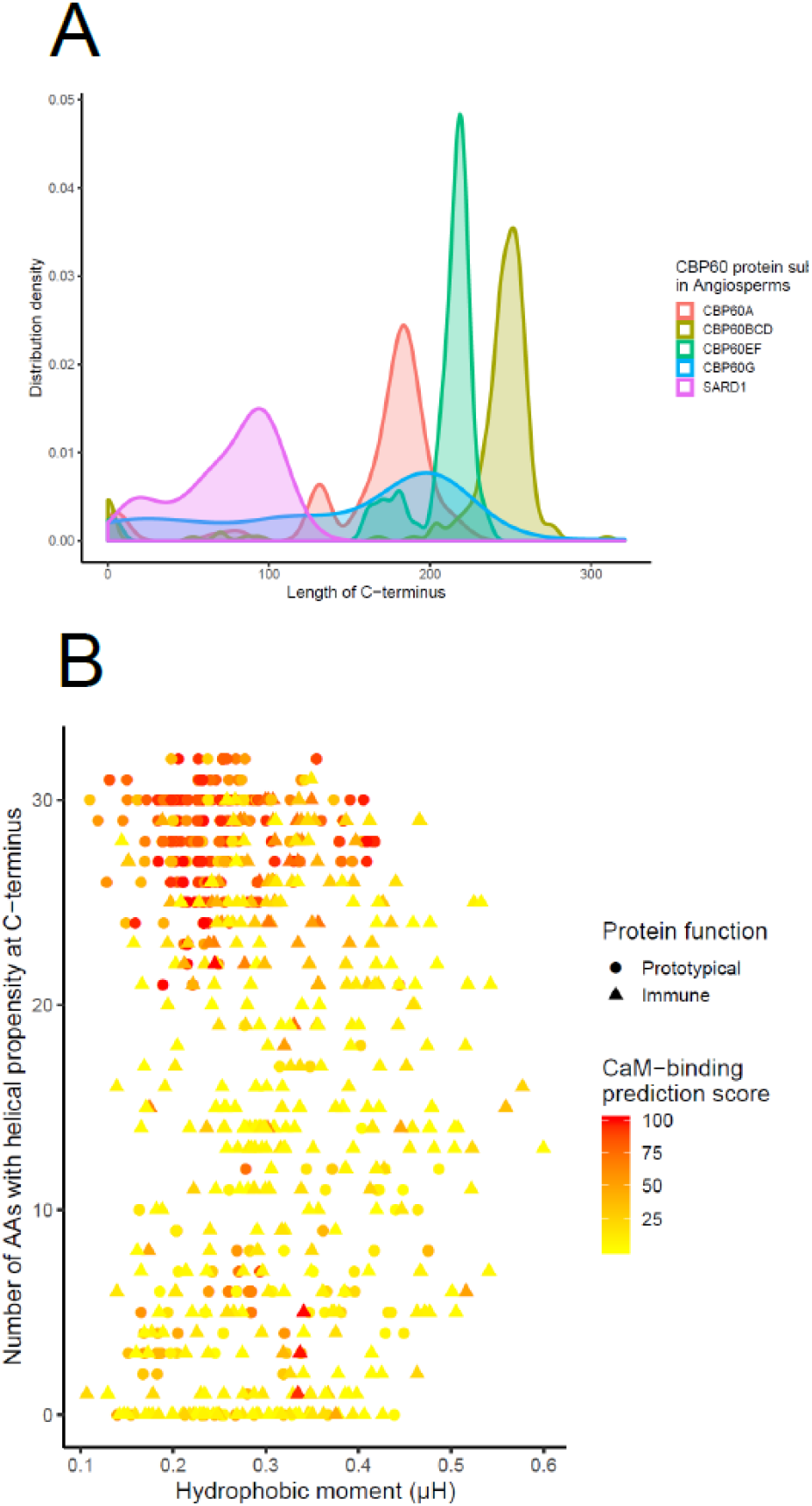
Loss of calmodulin-binding ability is caused by loss of the C-CaMBD region or mutations in C-CaMBD. (A) The distribution of the length of the region C-terminal to the CBP60-conserved domain in each subfamily in Angiosperms. (B) Contributions of the length of alpha-helix and its hydrophobic moment to the CaM-binding prediction score according to the CaMELS algorithm (Abbasi et al. 2017).

### Fast evolution of the CBP60 immune regulator subfamilies among Core Eudicots was caused by both high proportions of polymorphic sites and high evolution rates per site

The branch lengths of the CBP60 protein phylogenetic tree (Fig 2) strongly suggested fast evolution of the immune regulator subfamilies, especially the CBP60g subfamily. We closely investigated their fast evolution among Core Eudicot species. We chose Core Eudicots because genome sequences (vs. transcriptome sequences) of many diverse Core Eudicot species were available. To reduce the influence of conservation due to recently shared ancestry, a single species from each of 12 taxonomic orders was chosen for the analysis (Figs 4A and S3). To simplify the analysis, when one species has more than one member per subfamily, only one member was selected. This subfamily member selection should not introduce strong bias: in all such cases, the members in a single subfamily were paralogs diversified after divergence of the corresponding taxonomic orders. For example, AtCBP60b, AtCBP60c, and AtCBP60d are paralogs in this time scale, and one of them was selected as the Arabidopsis sequence for the CBP60bcd subfamily. We also conducted the subsequent analyses with an alternative set of selected CBP60 members and obtained similar observations (indicated as “the alternative set” for the corresponding supplemental figures), confirming the absence of strong bias due to CBP60 member selection.

We compared the CBP60 subfamily members to benchmark protein sequences in the 12 Core Eudicot species. The benchmark protein sequences were selected for high quality multiple sequence alignments from protein sequencies encoded by a set of 353 “single-copy” genes for Angiosperm phylogeny studies (Johnson et al. 2019). Single-copy genes suggest that the function of each gene has been conserved during Angiosperm evolution. The selected benchmark protein sequence alignment across the 12 species had 70,022 amino acid sites, including 39,561 polymorphic sites, which we consider as representing the “genomic norm”. Fifty thousand random samples with replication and with the same amino acid site number as that of the test protein alignment were made from the benchmark alignment for the benchmark distribution estimate, from which the median and the 95% and 99% confidence intervals were calculated for the measure of interest. We judge an observed value to be significantly different when it is outside the 99% confidence interval.

Outcomes of fast evolution could be detected as an increased proportion of polymorphic sites, an increased evolution rate per polymorphic site, or both. First, we examined the proportion of polymorphic sites among the amino acid sites of the high-quality alignments between the CBP60 subfamilies and the benchmark proteins. The proportions of polymorphic sites were significantly higher with the immune regulator subfamilies, especially with the CBP60g subfamily, while no significant difference was observed with the CBP60bcd subfamily (Fig 4B; Fig S4A for the alternative set).

**Fig 4.**
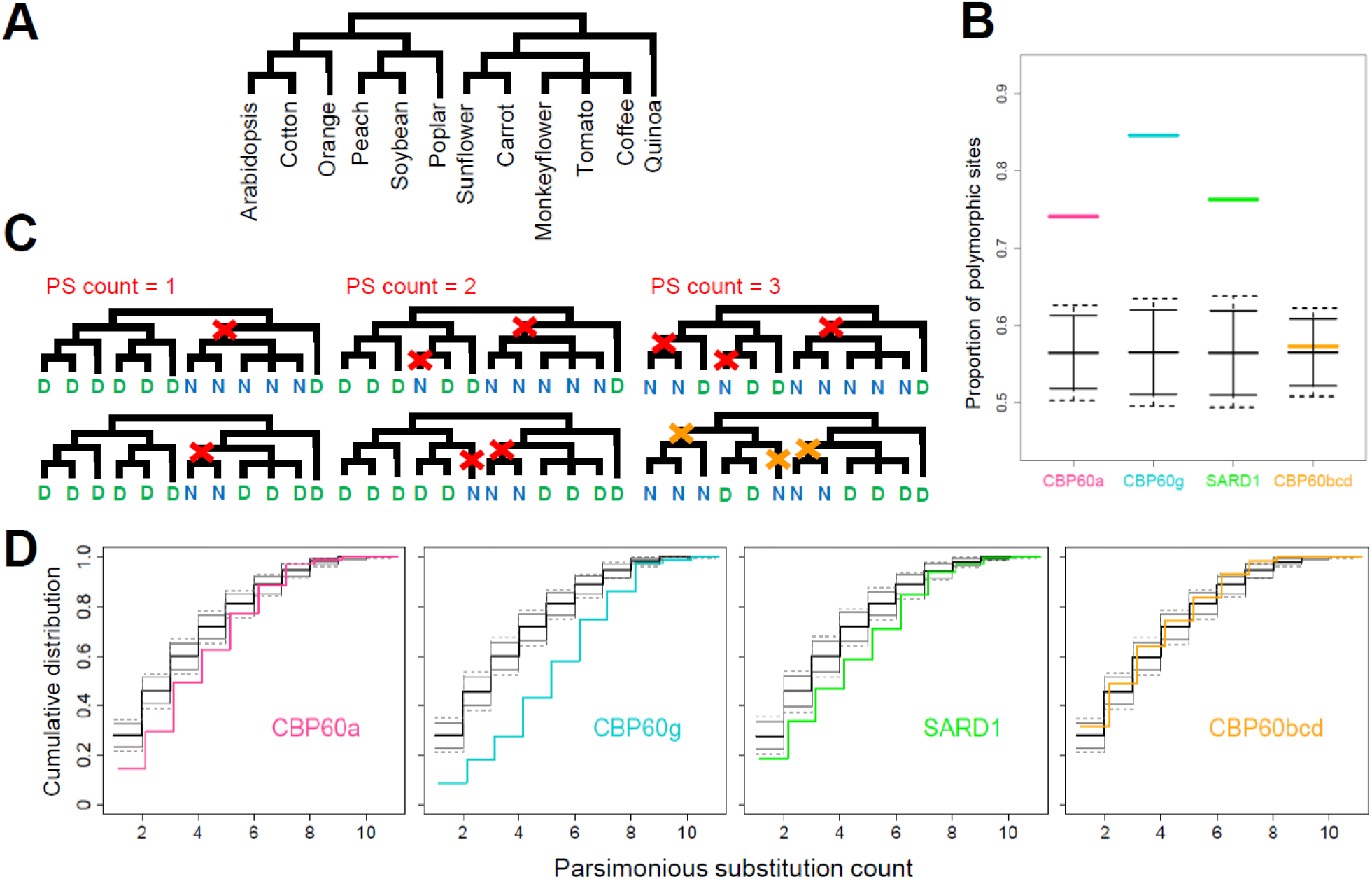
The immune regulator subfamilies have been evolving very fast among 12 diverse Core Eudicot species. (A) The phylogeny of the 12 Core Eudicot species used. (B) The proportions of the polymorphic sites of the CBP60 subfamilies are compared to the polymorphic site proportion distribution of the samples of the same sizes from the benchmark proteins. (C) Examples of the parsimonious substitution count. Two amino acid site examples with two variants, D and N, for parsimonious substitution count = 1, 2, or 3 are shown. While a red “X” in a tree shows the particular substitution event for the example amino acid variation, an orange “X” shows a possible substitution event as multiple sets of substitution event positions are possible in the latter case. (D) The cumulative distributions of the parsimonious substitution count of the CBP60 subfamilies are compared to those of samples from the benchmark proteins. Significant right shifts of the cumulative distributions for the immune regulator subfamilies indicate significantly higher parsimonious substitution counts than the benchmark proteins. The CBP60 subfamilies are color-coded in B and D: salmon, CBP60a; cyan, CBP60g; green, SARD1; orange, CBP60bcd. Horizontal thick solid line, median; thin solid line, 95% confidence interval, thin dashed line, 99% confidence interval in B and D. In D, the cumulative distributions for the CBP60 subfamilies are slightly positively offset for better visualization.

Second, we examined the evolution rate per polymorphic site using the parsimonious substitution counts per amino acid site among the 12 Core Eudicot species given their species phylogeny (examples of the parsimonious substitution count in Fig 4C). The protein sequences from 12 diverse species represent sparse data, and the actual substitution count is likely higher than the parsimonious count. Since we are interested in whether the parsimonious substitution count distribution of a CBP60 subfamily is shifted higher or lower compared to that of the benchmark, the cumulative distribution of the parsimonious substitution count was compared (Fig 4D; Fig S4B for the alternative set). Note that when the parsimonious substitution count distribution is shifted higher, the cumulative distribution curve is shifted toward the right in the plot. Significantly higher parsimonious substitution counts, which indicate significantly higher evolution rates per site, were observed with all immune regulator subfamilies, especially with the CBP60g subfamily. No significant difference was observed with the CBP60bcd subfamily. Thus, fast evolution of the CBP60 immune regulator subfamilies is evident in both higher proportions of polymorphic sties and higher evolution rates per polymorphic site, compared to the genomic norm.

Fast evolution could be caused by release from negative selection, including pseudogenization, as well as by strong and varying selection. Despite their very fast evolution relative to the CBP60bcd prototypical subfamily, pseudogenization is not the case for the CBP60 immune regulator subfamilies since their Arabidopsis members are all functional and since the vast majority of diverse Angiosperm species has maintained members of all three immune regulator subfamilies. Thus, their very fast evolution is likely a result of strong and varying selection, such as pressure from fast-evolving pathogen effectors targeting CBP60 immune regulator members. A fungal effector is known to target AtCBP60g (Qin et al. 2018). It is conceivable that multiple effectors target these major positive immune regulator subfamily members at a variety of sites.

### Protein Evolution Analysis in a Euclidean Space (PEAES)

Phylogenetic analysis of sequences using standard evolution models assumes stationary, reversible, and homogeneous evolution (Naser-Khdour et al. 2019). These conditions are clearly violated when evolution is under strong and varying selection. We postulated that when a site is under strong selection, the amino acid at that site is selected based mainly on physical-chemical characteristics of amino acids and is relatively independent of the lineage ancestry when the lineage divergence is sufficiently old. Divergence of the 12 Core Eudicot species lineage is likely sufficiently old for evolution of many amino acid sites in the CBP60 immune regulator subfamilies judged by their high parsimonious substitution counts. Thus, we decided to use a metric of physical-chemical characteristics of amino acids to evaluate relatedness of selection at each site. Venkatarajan and Braun described physical-chemical characteristics of 20 amino acids by five-dimensional Euclidean coordinates after applying linear dimensionality reduction to 237 physical-chemical properties (Venkatarajan and Braun 2001). We added one more dimension to include absence of an amino acid, i.e., a gap in an alignment (Table S3). This physical-chemical description of amino acids in a six-dimensional Euclidean space allows geometrical tracking of evolution at each site in a protein sequence alignment. For example, the pairwise Euclidean distance between amino acids can be defined as the amino acid dissimilarity measure (Table S4). We call this geometric analysis platform Protein Evolution Analysis in a Euclidean Space (PEAES).

In the subsequent section, we examine possible coevolutionary interactions among the CBP60 immune regulator subfamilies. We focused our analysis on the CBP60-conserved domain because it is impossible to obtain a reliable multiple sequence alignment across the subfamilies outside the conserved domain (Fig S5). The CBP60-conserved domain sequence that is highly conserved in the prototypical group contains 293 aa. To align the domain sequences from the immune regulator subfamilies together, three gaps were added, resulting in the multiple sequence alignment consisting of 296 sites (Fig S6). Site positions within this 296-site alignment are used subsequently.

With PEAES, the measure of diversity at each site in the aligned sequences was defined as the mean of all pairwise distances among amino acids at that site. Fig 5A (Fig S7A for the alternative set) shows the conservation and diversity at each site in different CBP60 subfamilies. The CBP60bcd subfamily represents the prototypical group. It is evident that while many sites are highly conserved in the CBP60bcd subfamily, many sites in each immune regulator subfamily, particularly the CBP60g subfamily, are highly diverse. The mean of the diversity measure across the sites for the CBP60bcd subfamily is significantly lower than that for the benchmark proteins (the 99% confidence interval shown in the parentheses in Fig 5B), indicating that the CBP60-convserved domain is extremely well conserved within the CBP60bcd subfamily in the Core Eudicots. However, the means of the diversity measures across the sites for the immune regulator subfamilies are significantly higher than that for the benchmark, indicating that the immune regulator subfamilies are highly diverse compared to the genomic norm even within the domain conserved across all CBP60 subfamilies (Fig 5B; Fig S7B for the alternative set). This trend of highly diverse immune regulator subfamilies and highly conserved prototypical group is also evident using other measures, including the number of highly diverse sites (site diversity measure > 0.4415, which is the 90^th^ percentile value among the benchmark proteins) and the number of strictly conserved sites (Fig 5B; the number of highly diverse sites in the CBP60a subfamily is the only exception as it is not significant although it is higher than the benchmark mean value).

**Fig 5.**
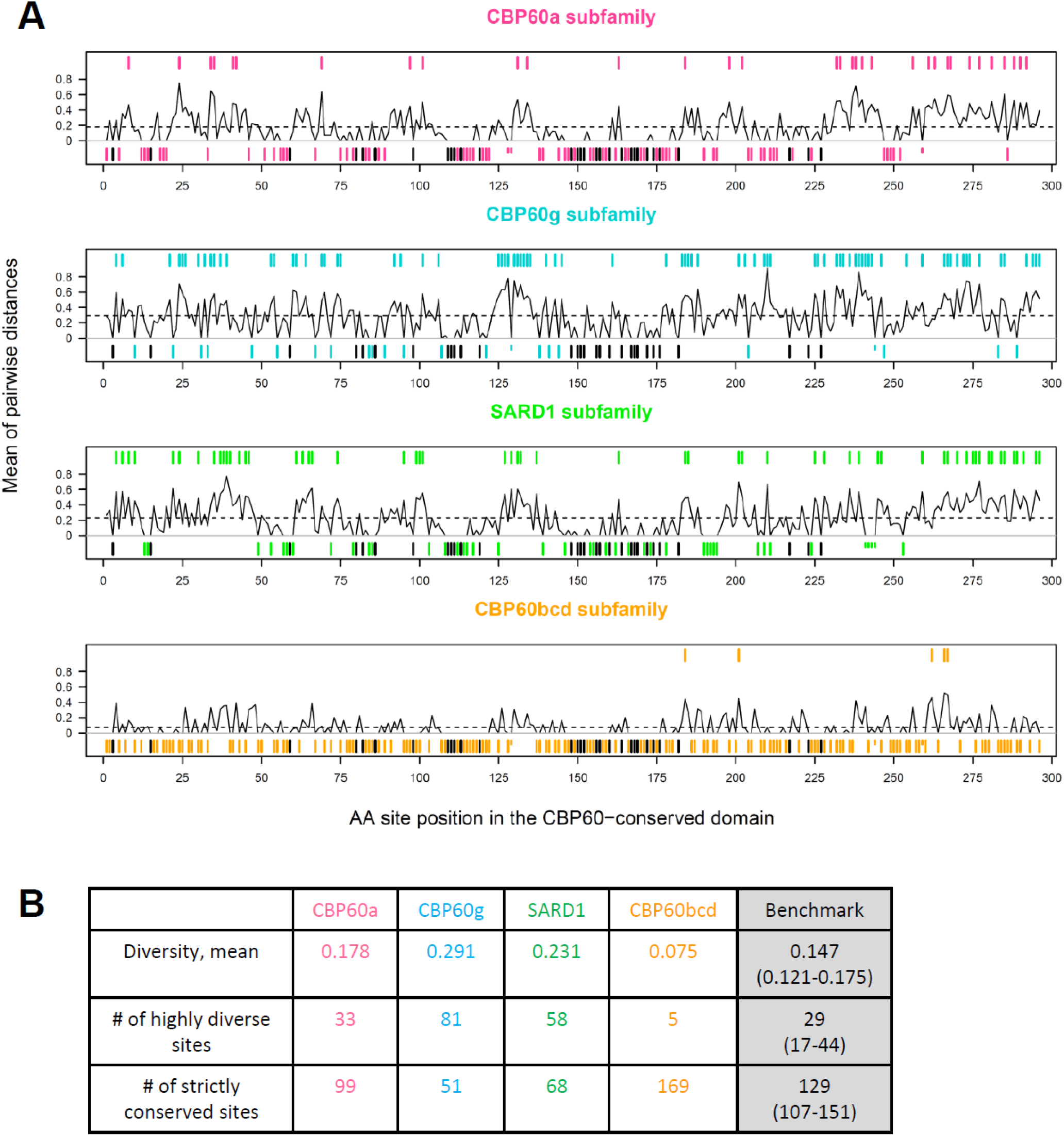
The CBP60-conserved domains of the immune regulator subfamilies are highly diverse among 12 Core Eudicot species. (A) The mean pairwise PEAES distance for each site across the species is shown as a line plot in each panel. One CBP60 member per subfamily was selected for each species. Strictly conserved sites are shown as segments at the bottom in each plot. The segments in black are the sites strictly conserved across all CBP60 members of the Core Eudicot species. The subfamily-specific sites are shown in salmon, cyan, green, and orange for the CBP60a, CBP60g, SARD1, and CBP60bcd subfamilies, respectively. Short segments represent subfamily-conserved gaps in the alignment. Highly diverse sites (pairwise distance mean > 0.4415) are shown at the top in each plot as segments colored according to the subfamily-color code. (B) Three measures of the diversity level, the mean of the PEAES diversity measure across the sites, the number of highly diverse sites, and the number of strictly conserved sites, for each subfamily and the benchmark protein are shown. For the benchmark (gray-shaded), the mean value and the 99% confidence interval in the parentheses are shown.

### Highly non-random, lineage-specific coevolutionary interactions across the immune regulator subfamilies are prevalent

As they share a common ancestor as recently as the time of Angiosperm divergence and they have overlapping or opposing functions in immune signaling, the immune regulators CBP60a, CBP60g, and SARD1 likely form a regulatory module in the immune signaling network. We investigated the possibility that coevolution of the regulatory module components confers resilience against negative impacts of pathogen effectors targeting the module components. There may be pressure for the partially functionally-redundant, positive immune regulators, CBP60g and SARD1, to be as dissimilar as possible at critical sites, which would reduce the probability that they are both targeted by a single pathogen effector (Hypothesis M1; “M” for mechanistic). On the other hand, if both the negative regulator CBP60a and one of the positive regulators are targeted by a single pathogen effector, simultaneous impairment of the positive and negative regulators would moderate the negative impact of the effector on immunity (Truman et al. 2013). If this is the case, the negative immune regulator CBP60a may be under pressure to remain similar to one or both of the positive regulators at critical sites (Hypothesis M2). Such selection imposed by pathogen effectors could be strong. In addition, the direction of selection and the sites under selection could change over short time periods within each plant species lineage. This is because pathogen effectors evolve very fast and multiple effectors from multiple pathogens may target the regulatory module of a single plant species. To detect coevolutionary interactions under rapidly changing selection, the analysis used must be specific to the site and the species lineage.

We defined the pairwise distance rank (PDR) of a single site as the rank of the pairwise PEAES distance between the members of two subfamilies of a particular species lineage among all permutations of the species lineages for the member of one of the two subfamilies (PEAES-PDR; Fig 6). To allow a species lineage-specific analysis, two hypotheses about the coevolutionary interactions among the three immune regulator subfamilies were derived from mechanistic hypotheses M1 and M2. If a site is not dissimilar between CBP60g and SARD1, CBP60a should have an amino acid similar to the amino acid in CBP60g or SARD1 to confer a protection effect (Hypothesis I1; “I” for interactions). On the other hand, at a site where CBP60a cannot provide protection through similarity to CBP60g or SARD1 (e.g., because it would impair CBP60a function), CBP60g and SARD1 should be dissimilar to avoid both proteins being targeted by a single effector (Hypothesis I2). To test hypotheses I1 and I2, the PDR between CBP60g (or SARD1) and CBP60a was plotted against the PDR between CBP60g and SARD1 for all sites of a single species. The plot was divided into four quadrants at the median rank of 12 on both axes (Fig 7A). Hypotheses I1 and I2 predict that the 1^st^ and 3^rd^ quadrants will be significantly enriched with sites. Enrichment in the 3^rd^ quadrant is consistent with Hypothesis I1, and enrichment in the 1^st^ quadrant is consistent with Hypothesis I2. On the other hand, if there are no coevolutionary interactions across the immune regulator subfamilies, the distributions of two PDRs should be independent, and no quadrant enrichment is expected. The significance of the site enrichment in the 1^st^ and 3^rd^ quadrants over the 2^nd^ and 4^th^ quadrants was tested by Fisher’s exact test (2-sided).

**Fig 6.**
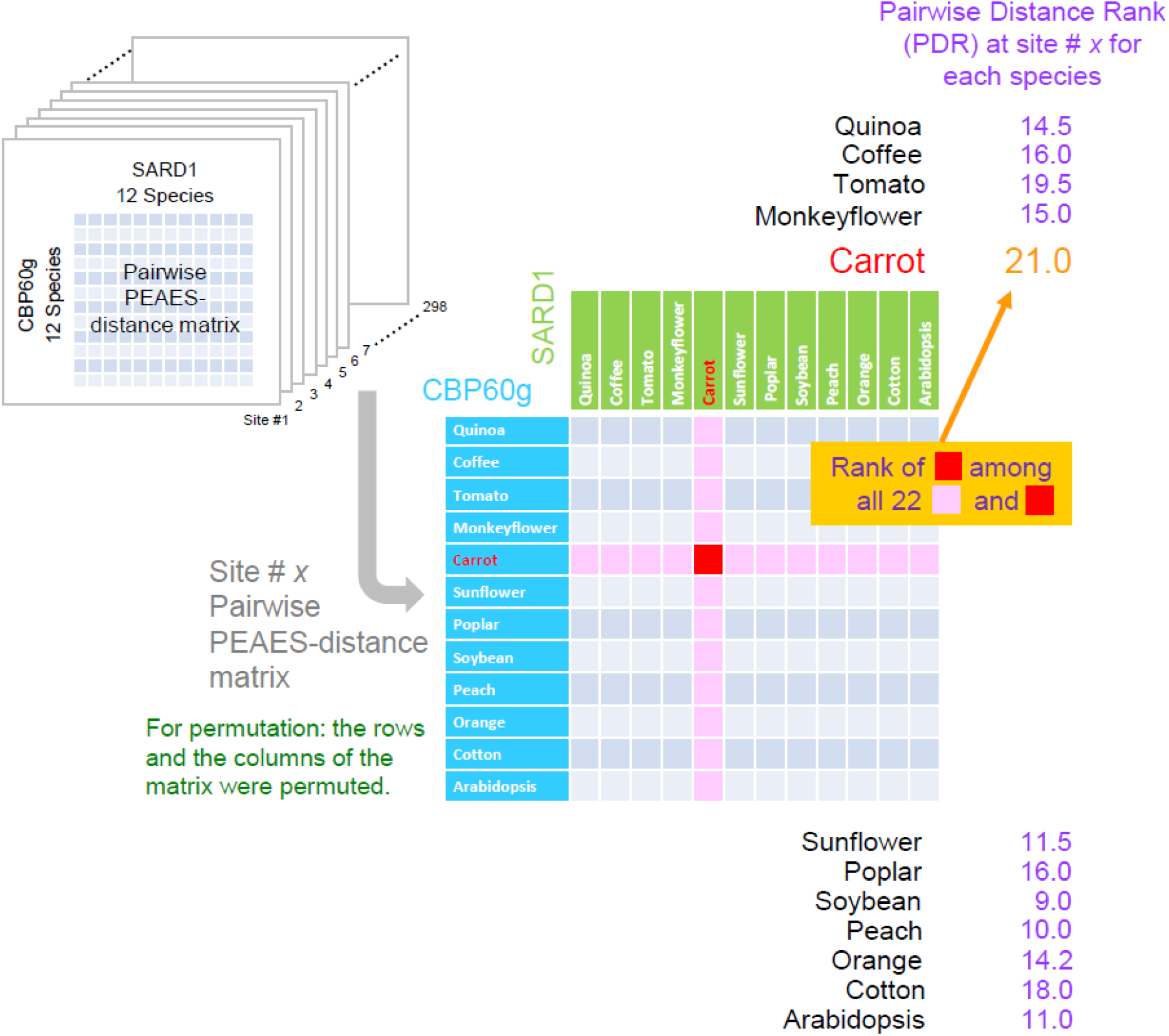
Pairwise distance ranks (PDRs) based on the PEAES distance metric. First, for each of the 296 amino acid sites in the multiple sequence alignment of the CBP60-conserved domain, a pairwise PEAES-distance matrix was made between members of two subfamilies (CBP60g and SARD1 in the figure as an example) in the 12 Core Eudicot species. Second, in the pairwise PEAES-distance matrix at a particular site (Site #*x*), for each species (Carrot in the figure as an example) the pairwise distance between the members of the two subfamilies of the species (red square in the matrix) was compared to the pairwise distance between the member of one subfamily of this species (Carrot) and the member of the other subfamily of a different species (pink squares). The rank of the within-species comparison was determined among these 23 pairwise distances. This rank (21.0 for Carrot) is called the pairwise distance rank (PDR) at each site in each species.

**Fig 7.**
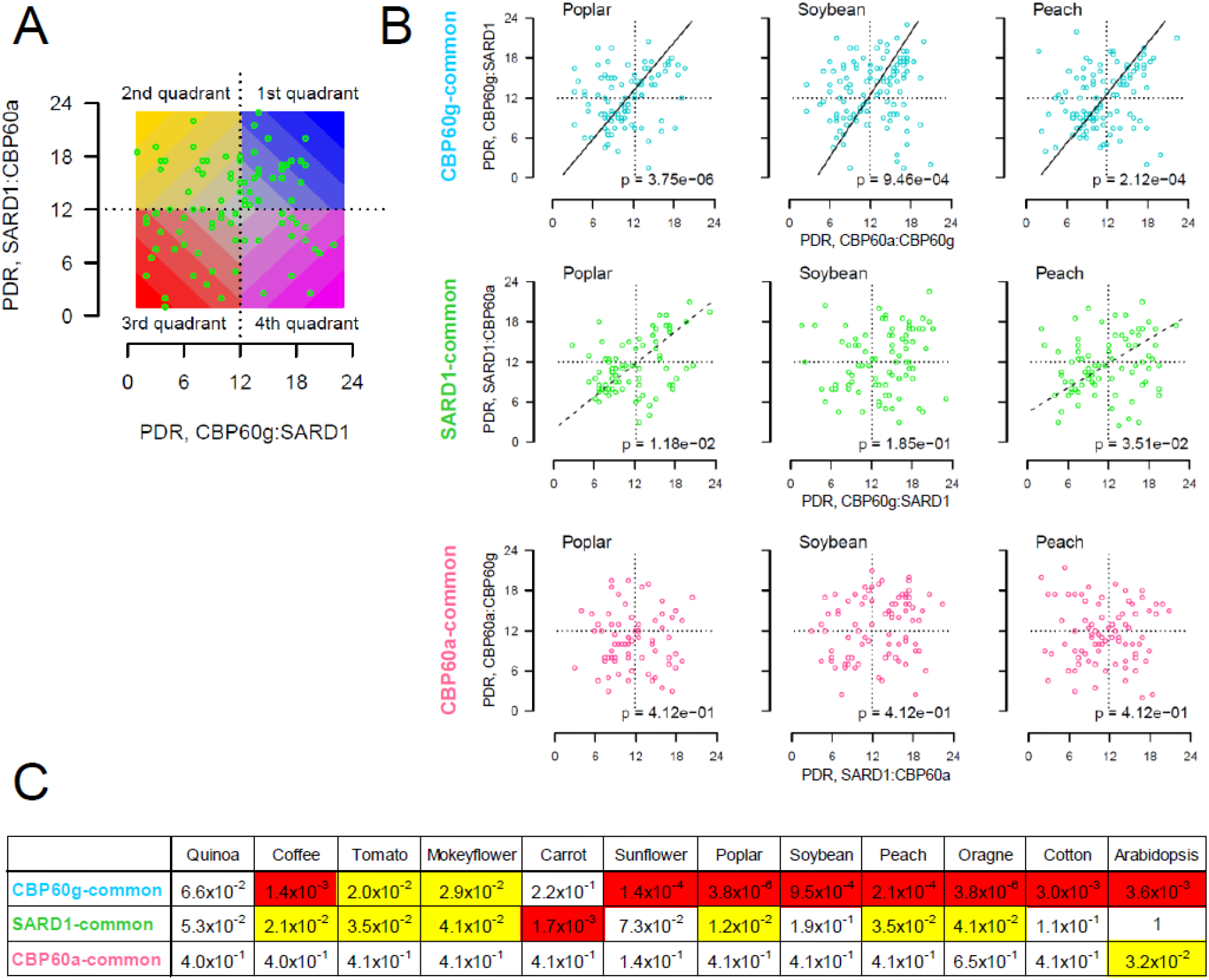
Non-random coevolutionary interactions that could protect the positive immune regulators are prevalent among the 12 Core Eudicot lineages. (A) Four quadrants in the SARD1:CBP60a PDR vs. CBP60g:SARD1 PDR plot (SARD1-common comparison) as an example. The color codes for the quadrants shown are used in Fig 8. (B) Plots for CBP60g:SARD1 PDR vs. CBP60a:CBP60g PDR (CBP60g common comparison; cyan color), SARD1:CBP60a PDR vs. CBP60g:SARD1 PDR (SARD1-common comparison; green color), and CBP60a:CBP60g PDR vs. SARD1:CBP60a PDR (CBP60a- common comparison; salmon color) are shown for three plant species, Poplar, Soybean, and Peach. Similar plots for all 12 Core Eudicot species are shown in Fig S8. The *p*-value (Fisher’s exact test, 2-sided, Benjamini-Hochberg FDR-corrected) is shown in the bottom right. When *p* < 0.01 or 0.01 ≤ *p* < 0.05, the PC1 (principal component 1) axis is shown as a black solid line or a dashed line, respectively, indicating significant enrichment of sites in the 1^st^ and 3^rd^ quadrants over the 2^nd^ and 4^th^ quadrants. (C) The *p*-values for CBP60g-common, SARD1-common, and CBP60a-common comparisons are shown for the 12 Core Eudicot species. Red background, *p* < 0.01; yellow background, 0.01 ≤ *p* < 0.05.

Significant enrichment of the sites in the 1^st^ and 3^rd^ quadrants (Benjamini-Hochberg FDR = 0.05) was observed in 10 out of 12 species lineages in the comparison of CBP60g:SARD1 PDR vs. CBP60a:CBP60g PDR (Fig 7B, 7C, and S8; CBP60g-common comparison) and in 7 out of 12 species lineages in the comparison of SARD1:CBP60a PDR vs. CBP60g:SARD1 PDR (SARD1-common comparison) (Fig S9 for the alternative set). In the union of the CBP60g- and SARD1-common comparisons, 11 out of 12 species lineages showed non-random coevolutionary interactions among the three immune regulator subfamilies, which is consistent with Hypotheses I1 and I2 (and consequently with Hypotheses M1 and M2). Even Quinoa, the only species that did not show significant enrichment, the FDR-corrected *p*-values were 0.066 and 0.053 for CBP60g- and SARD1-common comparisons, respectively (Fig 7C). Only one out of 12 species lineages shows a moderately significant enrichment in the 1^st^ and 3^rd^ quadrants in the comparison of CBP60a:CBP60g PDR vs. SARD1:CBP60a PDR (CBP60a-common comparison; Fig 7C), in which no enrichment was predicted by the hypotheses. The sites of CBP60g- and SARD1-common comparisons are shown along the CBP60-conserved domain in Fig 8 (Fig S10 for the alternative set), with the site-indicating bars color-coded according to the quadrants as in Fig 7A. The distribution of the colors of site bars does not show strong similarity between more closely related species, consistent with our assumption that the species lineage divergences were sufficiently old and that observed selections at the sites are largely consequences of species lineage-specific selection after lineage divergence. The notion of species lineage-specific selection is further supported by the fact that the values obtained by projecting the centered data onto *y* = *x* in each plot in Fig S8 show no obvious trend toward higher correlation between more closely related species pairs (Fig S11; Fig S12 for the alternative set). This projection onto *y* = *x* signifies the enrichment in the 1^st^ and 3^rd^ quadrants (Fig S11A). We conclude that many sites in the CBP60-conserved domain of the three immune regulator subfamilies have likely been coevolving under strong and varying pressure from pathogen effectors in a species lineage-specific manner, conferring a high level of resilience on the CBP60 immune regulatory module.

**Fig 8.**
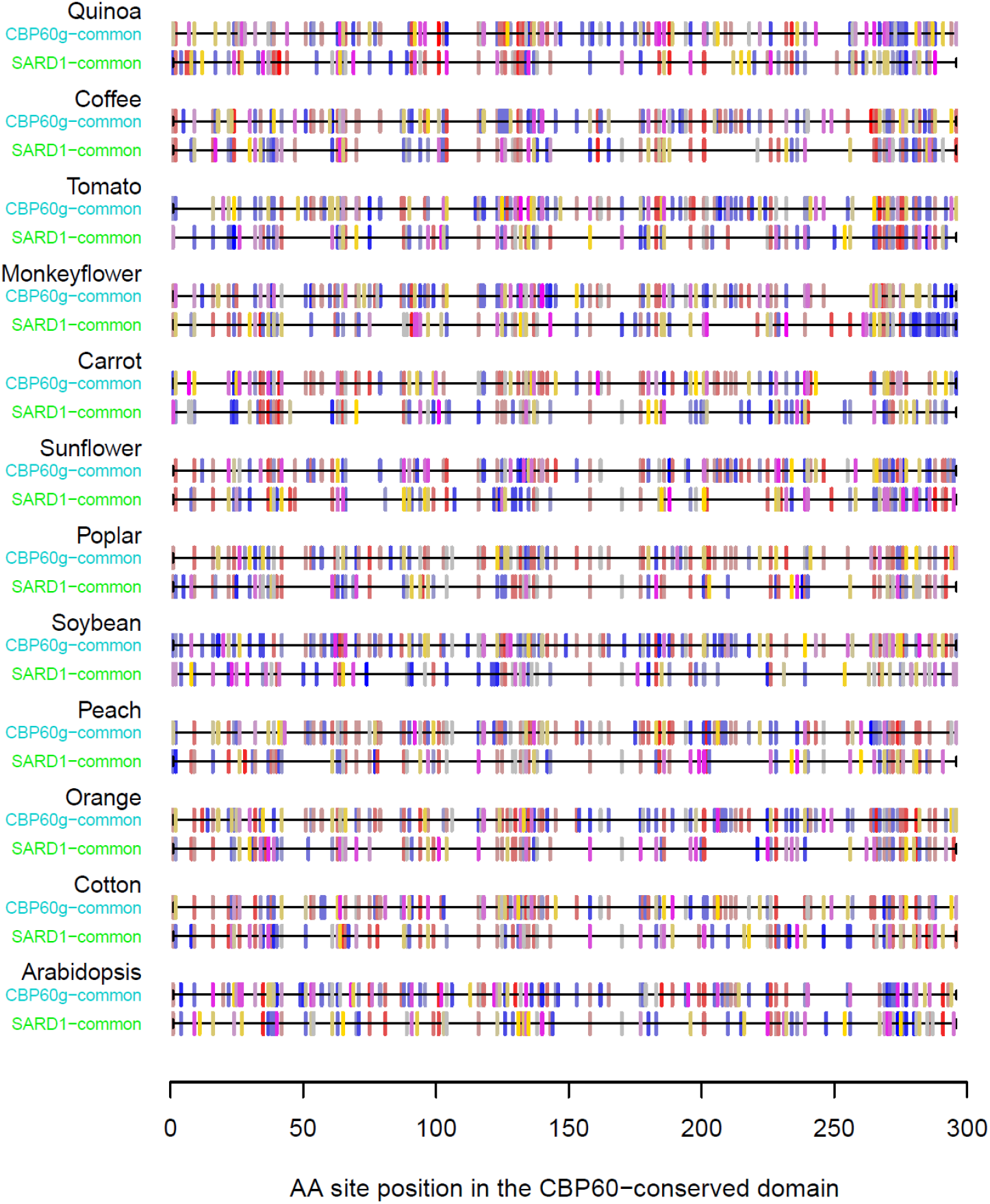
Site selections are largely specific to each of the Core Eudicot lineages. The sites in CBP60g- common and SARD1-common comparisons in each plant species (Figs S8A and S8B) are shown along the CBP60-conserved domain with the color-scaled quadrant information (Fig 7A).

## DISCUSSION

One type of protein family evolution analysis focuses on emergence of conserved domain structures (Domazet-Lošo and Tautz 2010). However, emergence of particular conserved domains may not provide information about evolution of particular biological processes and functions underlying them since some domains, such as DNA-binding domains, have versatile molecular functions that can be easily repurposed for different biological processes (e.g., (Gordân et al. 2011)). Since the CBP60-conserved domains of AtCBP60g and AtSARD1 have sequence-specific DNA-binding activity (Zhang et al. 2010; Qin et al. 2018), CBP60 proteins are likely DNA-binding transcription factors. A DNA-binding domain structure provides a backbone structure for physical interactions with DNA, and its binding specificity could vary in different proteins. It is possible that diversification of the immune-related clade from the prototypical group coincided with a substantial DNA-binding specificity change and consequently resulted in a large change in the gene set regulated by CBP60 transcription factors, i.e., immune-related CBP60 transcription factors have likely been neofunctionalized for a different biological process. Thus, tracking neofunctionalization of particular subfamilies important for biological processes of interest within a larger protein family defined by the conserved domain structure provides insights into evolution of biological processes.

AtCBP60g, AtSARD1, and AtCBP60a belong to the immune-related clade and regulate immune responses, including SA signaling. The fact that Non-Seed Land Plants, namely Liverworts, Mosses, Lycophytes, and Ferns, do not have a member of the immune-related clade strongly suggests that SA signaling and SA-dependent immunity are regulated differently between Angiosperms and Non-Seed Plants, or that SA-dependent immunity is absent in Non-Seed Plants. As SA is a major immune hormone in Angiosperms (Vlot et al. 2009), the organization of immune signaling networks could be very different between Angiosperms and Non-Seed Plants. However, this notion of different network organization does not exclude the possibility that Angiosperms and Non-Seed Plants share some elemental immune signaling machineries (de Vries et al. 2018). Instead, the notion implies that some properties of the immune signaling networks, such as network resilience, are very different between Angiosperms and Non-Seed Plants.

We discovered significant coevolutionary interactions at multiple sites in the CBP60- conserved domain across the CBP60a, CBP60g, and SARD1 immune regulator subfamilies in separate Core Eudicot species lineages, using the PEAES-PDR metric (Figs 6–8 and S8-S11). These observations indicate that although the three immune regulator subfamilies diversified around the time of Angiosperm divergence, the immune regulator subfamilies have been influencing fast evolution of one another in each plant species lineage. These coevolutionary interactions appear to have been maintained through the history of selection in the pathogen effector landscape that has been changing fast in a manner specific to plant species lineages. Thus, our discovery presents a new pattern of gene evolution after gene duplication: long lasting coevolution of duplicated (or multiplicated) genes while different species lineages have experienced different selection histories, resulting in lineage-specific selection outcomes for the coevolving gene pair.

The observed coevolutionary interactions are consistent with our hypotheses of coevolution mechanisms that promote functional resilience of the positive immune regulators, CBP60g and SARD1, against pathogen effectors. Hypothesis M1-Coevolution occurs via selection for amino acid dissimilarity between functionally overlapping, positive immune regulators to prevent simultaneous targeting by a single effector. Hypothesis M2 – Coevolution occurs via selection for amino acid similarity between functionally opposing, positive and negative immune regulators to assure simultaneous targeting by a single effector. There are other important immune signaling network components in Arabidopsis that contain multiple positive and negative regulator subfamilies within larger protein families, such as WRKY transcription factors (Birkenbihl et al. 2017). It will be interesting to apply the PEAES-PDR approach to these protein subfamilies to test whether their evolutionary patterns are also consistent with these hypotheses about coevolution mechanisms that lead to increased resilience of the regulatory modules against pathogen effectors.

The results of the PEAES-PDR analysis can also generate specific, mechanistic hypotheses at the molecular level. First, Qin et al. reported that AtCBP60g is targeted by the fungal effector VdSCP41 (Qin et al. 2018). Although AtSARD1 can coimmunoprecipitate VdSCP41 after overexpression in Arabidopsis protoplasts, this AtSARD1-VdSCP41 interaction appears to be much weaker than the AtCBP60g-VdSCP41 interaction (Figure 3A, Figure 4C, and Figure 3— figure supplement 2A in (Qin et al. 2018)). These observations may suggest that AtSARD1 mostly evades being targeted by VdSCP41. VdSCP41 binds a C-terminal region of AtCBP60g (corresponding to the region C-terminal to site 130 of the CBP60-conserved domain) (Qin et al. 2018). Our PEAES-PDR analysis detected multiple AtCBP60g sites C-terminal to site 130 potentially protected by dissimilarity between AtCBP60g and AtSARD1 (blue and magenta color bars on the line for Arabidopsis, CBP60g in Fig 8). If VdSCP41 or a very similar effector is a major driver for the dissimilarity between AtCBP60g and AtSARD1, substituting the amino acids at these sites in AtCBP60g to those at the sites in AtSARD1 may substantially weaken the interaction between AtCBP60g and VdSCP41.

Second, the sites indicated by yellow-colored bars in Fig 8 (i.e., sites in the 2^nd^ quadrant in Fig 7A) are those that are not protected either by dissimilarity between two positive regulators or by similarity between a positive regulator and the negative regulator. In some cases, these coevolutionarily unprotected sites coincide in both CBP60g- and SARD1-common comparisons in a single species. One possibility is that such sites may not have been targeted by any effectors in a recent lineage history, and hence there is no signature of protective selection. Another possibility is that they may be protected by some other molecular mechanisms, such as immunity mediated by specific resistance (R) proteins (Khan et al. 2016). A guard-type R protein detects some biochemical change caused by a pathogen effector in the cognate host protein and triggers immune responses. If there is protection by an R protein, keeping the amino acids at the coevolutionarily unprotected sites similar between CBP60 and SARD1 may be important for function of the R protein-based protection mechanism. For example, the Arabidopsis R protein SNC1 may guard AtCBP60g and AtSARD1 (Sun et al. 2018) and provide an opportunity to test this hypothesis. Furthermore, identification of proteins that interact with CBP60g and SARD1 whose interactions are sensitive to substitutions of the amino acids at the coevolutionarily unprotected sites to those at the same sites in other species may lead to discovery of such alternative protection mechanisms.

We demonstrated that coevolutionary interactions across the CBP60 immune regulator subfamilies of overlapping or opposing functions are consistent with the mechanistic hypotheses for increased resilience of this particular regulatory module in the immune signaling network. It is conceivable that this concept can be generalized among other immune signaling network components. Some pathogen effectors target multiple components of the immune signaling network (Toruño et al. 2016), suggesting existence of many other multiple-targeting effectors. At the same time, immune signaling network components are highly interconnected (Sato et al. 2010), and it is common for them to have overlapping or opposing effects on immunity. These two conditions, multiply targeting pathogen effectors and potential functional interactions among the effector targets, are exactly the conditions that appear to have been driving coevolution for increased resilience of the CBP60 immune regulator module. We developed a PEAES-PDR approach, which was powerful in detecting this type of coevolutionary interaction among alignable sequences and in generating specific and mechanistic hypotheses. Expanding coevolutionary interaction investigations to unalignable sequences of multiply targeted proteins will further advance our understanding of how the immune signaling network has been evolving to maintain or improve resilience in a strongly-selecting and rapidly-changing pathogenic landscape. This will be an exciting future direction in the study of biological network evolution.

## MATERIALS AND METHODS

### The CBP60 protein sequence set

A protein sequence database consisting of sequences from 271 species was constructed. The sequence data sources (Gonzales et al. 2005; Sato et al. 2008; Al-Dous et al. 2011; van Bakel et al. 2011; Sato et al. 2011; Bombarely et al. 2012; Goodstein et al. 2012; Ming et al. 2013; Singh et al. 2013; Kitashiba et al. 2014; Yagi et al. 2014; Proost et al. 2015; Unver et al. 2017; Li et al. 2018; Leebens-Mack et al. 2019; Zheng et al. 2019) are listed in Table S1. The database was searched by BLASTP (Camacho et al. 2009) using each of the Arabidopsis CBP60 members as a query and the union of the subject sequences that yielded bit scores higher than the Arabidopsis CBP60 that yielded the lowest bit score was identified as the CBP60 protein sequence set, consisting of 1432 sequences. Thus, the diversity of the CBP60 member set was controlled by the level of diversity among AtCBP60 members. We call this member identification procedure a Furthest Member Selection (FMS) approach. This FMS-based CBP60 protein sequence set did not include some distantly related CBP60 homologs (Text S1). We left them out of the set because they do not allow functional interpretations of CBP60 evolution since our knowledge of CBP60 functions is limited to the Arabidopsis members and their orthologs. The CBP60 protein sequence set was subjected to quality controls to remove possible products of assembly or annotation errors and to limit sequence representation to one species per genus for balanced overall representation. These quality controls resulted in the final set consisting of 1024 sequences from 247 species (Table S2). To ease identification of plant species, their common names or genus names are used.

### Phylogenetic analysis of CBP60 protein sequences

Generally, ClustalW (Thompson et al. 1994), a Maximum Likelihood (ML) method (Guindon and Gascuel 2003), and iTOL (Letunic and Bork 2019) were used for multiple sequence alignment, tree inference, and tree visualization, respectively. ML was also used for inference of most probable ancestor sequences at branch points in trees. MEGA7 (Kumar et al. 2016) was used for ClustalW and ML with the default parameters, except for 80% site coverage in ML (unless stated otherwise). We used the Most Recent Common Ancestor (MRCA)-Anchored, Clade-by-Clade Tree Inference (MACCTI) approach, in which the aligned sequence region used for tree inference is locally adjusted. Briefly, an overview tree for relationships among all major clades and MRCAs on the main CBP60 lineages was inferred using only the sequences with alignable C-terminal regions, the sequences of each clade and its flanking MRCAs on the main CBP60 lineages were used to infer the subtree for the clade, the MRCAs of the clades were used to infer the backbone tree showing the relationships among the clade MRCAs, and the clade MRCAs in the backbone tree were replaced with the corresponding clade trees to reconstruct the entire tree. Since species phylogeny, duplication, and deletion (SPDD rule) generally explained our tree of 1024 CBP60 sequences at high levels, we concluded that the tree was sufficiently accurate to support the analysis presented here.

### CaM-binding site prediction

CaMELS (Abbasi et al. 2017) was used for CaM-binding site prediction. The DECIPHER R Bioconductor package (Wright) was used to predict alpha-helical propensity. A Python script, hydrophobic_moment.py (https://gist.github.com/JoaoRodrigues/568c845915aea3efa3578babfd72423c) was used for calculation of the hydrophobic moment.

### Benchmark protein sequences and amino acid sites

Johnson et al. selected 353 genes that are single-copy in Angiosperm species for phylogenetic studies (Johnson et al. 2019). We selected 298 genes that had orthologs in Arabidopsis, rice, and the basal Angiosperm Amborella. The protein sequence database for the 12 Core Eudicot species we chose for the study was searched by BLASTP with default parameters (Camacho et al. 2009) using the Arabidopsis protein sequences for the 298 genes. We further selected 225 proteins that had High Scoring Pairs (HSP) in each of the 12 species. The HSP with the highest score was chosen if a single query resulted in multiple HSPs from a single species. The HSP subject protein sequences of the 12 species were subjected to multiple sequence alignment using ClustalW (Thompson et al. 1994) with default parameters in the msa package from Bio-Conductor project (Bodenhofer et al. 2015). High-quality aligned amino acid sites were selected using the following filters: (1) remove the sites that have gaps in more than three species; (2) remove the sites with more than two consecutive gaps in any of the sequences. The filtering resulted in 70,022 high-quality amino acid sites, including 39,561 polymorphic sites, in the multiple sequence alignment of the protein sequences from the 12 species. The high-quality amino acid sites for the 12 Core Eudicot species are given as a FASTA file in the supplement (“benchmark.aa.sites.12sp.fa”). The high-quality sites or polymorphic sites were sampled with replication with the appropriate sample size for a statistic of interest 50,000 times to estimate the benchmark sample distribution for the statistic. The median and the 95% and 99% confidence intervals of such distributions were used in Fig 4.

### Parsimonious substitution count algorithm

The principle of the parsimonious substitution count calculation is to parsimoniously infer a set of amino acids (potentially more than one amino acid) at each node of a tree with the topology of the species phylogenetic tree, and to add one count if a branch of the node has a non-overlapping set of amino acids. All nodes of the unrooted 12 Core Eudicot species phylogenetic tree, including the leaves, were numbered #1 to #21, starting with the leaves (Fig S13A). The nodes or leaves that directly connect to a particular node and have numbers lower than the node are the input nodes of the node. Nodes #13 and #21 have three input nodes while the other non-leaf nodes have two input nodes. The algorithm for a node is: (1) make a vector of amino acids by concatenating the amino acids from the input nodes; (2) from the amino acid vector, choose the amino acids with the maximum frequency as the inferred set of amino acids for the node (note that the maximum frequency could be 1, and in such a case, all input node amino acids are chosen); (3) add the number of the input nodes minus the maximum frequency value to the count. The final count after repeating the algorithm at each non-leaf node in the order of the node number is the parsimonious substitution count of the site. Although the inferred set of amino acids for a particular node may not cover all the possible cases, it does not affect the final parsimonious substitution count (e.g., Fig S13B). An example R script to calculate the parsimonious substitution count is given in a supplemental file (“parsimonious.substitution.count.r”).

### PEAES-PDR analysis

PEAES is a way to describe the physical-chemical characteristics of 20 amino acids and a gap in a 6-dimensional Euclidean space. A gap dimension was added to the 5-dimensional descriptions of physical-chemical characteristics of 20 proteinogenic amino acids by (Venkatarajan and Braun 2001) (Table S3). Thus, the dissimilarity between two amino acids (or between an amino acid and a gap) is defined by the Euclidean distance between them in the 6-dimensional space (Table S4). Twelve Eudicot species with high quality genome sequences were selected to represent 12 diverse taxonomic orders (Fig S3). One sequence for each of the CBP60a, CBP60bcd, CBP60g, and SARD1 subfamilies was selected for each species (48 sequences). Since some species had more than one paralog, two sets of 48 sequences were made by selecting different paralogs from these species. Each of these two sets was analyzed separately. The CBP60-conserved domains of 48 sequences were aligned by ClustalW (Thompson et al. 1994), and then the multiple sequence alignment was manually edited to remove insertional polymorphisms represented by only one or two sequences and to make the alignments consistent between the two sets. The alignments for the two sets are provided as supplemental files. To quantify the level of diversity at each site, the mean pairwise PEAES distance was calculated for each site for each subfamily (Fig 5). PDR was calculated at each site of the CBP60-conserved domain multiple sequence alignment between two CBP60 subfamilies for each of the 12 Eudicot species based on the PEAES metric according to the procedure depicted in Fig 6. If there are no species lineage-specific coevolutionary interactions between two CBP60 subfamilies, the mean expectation for PDR is the median rank of 12 among 23 pairwise PEAES distances. Thus, when PDR values for the corresponding sites in two subfamily pairs are plotted against each other, the random expectation is that the sites would distribute equally among the four quadrants of the plot divided at the median ranks of 12 (Fig 7A). Hypotheses I1 and I2 predicted enrichment of the sites in quadrants 1 and 3. The *p*-value associated with the odds ratio for this enrichment was calculated by Fisher’s exact test for each species for particular two-subfamily pairs. The *p*-value was corrected for multiple tests by Benjamini-Hochberg’s False Discovery Rate across the species in each combination of two-subfamily pairs.

Detailed methods and discoveries made based on the phylogenetic relationships but not described in the main text are in Text S2.

## AUTHOR CONTRIBUTIONS

F.K. conceived the project. Q.Z. assembled the protein sequence database. Q.Z., K.M., and F.K. analyzed the data. K.M. and F.K. wrote the manuscript.

## ACKNOWLEDGEMENTS

We thank Gane Ka-Shu Wong and Eric Carpenter for allowing us access to the 1000 Plants transcriptomic sequence database and Fay-Wei Li for allowing us access to the Waterfern and Floating fern genome sequences, prior to their publications. We also thank Yaniv Brandvain, Ya Yang, and Emma Goldberg for technical advice and discussion and Jane Glazebrook for editing of the manuscript. Q. Z. was supported by a scholarship from China Scholarship Council and funds from the Zhiming Zhang Laboratory. This work was supported by National Science Foundation (grant numbers MCB-1518058 and IOS-1645460 to F. K.).

## SUPPLEMENTAL INFORMATION

Text S1. Examples of CBP60 homologs that were not included in this study

Text S2. Detailed methods and the features of the CBP60 protein sequence phylogeny that are not discussed in the main text.

Fig S1. A pdf version of the CBP60 phylogenetic tree in Fig 2. This includes the name of each CBP60 member, containing the species name and the sequence ID in the originating database.

Fig S2. The three CBP60 immune regulator subfamilies diversified at or immediately before the emergence of Angiosperms. Two tree topology models, diversification of the CBP60g and SARD1 subfamilies at or before divergence of Basal Angiosperms (A) or after (B), were tested with CBP60 members from Basal Angiosperms, Magnoliids, and Chloranthales. Tree structure (A) was favored by the SH-test (p < 0.01) (Stamatakis 2014). Liverwort 0018s0017 was included as an outgroup. (C) SARD1 sequences were identified in recently published waterlily genome sequences (Zhang et al. 2020). The CBP60 sequence set used in (A) and (B) was supplemented with CBP60 sequences from *Nuphar advena* and *Nymphaea colorata* for the sequence phylogenetic tree. The *N. advena* and *N. colorata* SARD1 sequences are emphasized by a green dotted circle.

Fig S3. Each of 12 selected Core Eudicot species represents a different taxonomic order. The phylogeny of the Core Eudicot taxonomic orders and the 12 selected species representing diverse taxonomic orders are shown. These 12 Core Eudicot species and their lineages were used subsequently.

Fig S4. The analyses shown in Fig 4B and 4D were conducted with the alternative set. The alternative set of CBP60 members was made by selecting the member with the longest branch length, i.e., the most diverse member, among the paralogs of the species when the paralogs exist. This selection scheme was opposite from the set used in the main text. Therefore, all the CBP60 members in the alternative set are different from the ones in the set used in the main text for species with multiple members per subfamily. (A) The proportions of the polymorphic sites of the CBP60 subfamilies are compared to the polymorphic site proportion distribution of the samples of the same sizes from the benchmark proteins. (B) The cumulative distributions of the parsimonious substitution count of the CBP60 subfamilies are compared to those of samples from the benchmark proteins. Significant right shifts of the cumulative distributions for the immune regulator subfamilies indicate significantly higher parsimonious substitution counts than the benchmark proteins. The CBP60 subfamilies are color-coded: salmon, CBP60a; cyan, CBP60g; green, SARD1; orange, CBP60bcd. Horizontal thick solid line, median; thin solid line, 95% confidence interval, thin dashed line, 99% confidence interval in A and B. In B, the cumulative distributions for the CBP60 subfamilies are slightly positively offset for better visualization.

Fig S5. Difficulty in alignment of the immune regulator subfamilies outside the CBP60- conserved domain. Multiple sequence alignment of the regions N-terminal (A) and C-terminal (B) to the CBP60-conserved domains of CBP60a, CBP60g, SARD1, and CBP60bcd subfamily members of the 12 Core Eudicot species. The sequences from the top are 12 sequences each of the CBP60a, CBP60g, SARD1, and CBP60bcd subfamilies. Within each subfamily, the species order is the same as in Fig S3. The figures were made from screenshots of MEGA 7 alignments. The alignment was made by ClustalW in MEGA7 with the default parameter set values. No manual edits were made.

Fig S6. Multiple sequence alignment of the CBP60-conserved domains from CBP60a, CBP60g, SARD1, and CBP60bcd subfamily members of the 12 Core Eudicot species. One member per subfamily per species was selected (48 sequences). The alignment was divided into (A) sites 1-153 and (B) 154-296. The sequences from the top are 12 sequences each of the CBP60a, CBP60g, SARD1, and CBP60bcd subfamilies. Within each subfamily, the species order is the same as in Fig S3. The figures were made from screenshots of MEGA 7 alignments. The alignment FASTA file is provided as the supplemental file “CBP60.4subfam.12species.fas”. To make the alignment compact, the alignment was manually edited with the following amino acids deleted. The amino acid position number is counted from the first amino acid of the CBP60- conserved domain of each sequence. SARD1_Quinoa_62033803, #N261, #G210-#K214, #Y99; CBP60g_Coffee_02g34690, #K206; CBP60bcd_Sunflower_5g00005047, #N205-#K210; SARD1_Tomato_03g119250.2.1, #I219, #T66; SARD1_Coffee_04g06970, #A131-#A132; SARD1_Peach_5G223600, #A137, #S48-#D49, #N39-#Q42; CBP60g_Arabidopsis_AT5G26920.1, #T100; SARD1_Poplar_015G045300.1, #C91; CBP60g_Monkeyflower_B00990, #Q63; SARD1_Sunflower_16g00022942, #V60; SARD1_Soybean_09G182400, #D65; SARD1_Orange_1g011961m, #I40

Fig S7. (A) The analysis shown in Fig 5A was conducted with the alternative set. The alignment FASTA file is provided as the supplemental file “CBP60.4subfam.12species.set2.fas”. To make the alignment compact, the alignment was manually edited with the following amino acids deleted. CBP60g_Coffee_02g34690, #K206; SARD1_Quinoa_62029901, #N258, #E210-#K211, #C99; CBP60bcd_Sunflower_5g00005047, #N205-#K210; CBP60g_Soybean_03G232400, #L161-#K162; SARD1_Tomato_12g036390.1.1, #I127, #Y26; SARD1_Monkeyflower_I00405, #E130; SARD1_Coffee_04g06970, #A131-A132; SARD1_Peach_5G223600, #A137-#Y138, #I46-#A47, #N39-#Q42; SARD1_Arabidopsis_AT1G73805.1, #G126; CBP60g_Arabidopsis_AT5G26920.1, #T100; SARD1_Poplar_012G054900.1, #C91; SARD1_Soybean_08G044400, #Y71-#S72, #A63-#P64, #V40-#A41; SARD1_Orange_1g011961m, #I40. The amino acid position number is counted from the first amino acid of the CBP60-conserved domain of each sequence. (B) A table corresponding to Fig 5B with the alternative set.

Fig S8. Non-random coevolutionary interactions that could protect the positive immune regulators are prevalent among the 12 Core Eudicot lineages. (A) CBP60g:SARD1 PDR vs. CBP60a:CBP60g PDR (CBP60g common comparison) plot for each species. (B) SARD1:CBP60a PDR vs. CBP60g:SARD1 PDR (SARD1-common comparison) plot for each species. (C) CBP60a:CBP60g PDR vs. SARD1:CBP60a PDR (CBP60a-common comparison) plot for each species. In each plot, the *p*-value (Fisher’s exact test, 2-sided, Benjamini-Hochberg FDR-corrected) is shown in the bottom right. When *p* < 0.01 or 0.01 ≤ *p* < 0.05, the PC1 (principal component 1) axis is shown as a black solid line or a dashed line, respectively, indicating significant enrichment of sites in the 1^st^ and 3^rd^ quadrants over the 2^nd^ and 4^th^ quadrants. Significant enrichment in the 1^st^ and 3^rd^ quadrants (BH-FDR = 0.05) was observed with 10 out of 12 lineages for the CBP60g-common comparison and with 7 out of 12 lineages for the SARD1-common comparison (the union of CBP60g- and SARD1-common comparisons with significant enrichment are 11 out of 12), while only one lineage with moderately significant enrichment out of 12 lineages was observed for the CBP60a-common comparison.

Fig S9. The analysis shown in Fig S8 was conducted with the alternative set. Significant enrichment in the 1^st^ and 3^rd^ quadrants (BH-FDR = 0.05) was observed with 8 out of 12 lineages for the CBP60g-common comparison and with 6 out of 12 lineages for the SARD1-common comparison (the union of CBP60g- and SARD1-common comparisons with significant enrichment are 9 out of 12), while only one lineage with moderately significant enrichment out of 12 lineages was observed for the CBP60a-common comparison.

Fig S10. The analysis shown in Fig 8 was conducted with the alternative set.

Fig S11. Coevolutionary selections observed in the immune regulator subfamilies are largely lineage specific. (A) The site values projected onto *y* = *x* (red line) and centered at (*x*, *y*) = (12,12) as a measure of enrichment in the 1^st^ and 3^rd^ quadrants in a plot. Two example data points (blue) and their projected values as arrows (their sizes; green for negative and orange for positive values) are shown. (B-D) The projected values in each plot in Fig S8 were plotted between all pairs of 12 Core Eudicot species (B-D, respectively). There is no evident tendency that more closely related species have higher correlations in selected sites, which indicates that selections at the sites largely occurred after divergence of the 12 species lineages.

Fig S12. The analysis shown in Figs S11B-S11D was conducted with the projected values from each plot in Fig S9 for the alternative set.

Fig S13. The parsimonious substitution count calculation. (A) The node numbering in the 12 Core Eudicot species phylogeny. See Fig 4A for the actual species for the tree. (B) An example case in which the parsimonious substitution count algorithm inference of the node amino acids (in blue letters) does not cover all the possible cases, such as “a potential alternative” (in green letters). However, this type of potential discrepancy does not affect the parsimonious substitution count, which is two (i.e., the number of red “Xs” per tree).

Table S1. The list of 271 land plant species used in the study, including the nomenclature used in the text (“Database name”).

Table S2. The list of 1024 QC-ed CBP60 sequences (“Final set 1024” tab) and the list of 409 QC-omitted sequences (“QC fail” tab).

Table S3. The six-dimensional coordinates for amino acids in PEAES. The columns, E1-E6, represent the six dimensions in the Euclidean space.

Table S4. The pairwise Euclidean distance between amino acids in PEAES.

### Supplemental files

“CBP60.final.tree.nwk”: The Newick format tree file for the CBP60 sequence phylogenetic tree in Figs 2 and S1.

“CBP60.4subfam.12species.fas”: The FASTA multiple sequence alignment file for Fig S6.

“CBP60.4subfam.12species.set2.fas”: The FASTA multiple sequence alignment file for Fig S7 (“the alternative set”).

“peaes.pdr.figures.r”: an R script, which includes an algorithm for PEAES-PDR analysis of “CBP60.4subfam.12species.fas” and generates Figs 5, 8, and S8.

“aa.dist.mat.RData”: R object that is an equivalent of Table S4; required in “peaes.pdr.figures.r”

“benchmark.aa.sites.12sp.fa”: a FASTA file of the aligned high-quality amino acid sites for 12 Core Eudicot species (70,220 sites).

“parsimonious.substitution.count.r”: an R script, which is an example script to calculate the parsimonious substitution count for each CBP60 subfamilies from “CBP60.4subfam.12species.fas” above.

## REFERENCES

Abbasi WA, Asif A, Andleeb S, Minhas F ul AA. 2017. CaMELS: In silico prediction of calmodulin binding proteins and their binding sites. Proteins Struct. Funct. Bioinforma. [Internet] 85:1724–1740. Available from: https://pubmed.ncbi.nlm.nih.gov/28598584/

Al-Dous EK, George B, Al-Mahmoud ME, Al-Jaber MY, Wang H, Salameh YM, Al-Azwani EK, Chaluvadi S, Pontaroli AC, Debarry J, et al. 2011. De novo genome sequencing and comparative genomics of date palm (Phoenix dactylifera). Nat. Biotechnol. [Internet] 29:521–527. Available from: https://pubmed.ncbi.nlm.nih.gov/21623354/

van Bakel H, Stout JM, Cote AG, Tallon CM, Sharpe AG, Hughes TR, Page JE. 2011. The draft genome and transcriptome of Cannabis sativa. Genome Biol. [Internet] 12. Available from: https://pubmed.ncbi.nlm.nih.gov/22014239/

Birkenbihl RP, Liu S, Somssich IE. 2017. Transcriptional events defining plant immune responses. Curr. Opin. Plant Biol. [Internet] 38:1–9. Available from: https://pubmed.ncbi.nlm.nih.gov/28458046/

Bodenhofer U, Bonatesta E, Horejš-Kainrath C, Hochreiter S. 2015. Msa: An R package for multiple sequence alignment. Bioinformatics [Internet] 31:3997–3999. Available from: https://pubmed.ncbi.nlm.nih.gov/26315911/

Bombarely A, Rosli HG, Vrebalov J, Moffett P, Mueller LA, Martin GB. 2012. A draft genome sequence of Nicotiana benthamiana to enhance molecular plant-microbe biology research. Mol. Plant-Microbe Interact. [Internet] 25:1523–1530. Available from: https://pubmed.ncbi.nlm.nih.gov/22876960/

Camacho C, Coulouris G, Avagyan V, Ma N, Papadopoulos J, Bealer K, Madden TL. 2009. BLAST+: Architecture and applications. BMC Bioinformatics [Internet] 10. Available from: https://pubmed.ncbi.nlm.nih.gov/20003500/

Degrado WF, Erickson-Viitanen S, Wolfe HR, O’Neil KT. 1987. Predicted calmodulin-binding sequence in the γ subunit of phosphorylase b kinase. Proteins Struct. Funct. Bioinforma. [Internet] 2:20–33. Available from: https://pubmed.ncbi.nlm.nih.gov/3447166/

Domazet-Lošo T, Tautz D. 2010. A phylogenetically based transcriptome age index mirrors ontogenetic divergence patterns. Nature [Internet] 468:815–819. Available from: https://pubmed.ncbi.nlm.nih.gov/21150997/

Gonzales MD, Archuleta E, Farmer A, Gajendran K, Grant D, Shoemaker R, Beavis WD, Waugh ME. 2005. The Legume Information System (LIS): An integrated information resource for comparative legume biology. Nucleic Acids Res. [Internet] 33. Available from: https://pubmed.ncbi.nlm.nih.gov/15608283/

Goodstein DM, Shu S, Howson R, Neupane R, Hayes RD, Fazo J, Mitros T, Dirks W, Hellsten U, Putnam N, et al. 2012. Phytozome: A comparative platform for green plant genomics. Nucleic Acids Res. [Internet] 40. Available from: https://pubmed.ncbi.nlm.nih.gov/22110026/

Gordân R, Murphy KF, McCord RP, Zhu C, Vedenko A, Bulyk ML. 2011. Curated collection of yeast transcription factor DNA binding specificity data reveals novel structural and gene regulatory insights. Genome Biol. [Internet] 12. Available from: https://pubmed.ncbi.nlm.nih.gov/22189060/

Guindon S, Gascuel O. 2003. A Simple, Fast, and Accurate Algorithm to Estimate Large Phylogenies by Maximum Likelihood. Rannala B, editor. Syst. Biol. [Internet] 52:696–704. Available from: http://academic.oup.com/sysbio/article/52/5/696/1681984

Hillmer RA, Tsuda K, Rallapalli G, Asai S, Truman W, Papke MD, Sakakibara H, Jones JDG, Myers CL, Katagiri F. 2017. The highly buffered Arabidopsis immune signaling network conceals the functions of its components. PLoS Genet. 13.

Johnson MG, Pokorny L, Dodsworth S, Botigué LR, Cowan RS, Devault A, Eiserhardt WL, Epitawalage N, Forest F, Kim JT, et al. 2019. A Universal Probe Set for Targeted Sequencing of 353 Nuclear Genes from Any Flowering Plant Designed Using k-Medoids Clustering. Syst. Biol. [Internet] 68:594–606. Available from: https://pubmed.ncbi.nlm.nih.gov/30535394/

Katagiri F. 2018. Review: Plant immune signaling from a network perspective. Plant Sci. 276:14–21.

Khan M, Subramaniam R, Desveaux D. 2016. Of guards, decoys, baits and traps: Pathogen perception in plants by type III effector sensors. Curr. Opin. Microbiol. [Internet] 29:49–55. Available from: https://pubmed.ncbi.nlm.nih.gov/26599514/

Kitashiba H, Li F, Hirakawa H, Kawanabe T, Zou Z, Hasegawa Y, Tonosaki K, Shirasawa S, Fukushima A, Yokoi S, et al. 2014. Draft Sequences of the Radish (Raphanus sativus L.) Genome. DNA Res. [Internet] 21:481–490. Available from: https://academic.oup.com/dnaresearch/article-lookup/doi/10.1093/dnares/dsu014

Kumar S, Stecher G, Tamura K. 2016. MEGA7: Molecular Evolutionary Genetics Analysis Version 7.0 for Bigger Datasets. Mol. Biol. Evol. [Internet] 33:1870–1874. Available from: https://pubmed.ncbi.nlm.nih.gov/27004904/

Leebens-Mack JH, Barker MS, Carpenter EJ, Deyholos MK, Gitzendanner MA, Graham SW, Grosse I, Li Z, Melkonian M, Mirarab S, et al. 2019. One thousand plant transcriptomes and the phylogenomics of green plants. Nature [Internet] 574:679–685. Available from: https://pubmed.ncbi.nlm.nih.gov/31645766/

Letunic I, Bork P. 2019. Interactive Tree of Life (iTOL) v4: Recent updates and new developments. Nucleic Acids Res. [Internet] 47. Available from: https://pubmed.ncbi.nlm.nih.gov/30931475/

Li FW, Brouwer P, Carretero-Paulet L, Cheng S, De Vries J, Delaux PM, Eily A, Koppers N, Kuo LY, Li Z, et al. 2018. Fern genomes elucidate land plant evolution and cyanobacterial symbioses. Nat. Plants [Internet] 4:460–472. Available from: https://pubmed.ncbi.nlm.nih.gov/29967517/

Lu Y, Truman W, Liu X, Bethke G, Zhou M, Myers CL, Katagiri F, Glazebrook J. 2018. Different modes of negative regulation of plant immunity by calmodulin-related genes. Plant Physiol. 176:3046–3061.

Ming R, VanBuren R, Liu Y, Yang M, Han Y, Li LT, Zhang Q, Kim MJ, Schatz MC, Campbell M, et al. 2013. Genome of the long-living sacred lotus (Nelumbo nucifera Gaertn.). Genome Biol. [Internet] 14. Available from: https://pubmed.ncbi.nlm.nih.gov/23663246/

Naser-Khdour S, Minh BQ, Zhang W, Stone EA, Lanfear R, Bryant D. 2019. The Prevalence and Impact of Model Violations in Phylogenetic Analysis. Genome Biol. Evol. [Internet] 11:3341–3352. Available from: https://github.com/roblanf/SRHtests/tree/master/datasets

Proost S, Bel M Van, Vaneechoutte D, Van De Peer Y, Inzé D, Mueller-Roeber B, Vandepoele K. 2015. PLAZA 3.0: An access point for plant comparative genomics. Nucleic Acids Res. [Internet] 43:D974–D981. Available from: https://pubmed.ncbi.nlm.nih.gov/25324309/

Qin J, Wang K, Sun L, Xing H, Wang S, Li L, Chen S, Guo HS, Zhang J. 2018. The plant-specific transcription factors CBP60G and SARD1 are targeted by a verticillium secretory protein VDSCP41 to modulate immunity. Elife [Internet] 7. Available from: https://pubmed.ncbi.nlm.nih.gov/29757140/

Reddy VS, Ali GS, Reddy ASN. 2002. Genes encoding calmodulin-binding proteins in the Arabidopsis genome. J. Biol. Chem. 277:9840–9852.

Sato M, Tsuda K, Wang L, Coller J, Watanabe Y, Glazebrook J, Katagiri F. 2010. Network modeling reveals prevalent negative regulatory relationships between signaling sectors in arabidopsis immune signaling. PLoS Pathog. 6:1–15.

Sato S, Hirakawa H, Isobe S, Fukai E, Watanabe A, Kato M, Kawashima K, Minami C, Muraki A, Nakazaki N, et al. 2011. Sequence analysis of the genome of an oil-bearing tree, jatropha curcas L. DNA Res. [Internet] 18:65–76. Available from: https://pubmed.ncbi.nlm.nih.gov/21149391/

Sato S, Nakamura Y, Kaneko T, Asamizu E, Kato T, Nakao M, Sasamoto S, Watanabe A, Ono A, Kawashima K, et al. 2008. Genome structure of the legume, Lotus japonicus. DNA Res.

Singh R, Ong-Abdullah M, Low ETL, Manaf MAA, Rosli R, Nookiah R, Ooi LCL, Ooi SE, Chan KL, Halim MA, et al. 2013. Oil palm genome sequence reveals divergence of interfertile species in Old and New worlds. Nature [Internet] 500:335–339. Available from: https://pubmed.ncbi.nlm.nih.gov/23883927/

Stamatakis A. 2014. RAxML version 8: A tool for phylogenetic analysis and post-analysis of large phylogenies. Bioinformatics [Internet] 30:1312–1313. Available from: https://pubmed.ncbi.nlm.nih.gov/24451623/

Sun T, Liang W, Zhang Y, Li X. 2018. Negative regulation of resistance protein-mediated immunity by master transcription factors SARD1 and CBP60g. J. Integr. Plant Biol. [Internet] 60:1023–1027. Available from: https://pubmed.ncbi.nlm.nih.gov/30007010/

Thompson JD, Higgins DG, Gibson TJ. 1994. CLUSTAL W: Improving the sensitivity of progressive multiple sequence alignment through sequence weighting, position-specific gap penalties and weight matrix choice. Nucleic Acids Res. [Internet] 22:4673–4680. Available from: https://pubmed.ncbi.nlm.nih.gov/7984417/

Toruño TY, Stergiopoulos I, Coaker G. 2016. Plant-Pathogen Effectors: Cellular Probes Interfering with Plant Defenses in Spatial and Temporal Manners. Annu. Rev. Phytopathol. [Internet] 54:419–441. Available from: https://pubmed.ncbi.nlm.nih.gov/27359369/

Truman W, Sreekanta S, Lu Y, Bethke G, Tsuda K, Katagiri F, Glazebrook J. 2013. The CALMODULIN-BINDING PROTEIN60 family includes both negative and positive regulators of plant immunity. Plant Physiol. 163:1741–1751.

Tsuda K, Sato M, Stoddard T, Glazebrook J, Katagiri F. 2009. Network properties of robust immunity in plants. PLoS Genet. 5.

Unver T, Wu Z, Sterck L, Turktas M, Lohaus R, Li Z, Yang M, He L, Deng T, Escalante FJ, et al. 2017. Genome of wild olive and the evolution of oil biosynthesis. Proc. Natl. Acad. Sci. U. S. A. [Internet] 114:E9413–E9422. Available from: https://pubmed.ncbi.nlm.nih.gov/29078332/

Venkatarajan M, Braun W. 2001. New quantitative descriptors of amino acids based on multidimensional scaling of a large number of physical-chemical properties. J. Mol. Model. [Internet] 7:445–453. Available from: http://dx.doi.org/10.1007/s00894-001-0058-5.

Vlot AC, Dempsey DA, Klessig DF. 2009. Salicylic Acid, a Multifaceted Hormone to Combat Disease. Annu. Rev. Phytopathol.

de Vries S, de Vries J, von Dahlen JK, Gould SB, Archibald JM, Rose LE, Slamovits CH. 2018. On plant defense signaling networks and early land plant evolution. Commun. Integr. Biol. [Internet] 11:1–14. Available from: https://www.tandfonline.com/doi/full/10.1080/19420889.2018.1486168

Wang L, Tsuda K, Sato M, Cohen JD, Katagiri F, Glazebrook J. 2009. Arabidopsis CaM binding protein CBP60g contributes to MAMP-induced SA accumulation and is involved in disease resistance against Pseudomonas syringae. Ausubel FM, editor. PLoS Pathog. [Internet] 5:e1000301. Available from: https://dx.plos.org/10.1371/journal.ppat.1000301

Wang L, Tsuda K, Truman W, Sato M, Nguyen L V., Katagiri F, Glazebrook J. 2011. CBP60g and SARD1 play partially redundant critical roles in salicylic acid signaling. Plant J. [Internet] 67:1029–1041. Available from: http://www.ncbi.nlm.nih.gov/pubmed/21615571

Wright ES. Using DECIPHER v2.0 to Analyze Big Biological Sequence Data in R.

Yagi M, Kosugi S, Hirakawa H, Ohmiya A, Tanase K, Harada T, Kishimoto K, Nakayama M, Ichimura K, Onozaki T, et al. 2014. Sequence analysis of the genome of carnation (Dianthus caryophyllus L.). DNA Res.

Zhang L, Chen Fei, Zhang X, Li Z, Zhao Y, Lohaus R, Chang X, Dong W, Ho SYW, Liu X, et al. 2020. The water lily genome and the early evolution of flowering plants. Nature [Internet] 577:79–84. Available from: https://pubmed.ncbi.nlm.nih.gov/31853069/

Zhang Yaxi, Xu S, Ding P, Wang D, Cheng YT, He J, Gao M, Xu F, Li Y, Zhu Z, et al. 2010. Control of salicylic acid synthesis and systemic acquired resistance by two members of a plant-specific family of transcription factors. Proc. Natl. Acad. Sci. U. S. A. [Internet] 107:18220–18225. Available from: https://pubmed.ncbi.nlm.nih.gov/20921422/

Zheng Y, Wu S, Bai Y, Sun H, Jiao C, Guo S, Zhao K, Blanca J, Zhang Z, Huang S, et al. 2019. Cucurbit Genomics Database (CuGenDB): A central portal for comparative and functional genomics of cucurbit crops. Nucleic Acids Res. [Internet] 47:D1128–D1136. Available from: https://pubmed.ncbi.nlm.nih.gov/30321383/

