## Supplemental figs, tables, and files for "Pathogen-driven coevolution across CBP60 plant immune regulator subfamilies confers resilience on the regulator module": FigS7.v10.pdf

## A

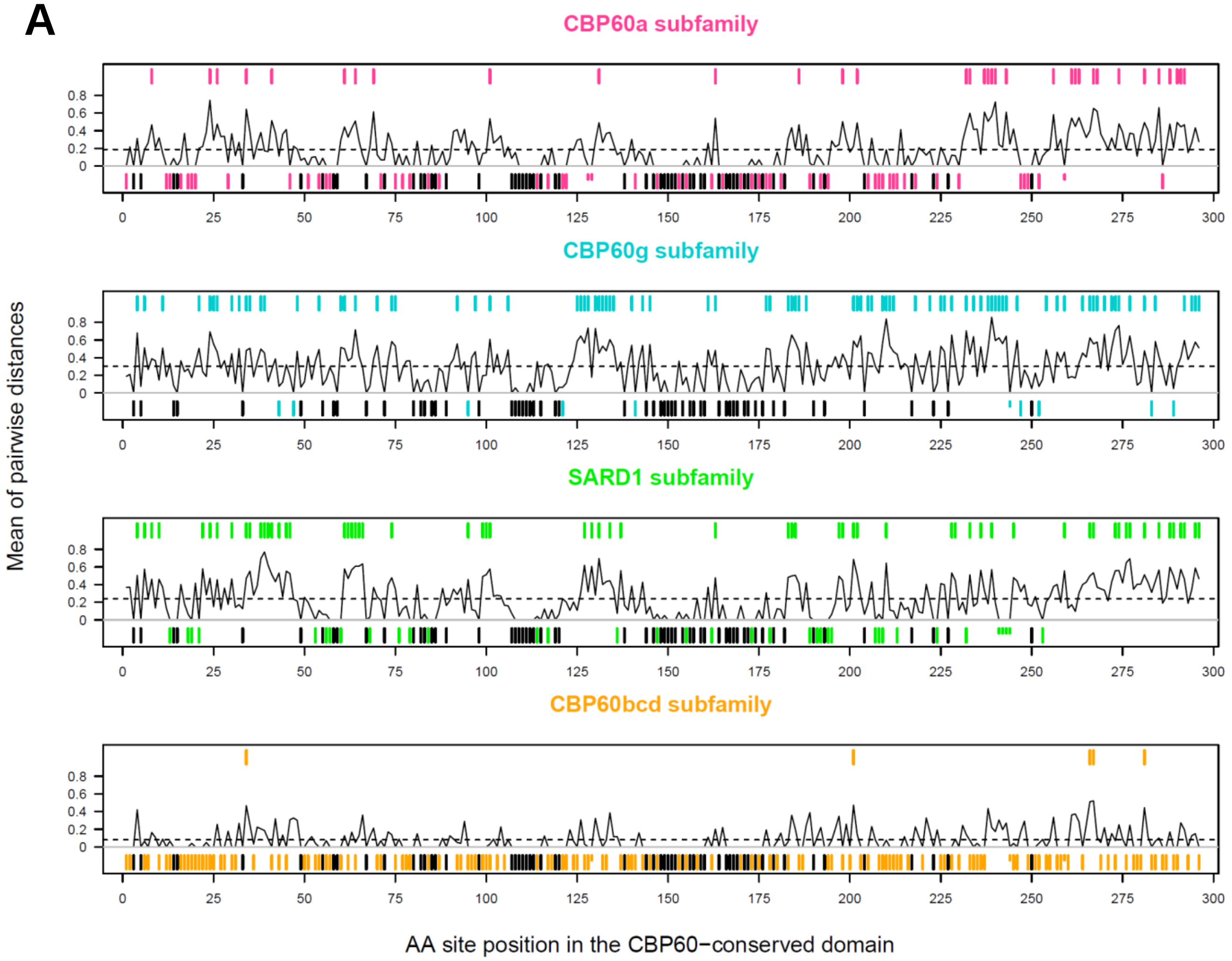


## B

| | CBP60a | CBP60g | SARD1 | CBP60bcd | Benchmark |
|---|---|---|---|---|---|
| Diversity, mean | 0.187 | 0.301 | 0.239 | 0.080 | 0.147 (0.121-0.175) |
| # of highly diverse sites | 34 | 89 | 63 | 5 | 29 (17-44) |
| # of strictly conserved sites | 94 | 42 | 68 | 163 | 129 (107-151) |

Fig S7 – set2
