## Supplemental figs, tables, and files for "Pathogen-driven coevolution across CBP60 plant immune regulator subfamilies confers resilience on the regulator module": parsimonious.substitution.count.r

```
##### parsimonious substitution count algorithm

#### load package
library(seqinr)

#### function for parsimonious substitution count at a node
poss.node.aa = function(branch.aa) {
  br.numb = length(branch.aa)
  all.aa = unlist(branch.aa)
  tab.aa = table(all.aa)
  max.com = max(tab.aa)
  node.aa = names(tab.aa)[tab.aa == max.com]
  ppti.sc = br.numb - max.com
  return(list(subst.numb=ppti.sc, node.aa = node.aa))
}

#### The node connection definition
### The (comopartment and value) numbers in node.c list 
### corresponds to the node number in Fig S13A.
node.c = list()
node.c[1:12] = 1:12
node.c[[13]] = 2:4
node.c[[14]] = 5:6
node.c[[15]] = 8:9
node.c[[16]] = 11:12
node.c[[17]] = 13:14
node.c[[18]] = c(7,15)
node.c[[19]] = c(10,16)
node.c[[20]] = c(1,17)
node.c[[21]] = 18:20

#### common object
## 4 subfamilies of CBP60
subfams = c('CBP60a','CBP60g','SARD1','CBP60bcd')

#################################
#### use the CBP60 conserved domain alignment as an example
cons.eud1 = read.fasta("CBP60.4subfam.12species.fas", seqtype = 'AA')
all.seq.names = names(cons.eud1)

#### alignment per subfam
subfam.cons.eud = list()
for (i in subfams) {
  subfam.cons.eud[[i]] = cons.eud1[grep(i,names(cons.eud1))]
}

#### for each subfam alignment, remove the conserved (including all gap sites) and 
#### make a polymorphic site only alignment matrix
polymor.CBP60 = lapply(subfam.cons.eud, function(sf.aln) {
  aln.mat = t(sapply(sf.aln, function(x) x))
  colnames(aln.mat) = 1:ncol(aln.mat)
  var.numb = apply(aln.mat, 2, function(x) length(unique(x)))
  aln.mat = aln.mat[, var.numb > 1]
  return(aln.mat)
})
sapply(polymor.CBP60, ncol)
## the number of polymorphic sites for each subfamily
#CBP60a   CBP60g    SARD1 CBP60bcd 
#   194      243      224      124

CBP60.pcs = lapply(polymor.CBP60, function(sf.aln) {
  minsens.stat = apply(sf.aln, 2, function(aa.site) {
    var.numb = length(unique(aa.site))
    all.subst.numb = 0
    node.aas = list()
    node.aas[1:12] = aa.site
    for (node.numb in 13:21) {  # these ar
      aa.list = node.aas[node.c[[node.numb]]]
      at.node = poss.node.aa(aa.list)
      node.aas[[node.numb]] = at.node$node.aa
      all.subst.numb = all.subst.numb + at.node$subst.numb
    }
    ppti.sc = all.subst.numb
    return(c(var.numb, ppti.sc))
  })
  rownames(minsens.stat) = c('variant.count', 'par.sub.count')
  return(minsens.stat)
})   

### the list CBP60.pcs has subfamily compartments
### in each compartment, variant number count (row 1) and parsimonious substitution count (row 2)
## e.g., the parsimonious substitution count distribution for each CBP60 subfamily
sapply(CBP60.pcs, function(x) table(x[2,]))
```
