## Supplemental figs, tables, and files for "Pathogen-driven coevolution across CBP60 plant immune regulator subfamilies confers resilience on the regulator module": peaes.pdr.figures.r

```
##### PEAES-PDR analysis
#####

### input, multiple sequence alignment FASTA
# "CBP60.4subfam.12species.fas"
# it is sequences for 4-subfamily members (CBP60a,g,SARD1,60bcd) in 12 selected core eudicot species. 
# aligned to the CBP60 conserved domain from Mapoly (293aa plus 3aa insertions, total 296 sites)

### required .RData
# 'aa.dist.mat.RData' the PEAEA pairwise distance matrix 

#### load packages
library(seqinr)
library(RColorBrewer)

#######################################################
#### all pairwise distance at every site
#### but it will be a huge data, so for each site, summerize the info by
#### 1. mean and upper and lower quartiles
#### 2. every distance value and its number of occurence

load('aa.dist.mat.RData')  # the PEAEA pairwise distance matrix 
aa.dist.mat = round(aa.dist.mat, digits=3)

#### input, multiple sequence alignment FASTA
cons.eud1 = read.fasta("CBP60.4subfam.12species.fas", seqtype = 'AA')
all.seq.names = names(cons.eud1)

cons.epi = list()
seq.numb = c()
sf.names = c('All','CBP60bcd','CBP60a','CBP60g','SARD1')
## each of sf.names class
for (i in sf.names) {
  if (i == 'All') {
    cons.eud = cons.eud1
  } else {
    cons.eud = cons.eud1[grep(i, all.seq.names)]
  }
  cons.eud.mat = t(sapply(cons.eud, function(x) x))
  cons.eud.pwd.info = apply(cons.eud.mat, 2, function(x) {
    n = length(x)
    aa.count = table(x)
    pwd = lapply(1:(n-1), function(y) {
      y.aa = x[y]
      sapply((y+1):n, function(z) {
        aa.dist.mat[y.aa, x[z]]
      })
    })
    pwd= unlist(pwd)
    mean.v = mean(pwd)
    quart.vs = quantile(pwd, probs = c(0.25, 0.75))
    dist.v = table(pwd)
    return(list(mean=mean.v, quartiles=quart.vs, distrib=dist.v,
                aa.count=aa.count))
  })
  cons.epi[[i]] = cons.eud.pwd.info
  seq.numb = c(seq.numb, nrow(cons.eud.mat))
  names(seq.numb)[length(seq.numb)] = i
}  # ~3 sec

quantile(aa.dist.mat[aa.dist.mat!=0],probs = c(0.01, 0.02, 0.05,0.1,0.5,0.9,0.95))
#    1%     2%     5%    10%    50%    90%    95% 
#0.2910 0.3040 0.3820 0.4417 0.7325 1.0553 1.1220 
hcons.th = 0.382/5 # threshold value for "conserved"

### visualization of highly conserved and diverse sites
p.names = c('all CBP60 subfamilies', 'CBP60bcd subfamily', 
            'CBP60a subfamily','CBP60g subfamily', 'SARD1 subfamily')
p.names = paste0('Core Eudicot ', p.names, ', ', seq.numb, ' members')
sf.col = c('black','orange','violetred1','cyan3','green2')

str.cons = list()
nongap.cons = list()
h.div = list()
mm.v = c()

pdf('Fig4.pdf', width=10, height=8)
opar=par(mfrow=c(4,1), mar=c(1,1.7,5,0.4), oma=c(5,5,0.2,0.6))

for (i in 1:5) {
  cons.eud.pwd.info=cons.epi[[sf.names[i]]]
  aa.c.tabs = lapply(cons.eud.pwd.info, '[[','aa.count') ## aa composition counts at every site

  mean.vals = sapply(cons.eud.pwd.info, '[[','mean')
  mm.v = c(mm.v, mean(mean.vals))
  
  # highly conserved sites
  mean.hc = which(mean.vals <= hcons.th)
  # is it a gap, "-"?
  major.aa = sapply(aa.c.tabs, function(x) {
    names(x)[which.max(x)]
  })
  major.gap = major.aa == '-'
  major.nongap = !major.gap[mean.hc]
  str.cons[[sf.names[i]]] = mean.hc
  nongap.cons[[sf.names[i]]] = major.nongap
  if (i == 1) mean.hc.all = mean.hc

  aa.hd = which(mean.vals > 0.4417)  # threshold for "highly diverse"
  h.div[[sf.names[i]]] = aa.hd
  
  if (i != 1) {
    p.title = p.names[i]
    plot(1:length(mean.vals), mean.vals,
         type='l', axes=F,
         xlim = c(6,290),
         ylim = c(-0.25, 1.1), yaxt="n",
         xlab=NA,
         ylab=NA,
         main=p.title, col.main = sf.col[i])
    box(which='plot')
    axis(side = 1, at=seq(0,300,25), labels = seq(0,300,25) )
    axis(side =2, at=seq(0,0.8,0.2), labels=seq(0,0.8,0.2), las=2)
    abline(h=mean(mean.vals), lty=2)
    abline(h=0, col='gray')
    
    segments(mean.hc, -0.1, mean.hc, -0.15-0.1*major.nongap, lwd=2,
             col=sf.col[i])
    segments(mean.hc.all, -0.1, mean.hc.all, -0.25, lwd=2 )
    
    if (length(aa.hd) > 0) {
      segments(aa.hd, 1.09, aa.hd, 0.94, lwd =2,
               col=sf.col[i])
    }
  }
}
title(xlab='AA site position in the CBP60-conserved domain', 
      ylab='Mean of pairwise distances', 
      outer=T, line=3, cex.lab=1.6)
par(opar)
dev.off()

#### stats for each subfamily
names(mm.v) = sf.names
mm.v   # mean of average pairwise distances across the sites, each subfam 
#       All   CBP60bcd     CBP60a     CBP60g      SARD1 
#0.29270872 0.07536614 0.17839087 0.29065474 0.23068054   

sapply(h.div, length)  # the number of highly diverse sites, each subfam
#All CBP60bcd   CBP60a   CBP60g    SARD1 
# 83        5       33       81       58

sapply(sf.names[2:5], function(x) {  # the number of highly conserved sites, non-gap, not-all-cbp-conserved
  cons.spec.ng = setdiff(str.cons[[x]][nongap.cons[[x]]], str.cons[[1]])
  length(cons.spec.ng)
})
#CBP60bcd   CBP60a   CBP60g    SARD1 
#     134       60       13       33

#### common objects
## 12 Core Eudicot species used
sel.sp = c('Quinoa','Coffee','Tomato','Monkeyflower','Carrot','Sunflower',
           'Poplar','Soybean','Peach','Orange','Cotton','Arabidopsis')
## 4 subfamilies of CBP60
subfams = c('CBP60a','CBP60g','SARD1','CBP60bcd')
## all 6 pairwise combinations of 4 subfamilies
subfam.pairs = c('CBP60a:CBP60g', 'CBP60g:SARD1', 'SARD1:CBP60a',
                 'CBP60a:CBP60bcd', 'CBP60g:CBP60bcd', 'SARD1:CBP60bcd')
## the number of sites in the multiple sequence alignment
n = length(cons.eud1[[1]])

###### PEAES-PDR
###### aa similar or dissimilar between two subfamilies at a single site for a species

#### organize the aligned sequences
seq.for.sp = list()
for (sp in sel.sp) {
  seq.for.sp[[sp]] = all.seq.names[grep(sp,all.seq.names)]
}
## subfam.seqs: 12 species x 296 sites matrix in each subfamily, list component
subfam.seqs = list()  
for (sf in 1:3) {
  sf.memb = sapply(seq.for.sp, '[[', sf)
  sf.seq = cons.eud1[sf.memb]
  sf.seq = t(sapply(sf.seq, function(x) x))
  rownames(sf.seq) = sf.memb
  subfam.seqs[[subfams[sf]]] = sf.seq
}

#### select only sites with >= 12 variant combinations between two subfams 
vari.co = 12 # cutoff for 12 variant combinations

vari.numb = c()  # 4 subfamiles x 296 sites matrix for the number of aa variants
for (sf in 1:length(subfams)) {
  sf1.memb = sapply(seq.for.sp, '[[', sf)
  sf1.seq = cons.eud1[sf1.memb]
  sf1.seq = sapply(sf1.seq, function(x) x)
  vari.numb = rbind(vari.numb, apply(sf1.seq, 1, function(x) length(unique(x))))
}
rownames(vari.numb) = subfams

v.sel.m = c()  # the sites that meet vari.co, logical matrix: 6 subfam.pairs x 296 sites
for (sfp in subfam.pairs) {
  sfp.split = unlist(strsplit(sfp, split=':'))
  v.sel = vari.numb[sfp.split[1],] * vari.numb[sfp.split[2],] >= vari.co
  v.sel.m = rbind(v.sel.m, v.sel)
}
rownames(v.sel.m) = subfam.pairs

to.sites.pair = apply(v.sel.m, 1, sum)
to.sites.pair # total number of sites that passed between each subfam.pair
#CBP60a:CBP60g    CBP60g:SARD1    SARD1:CBP60a CBP60a:CBP60bcd CBP60g:CBP60bcd  SARD1:CBP60bcd 
#          122             128             101              43              65              50 

#### distance at every site between two sequences from pairs of subfamilies
# dist.sfpx[[6 subfam.pair]][[296 site.position]]
# a PEAES distance matrix, 12 species, subfam1 x 12 species, subfam2 at the site
dist.sfpx = list()
for (sfp in subfam.pairs) {
  sfp.split = unlist(strsplit(sfp, split=':'))
  sf1 = which(subfams == sfp.split[1]) 
  sf2 = which(subfams == sfp.split[2]) 
  sf1.memb = sapply(seq.for.sp, '[[', sf1)
  sf2.memb = sapply(seq.for.sp, '[[', sf2)
  sf1.seq = cons.eud1[sf1.memb]
  sf2.seq = cons.eud1[sf2.memb]
  dist.sf12 = as.numeric(sapply(sf1.seq, function(x) {
    d.sf12 = as.numeric(sapply(sf2.seq, function(y) {
      apply(cbind(x,y), 1, function(z) aa.dist.mat[z[1],z[2]])
    }))
    return(d.sf12)
  }))
  dist.sf12 = array(dist.sf12, dim=c(n, length(sf2.memb), length(sf1.memb)))
  dimnames(dist.sf12) = list(1:n, sf2.memb, sf1.memb)
  dist.sf12 = lapply(1:n, function(x) t(dist.sf12[x,,]))
  dist.sfpx[[sfp]] = dist.sf12
}

##### PDR
### wisp.sfp[[6 subfam.pair]][[296 site.position]] list:
# a matrix, c('rank','category','sd') x 12 species
# 'rank" is PDR, 'sd' is SD of all permuted (including non-permuted) distances
# 'category', not used in the final version
# NA for all, if there is no distance variation at the site.
wisp.sfp = lapply(dist.sfpx, function(x) {
  return(lapply(x, function(y) {
    wi.rank = sapply(1:length(sel.sp), function(z) {
      other.sp = (1:length(sel.sp))[-z]
      other.v = c(y[other.sp,z], y[z,other.sp])
      if (sum(other.v == y[z,z]) == length(other.v)) {
        rec.rank = NA; rec.cat = NA; rec.sd = NA;
      } else {
        rec.rank = rank(c(y[z,z], other.v))[1]
        rec.sd = sd(c(y[z,z], other.v))
        if (sum(other.v > y[z,z]) == 0) {
          rec.cat = 3
        } else {
          if (sum(other.v < y[z,z]) == 0) {
            rec.cat = 1
          } else {
            rec.cat = 2
          }
        }
      }
      return(c(rank=rec.rank, category=rec.cat, sd=rec.sd))
    })
    colnames(wi.rank) = sel.sp
    return(wi.rank)
  }))
})

###############
r.sd = unlist(lapply(wisp.sfp, function(x) {
  lapply(x, function(y) {
    y['sd',]
  })
}))
hist(r.sd)
quantile(r.sd, na.rm=T)
#          0%          25%          50%          75%         100% 
#0.0004170288 0.1311481527 0.1911211668 0.2602463412 0.5991143463
sd.th = quantile(r.sd, na.rm=T, probs = 0.5) # use the site PDR for r.sd > 50%-tile

### rank.sfp1[[6 subfam.pair]], a matrix of PDR ('rank') for 12 species x 296 sites
# PDR ('rank') = NA when SD <= sd.th
rank.sfp1 = lapply(wisp.sfp, function(x) {
  r.val = sapply(x, function(y) y['rank',])
  sd.val = sapply(x, function(y) y['sd',])
  r.val[sd.val <= sd.th] = NA
  return(r.val)
})

### rank.sp1: reorganize rank.sfp1
# rank.sp1[[12 species]], a PDR matrix (NA included) for 296 sites x 6 subfam.pairs
rank.sp1 = list()
for (sp in sel.sp) {
  rank.sp1[[sp]] = sapply(rank.sfp1, function(x) {
    x[sp,]
  })
}

#### use Fisher test (2-sided) for p-value (1st&3rd quadrants over 2nd&4th quadrants)
#### PC1 for the line (positive PC1 means 1st&3rd quadrants enriched)

comb.m = matrix(c(1,2,2,3,3,1), nrow=2) # 3 immune subfam.pair combinations
p.val.comp = list()  # p.val.comp[[3 immune subfam.pair]], p-values for 12 species
for (combn1 in 1:ncol(comb.m)) {
  x.sfp = subfam.pairs[comb.m[1,combn1]]
  y.sfp = subfam.pairs[comb.m[2,combn1]]
  p.vals = c()
  for (sp in sel.sp) {
    r.x = rank.sp1[[sp]][,x.sfp]
    r.y = rank.sp1[[sp]][,y.sfp]
    r.xy.nna = !is.na(r.x) & !is.na(r.y)
    r.x = r.x[r.xy.nna]
    r.y = r.y[r.xy.nna]
    r.x.small = r.x < 12
    r.y.small = r.y < 12
    cont.tab = table(r.x.small, r.y.small)
    p.val = fisher.test(cont.tab)$p.value
    p.vals = c(p.vals, p.val)
  }
  names(p.vals) = sel.sp
  p.val.comp[[combn1]] = p.vals
}
ad.p.vals = lapply(p.val.comp, p.adjust, method='BH') # p-val correction by BH-FDR

### generate Fig S7, 12 species, immune subfam-common plots in each page 
pdf('FigS7.pdf', width=7, height=9)
opar=par(mfrow=c(4,3), mar=c(0,0,1.5,1.5)+0.1, oma=c(6,6,0,0)+0.1)
count=0
x2 = c(0.5,23.5)
for (combn1 in 1:ncol(comb.m)) {
  x.sfp = subfam.pairs[comb.m[1,combn1]]
  y.sfp = subfam.pairs[comb.m[2,combn1]]
  count1=0
  for (sp in sel.sp) {
    plot(0,0, type='n',
         xlim=c(-2,27), ylim=c(-2,27),
         axes=F)
    axis(side = 1,
         at=seq(0,24,6),
         labels = (count1 %/% 3 == 3) )
    axis(side = 2, las=2,
         at=seq(0,24,6),
         labels = (count1 %% 3 == 0))
    text(0,25, sp, pos=4, cex=1.4)
    segments(x2[1],12,x2[2],12, lty=3)
    segments(12,x2[1],12,x2[2], lty=3)
    r.x = rank.sp1[[sp]][,x.sfp]
    r.y = rank.sp1[[sp]][,y.sfp]
    r.xy.nna = !is.na(r.x) & !is.na(r.y) # not NA for both axes
    r.x = r.x[r.xy.nna]
    r.y = r.y[r.xy.nna]
    points(r.x, r.y,
           col=sf.col[(c(4,5,3))[combn1]], cex=0.7)
    
    count = count+1
    p.val = ad.p.vals[[combn1]][sp]
    text(25, -0.5, paste0('p = ',formatC(p.val, format = "e", digits = 2)), pos=2, cex=1.3)
    if (p.val < 0.05) {
      xy.mat = cbind(r.x, r.y)
      pca.xy =prcomp(xy.mat)
      pc1.slope = pca.xy$rotation[,'PC1'][2]/pca.xy$rotation[,'PC1'][1]
      pc1.interc = pca.xy$center['r.y'] - pca.xy$center['r.x']*pc1.slope 
      if (pc1.slope <= 1) {
        segments(x2[1],pc1.interc+ pc1.slope*x2[1],x2[2],pc1.interc+ pc1.slope*x2[2], lty=2)
        if (p.val < 0.01) segments(x2[1],pc1.interc+ pc1.slope*x2[1],x2[2],pc1.interc+ pc1.slope*x2[2])
      } else {
        segments((x2[1]-pc1.interc)/pc1.slope,x2[1],(x2[2]-pc1.interc)/pc1.slope,x2[2], lty=2)
        if (p.val < 0.01) segments((x2[1]-pc1.interc)/pc1.slope,x2[1],(x2[2]-pc1.interc)/pc1.slope,x2[2])
      }
    }
    count1=count1+1
  }
  title(xlab=paste0('PDR, ',x.sfp), 
        ylab=paste0('PDR, ',y.sfp), 
        outer=T, line=3, cex.lab=1.6)
}
par(opar)
dev.off()


##### distribution of the PDR compared sites
## color gradients corresponding to Fig 6A (below)
col.grad1 = colorRampPalette(c("red1","gray","blue"))
col.grad1 = col.grad1(11)
col.grad2 = colorRampPalette(c("magenta2","gray","gold"))
col.grad2 = col.grad2(11)

## generate Fig 7
pdf('Fig7.pdf', width=7, height=9)
opar=par(mar=c(5,6,1,1))
plot(0,0, type='n', axes=F,
     xlim = c(1, nrow(rank.sp1[[1]])),
     ylim = c(-0.5, length(sel.sp)),
     xlab = 'AA site position in the CBP60-conserved domain',
     ylab = NA
)
axis(side=1)

for (combn1 in 1:2) {
  x.sfp = subfam.pairs[comb.m[1,combn1]]
  y.sfp = subfam.pairs[comb.m[2,combn1]]
  for (sp.n in 1:length(sel.sp)) {
    sp = sel.sp[sp.n]
    sp.pos1 = length(sel.sp) - sp.n - 0.33 * (combn1-1) 
    r.x = rank.sp1[[sp]][,x.sfp]
    r.y = rank.sp1[[sp]][,y.sfp]
    mean.vs = apply(rbind(r.x, r.y), 2, function(x) {
      if (sum(is.na(x)) > 0) {
        mean.v = c(NA, NA)
      } else {
        if ((x[1]-12) * (x[2]-12) >= 0) {
          mean.v = c(mean(x), NA)
        } else {
          mean.v = c(NA, mean(24-x[1], x[2]))
        }
      }
      return(mean.v)
    })
    
    if (combn1 ==1) {
      mtext(sp, side=2, line=0, at=sp.pos1+0.26, adj=1, las=1,cex=0.9)
      mtext('CBP60g-common', side=2, line=0, at=sp.pos1, adj=1, las=1, cex=0.7, col=sf.col[4])
    } else {
      mtext('SARD1-common', side=2, line=0, at=sp.pos1, adj=1, las=1, cex=0.7, col=sf.col[5])
    }
    
    arrows(1,sp.pos1, nrow(rank.sp1[[sp]]), sp.pos1, angle=90, code=3, length=0.03)
    segments(1:nrow(rank.sp1[[sp]]), rep(sp.pos1,nrow(rank.sp1[[sp]]))-0.12, 
          1:nrow(rank.sp1[[sp]]), rep(sp.pos1,nrow(rank.sp1[[sp]]))+0.12,
          col=col.grad1[round(mean.vs[1,]/2)], lwd=2)

   
    arrows(1,sp.pos1, nrow(rank.sp1[[sp]]), sp.pos1, angle=90, code=3, length=0.03)
    segments(1:nrow(rank.sp1[[sp]]), rep(sp.pos1,nrow(rank.sp1[[sp]]))-0.12, 
             1:nrow(rank.sp1[[sp]]), rep(sp.pos1,nrow(rank.sp1[[sp]]))+0.12,
             col=col.grad2[round(mean.vs[2,]/2)], lwd=2)
  }
}
par(opar)
dev.off()

#### color scale for 'Fig7.pdf'
opar=par(mfrow=c(4,3), mar=c(5,5,1,1))
sp = sel.sp[1]
plot(0,0, type='n',
     xlim=c(0,25), ylim=c(0,25),
     xlab=paste0('PDR, ',x.sfp),
     ylab=paste0('PDR, ',y.sfp), axes=F)
box(which='plot')
axis(side = 1, at=seq(0,24,6), labels = T )
axis(side =2, at=seq(0,24,6), labels = T, las=2)

for (i in seq(0.01,22,0.01)) {
  if (i <=11 ) {
    segments(12,i+12,i+12,12, col=col.grad1[round(i/4+6)])
  } else {
    segments(i+1,23,23,i+1, col=col.grad1[round(i/4+6)])
  }
  if (i <=11 ) {
    segments(1,i+1,i+1,1, col=col.grad1[round((i+2)/4)])
  } else {
    segments(i-10,12,12,i-10, col=col.grad1[round((i+2)/4)])
  }
  
  if (i <=11 ) {
    segments(12,12-i,i+12,12, col=rev(col.grad2)[round(i/4+6)])
  } else {
    segments(i+1,1,23,23-i, col=rev(col.grad2)[round(i/4+6)])
  }
  if (i <=11 ) {
    segments(1,23-i,i+1,23, col=rev(col.grad2)[round((i+2)/4)])
  } else {
    segments(i-10,12,12,34-i, col=rev(col.grad2)[round((i+2)/4)])
  }
}
abline(v=12, lty=3)
abline(h=12, lty=3)
r.x = rank.sp1[[sp]][,x.sfp]
r.y = rank.sp1[[sp]][,y.sfp]
r.xy.nna = !is.na(r.x) & !is.na(r.y)
r.x = r.x[r.xy.nna]
r.y = r.y[r.xy.nna]
points(r.x, r.y,
       col=combn1+1, cex=0.7)
par(opar)
# saved as 'Fig6A.pdf'
```
