## Supplemental figs, tables, and files for "Pathogen-driven coevolution across CBP60 plant immune regulator subfamilies confers resilience on the regulator module": TextS1.pdf

Supplemental Text 1

**Examples of CBP60 homologs that were not included in this study**

In the study described in the main text, the CBP60 protein family members were operationally defined by the range of diversity among the Arabidopsis CBP60 members. Here we call this CBP60 family member set the Arabidopsis-defined set (Ad set). The diversity in the protein sequence set was limited to assure reasonable accuracy in phylogenetic analysis and to allow functional interpretations of the subfamilies. Consequently, there were some CBP60 homologs that were omitted from the Ad set.

To show examples of CBP60 homologs omitted in the Ad set, we selected four Angiosperm species with high quality genome sequences and annotations, Arabidopsis, Tomato, Rice, and Maize. Each of the eight Arabidopsis CBP60 members was used as a BLASTP query against the annotated protein sequences of these four genomes. Instead of using the furthest Arabidopsis member as the bit score cutoff in each BLASTP search as for the Ad set selection, the following arbitrary cutoff conditions were used: query alignment length > 250 AND bit score > 100. Then the union of sequences selected from the BLASTP search results for each query was taken as an expanded set.

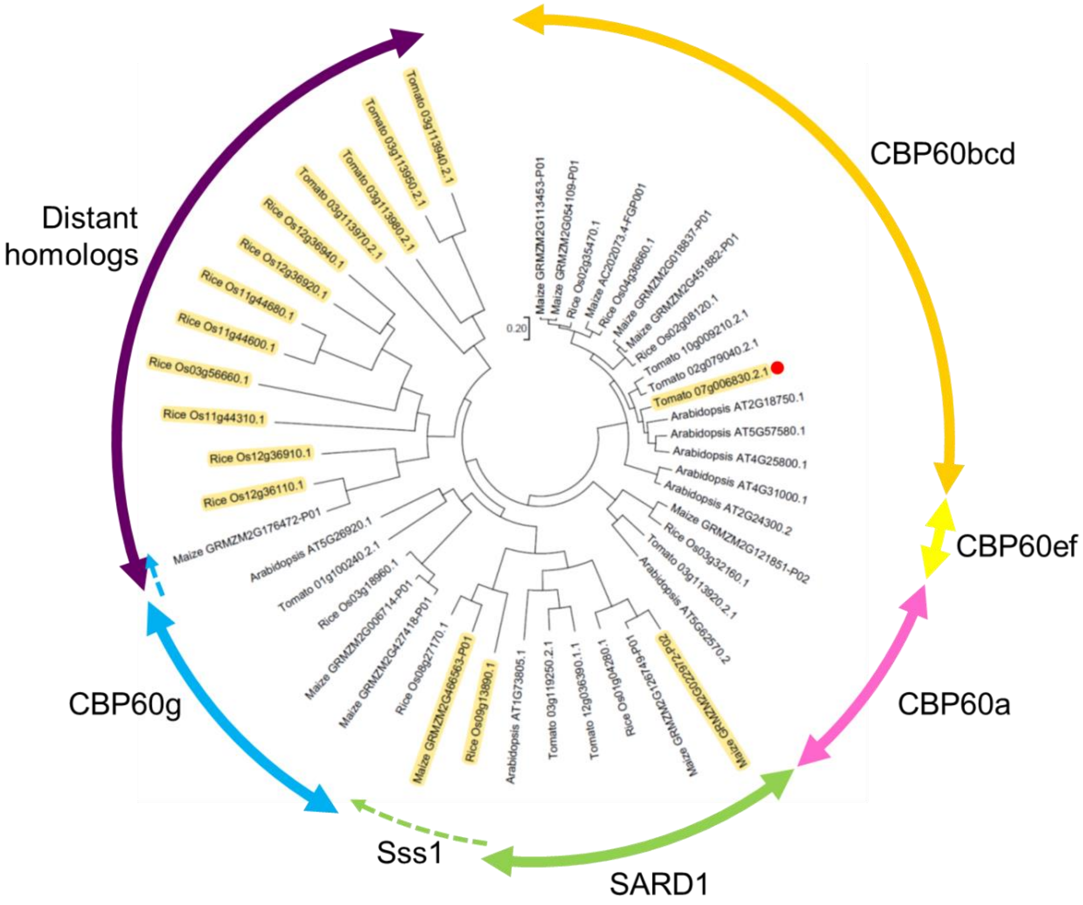


Fig TS1A. Phylogenetic tree and subfamily membership annotations with the expanded set.

The sequences in the expanded set were aligned using ClustalW (with the default parameter values) and the phylogenetic tree was inferred using the Maximum Likelihood method in MEGA 7 (with the default parameter values except for the Site Coverage Cutoff, which was 70%) [1].

Fig TS1A shows the phylogenetic tree made using the expanded set, annotated with the subfamily memberships. The yellow highlighted leaves are the CBP60 homologs that were omitted in the Ad set. Among these, Tomato 07g006830.2.1 (indicated by a red dot) was omitted because it did not pass the sequence quality control following the BLASTP cutoff (Text S2). One Maize SARD1 member (Maize GRMZM2G022972-P02) was omitted although the other Maize SARD1 member (Maize GRMZM2G126749-P01) was included. The former is a little more diverse than the latter, and the bit score with AtSARD1 (Arabidopsis AT1G73805.1) as a query was lower than any of the Arabidopsis members. This indicates that some true but diverse members of the SARD1 or CBP60g subfamilies may be omitted from the Ad set. For this reason, we carefully examined the BLASTP results beyond the cutoff bit scores before stating that a particular species may lack members of a particular subfamily. Monocots have members of a SARD1 sub-subfamily (Sss1), and some members are included while others are omitted from the Ad set. Since eudicots, including Arabidopsis, do not have Sss1 sub-subfamily members, we did not do anything special about the Sss1 members. Sss1 members in the Ad set were annotated as SARD1 subfamily members in the main text. There is a clade of "Distant homologs", which mostly consists of those omitted from the Ad set. Whereas Tomato has members of the distant homolog clade, Arabidopsis does not. Apparently the distant homolog clade was diversified after divergence of the CBP60a subfamily but before diversification between the CBP60g and SARD1 subfamilies in the immune-related clade. It should be noted that the common ancestors of these subfamilies were quite different from the subfamily members of the extant species, so that the apparent relationships among the distant homolog clade and the immune regulator CBP60 subfamilies may not be accurate. One maize member of the distant homolog clade (Maize GRMZM2G176472-P01) was included in the Ad set and annotated as a CBP60g subfamily member in the main text. Existence of highly divergent clades, such as the distant homolog clade, and inclusion of a small number of distant homolog clade members in the Ad set may cause minor inaccuracy in the phylogenetic analysis.

One conclusion that can be drawn from the above analysis of the expanded and Ad sets is that there is no universal way to select protein family members solely based on similarity measures. Note that the alignment length is a similarity measure. Whatever similarity measure cutoffs are used, there will be some false positives (e.g., a distant homolog member included in the Ad set) and false negatives (e.g., a SARD1 subfamily member omitted from the Ad set). This situation with false positives and false negatives is exacerbated by highly diverse evolution rates within a single set of sequences, as in the CBP60 family.

### REFERENCE

1. Kumar, S., Stecher, G., and Tamura, K. (2016). MEGA7: Molecular Evolutionary Genetics Analysis Version 7.0 for Bigger Datasets. Mol. Biol. Evol.
