## Supplemental figs, tables, and files for "Pathogen-driven coevolution across CBP60 plant immune regulator subfamilies confers resilience on the regulator module": TextS2.pdf

Supplemental Text 2. Detailed methods and the features of the CBP60 protein sequence phylogeny that are not discussed in the main text.

### Protein Database Assembly

We downloaded protein sequences from 271 land plant genomes [1–16]. Table S1 contains the list of all species used, together with information on whether the genome or transcriptome was sequenced and where the protein sequence data was downloaded from. We then used the NCBI BLAST+ suite to create a searchable protein database. Initially, we used BLASTP to search this database using Arabidopsis CBP60 proteins as queries. Since the commonly used e-value depends on database size, we used the bit score value as the library-independent similarity measure. In addition, the bit score allows the sum of the values from multiple non-overlapping HSPs to be used as the similarity measure of an entire sequence. For each BLASTP search with each of the eight Arabidopsis CBP60 members, we ranked the results by bit score and selected all proteins with a higher bit score than the bit score of the lowest-ranked Arabidopsis CBP60 member. This Furthest Member Selection (FMS) approach yielded 1432 proteins. Some CBP60 members were omitted due to technical issues, such as inaccurate sequence assembly or annotation. We did not attempt to correct these potentially inaccurate and omitted protein sequences. Some distantly-related CBP60 homologs were omitted, as described in Text S1.

### Quality control of the identified CBP60 sequences

We performed multiple steps of quality control on this list of proteins to exclude potentially inaccurate sequences as inaccurate sequences lower the quality of multiple sequence alignment. The first QC step was using the conserved domain of Liverwort_0018s0017 CBP60 protein, which was the sequence closest to the most recent common ancestor (MRCA) of all land plant CBP60 sequences we identified, as a BLASTP query and removing all proteins that had low homology or too many gaps in the alignment. Criteria for low homology were: percentage of identity < 40.0, bit score < 200, alignment length < 250 or total gaps > 20. Additionally, we removed those proteins that had overly long sequences N-terminal or C-terminal to the CBP60 conserved domain (> 200 or > 400, respectively). The second QC step was a HMMER scan [17] for the CBP60 conserved domain (PF07887.11) to identify and exclude protein sequences that have multiple units of the CBP60 conserved domain. Such protein sequences cause difficulties in multiple sequence alignment. Third, we performed a HMMER search to identify and exclude proteins with additional PFAM domains. We used i-evalue cutoff of < 0.01 or domain score > 15 as thresholds for additional domain significance. Protein sequences excluded due to multiple CBP60 conserved domains or an additional significant PFAM domain (other than the CBP60 conserved domain), were most likely results of assembly or annotation errors since we generally did not observe CBP60 sequences with similar domain organizations in closely related species.

The fourth QC step was reducing our protein data set so that each genus is represented by proteins from only one species. The intention of this QC was to achieve relatively balanced diversity representations in different parts of the sequence phylogenetic tree for better accuracy in tree inference. When a genus was represented by more than one species, we kept the species with the likely highest quality sequences according to the following rules in this order: (i) A dataset generated by genome, rather than transcriptome, sequencing was given higher priority; (ii) A dataset specified only by a genus name but not by a specific species name was given lower priority; (iii) A dataset with a higher total protein sequence length was given higher priority. Table S2, Sheet 1 lists the final set of 1,024 CBP60 protein sequences and Sheet 2 lists the sequences excluded by the QCs and the reasons for their exclusion.

**Protein sequence phylogenetic tree inference procedure, general**

In the subsequent tree inference procedures, ClustalW [18], a Maximum Likelihood (ML) method [19], and iTOL [20] were used for multiple sequence alignment, phylogenetic tree inference, and tree visualization, respectively. ML was also used for inference of most probable ancestor sequences at branching points in a tree. MEGA7 [21] was used for ClustalW and ML with the default parameter values, except for 80% site coverage in ML (unless stated otherwise).

The ancestral species lineage starting from the MRCA of all land plants leading to Arabidopsis is called the land plant lineage. The plant groups, Liverwort, Mosses, Lycophytes, Ferns, and Gymnosperms, diversified from the land plant lineage as monophyletic groups in this order [22]. Within Angiosperms, Basal Angiosperms diversified as a paraphyletic group from the land plant lineage first, followed by Magnoliids/Chloranthales and Monocots [23]. See Fig S3 for the taxonomic order phylogeny within Eudicots.

In the CBP60 protein sequence phylogeny, the ancestral lineage starting from the MRCA of all 1024 land plant CBP60s and leading to the divergence of the CBP60ef, CBP60bcd, CBP60a, CBP60g, and SARD1 subfamilies is called the main CBP60 lineage. After diversification of the Fern sequence clade, the main CBP60 lineage was split into two due to diversification of the immune-related clade, which later diversified into the CBP60a, CBP60g, and SARD1 subfamilies, from the prototypical group, which later diversified into the CBP60ef and CBP60bcd subfamilies (see below).

**Evaluation of the topology of a protein sequence phylogenetic tree**

Instead of evaluating the tree statistically, such as by using the bootstrap value, we evaluated the consistency of the obtained tree by visual comparison to the topology of the organismal species phylogenetic tree. The main reason for this was that some clades of CBP60s are under strong selection and not well explained by standard evolution models or values calculated based on standard evolution models, such as the bootstrap value. Another reason is that the topology of a protein family sequence phylogenic tree is highly predictable based on an organismal species phylogenic tree. Barring horizontal gene transfer and recombination between sequences of distantly related subfamilies, which are very rare events in plants, deviation of the sequence tree topology from the species tree topology should be explained by duplication and deletion of sequences. The extent to which this rule of species phylogeny, duplication, and deletion (SPDD rule) can explain the high-level topology of a sequence phylogenetic tree was used as the tree evaluation criterion.

### Challenges in phylogenetic tree inference of CBP60 sequences

From preliminary studies, we identified several challenges in inferring a CBP60 sequence phylogenetic tree. First, several clades, including the immune regulator subfamilies and a Fern sequence subfamily (hereafter referred to as Fern subfamily 1), have sequences very different from those of many prototypical members in the region C-terminal to the CBP60-conserved domain. Thus, inclusion of the C-terminal region results in inaccurate multiple sequence alignment across the clades in the large C-terminal region. The members of the SARD1 subfamily and the Fern subfamily 1 lack most of the C-terminal region. Since site-by-site information is used in tree inference, strong dissimilarity is assumed between the clades with the C-terminal region and those without. Second, the diversity of the evolution rate across the clades is very high. For example, the evolution rates of the immune regulator CBP60g and SARD1 subfamilies are much higher than most of the clades in the prototypical group. The issues discussed above regarding the first challenge of a high level of variation in the C-terminal region would favor use of the CBP60-conserved domain only in tree inference. However, the second challenge presents issues associated with focusing only on the CBP60-conserved domain. The CBP60-conserved domains are highly conserved among most of the prototypical members and thus phylogenetic information would be very limited within the prototypical group. On the other hand, variant sites in the CBP60-conserved domain have been evolving very fast among the immune regulator subfamilies, and many variant sites underwent multiple substitution events. Multiple substitutions per site make phylogenetic inference inaccurate. Furthermore, many of these sites have been under strong selection, as described for Eudicot immune regulator subfamilies in the main text. Evolution of strongly selected sites cannot be inferred well based on standard evolution models, which are commonly used in sequence alignment and tree inference methods.

### Inference of CBP60 sequence phylogenetic tree using a MRCA-Anchored, Clade-by-Clade Tree Inference (MACCTI) strategy

With all these challenges considered, we developed a tree inference strategy in which the aligned sequence region used for tree inference is locally adjusted. We named this strategy MRCA-Anchored, Clade-by-Clade Tree Inference (MACCTI). In short, an overview tree for relationships among all major clades and MRCAs on the main CBP60 lineage was inferred only using the sequences with moderately alignable C-terminal regions, the sequences of each clade and its flanking MRCAs on the main CBP60 lineage were used to infer the tree for the clade, the MRCAs of the clades were used to infer the backbone tree showing the relationships among the clade MRCAs, and the clade MRCAs in the backbone tree were replaced with the corresponding clade trees to reconstruct the entire tree with all sequences.

In the first step of MACCTI, an overview tree was inferred using only sequences longer than 524 amino acids. The length cutoff of 524 amino acids was determined based on the length distribution among the 1024 QC-ed sequences: the length of 524 amino acids is the middle of the valley between the two peaks in the length distribution of the sequences (Fig T2S1). The idea is to select only sequences with substantial lengths in the C-terminal region, so that almost the entire lengths of the sequences can be used for tree inference. Note that this length selection removed all members of the SARD1 subfamily and the Fern subfamily 1.

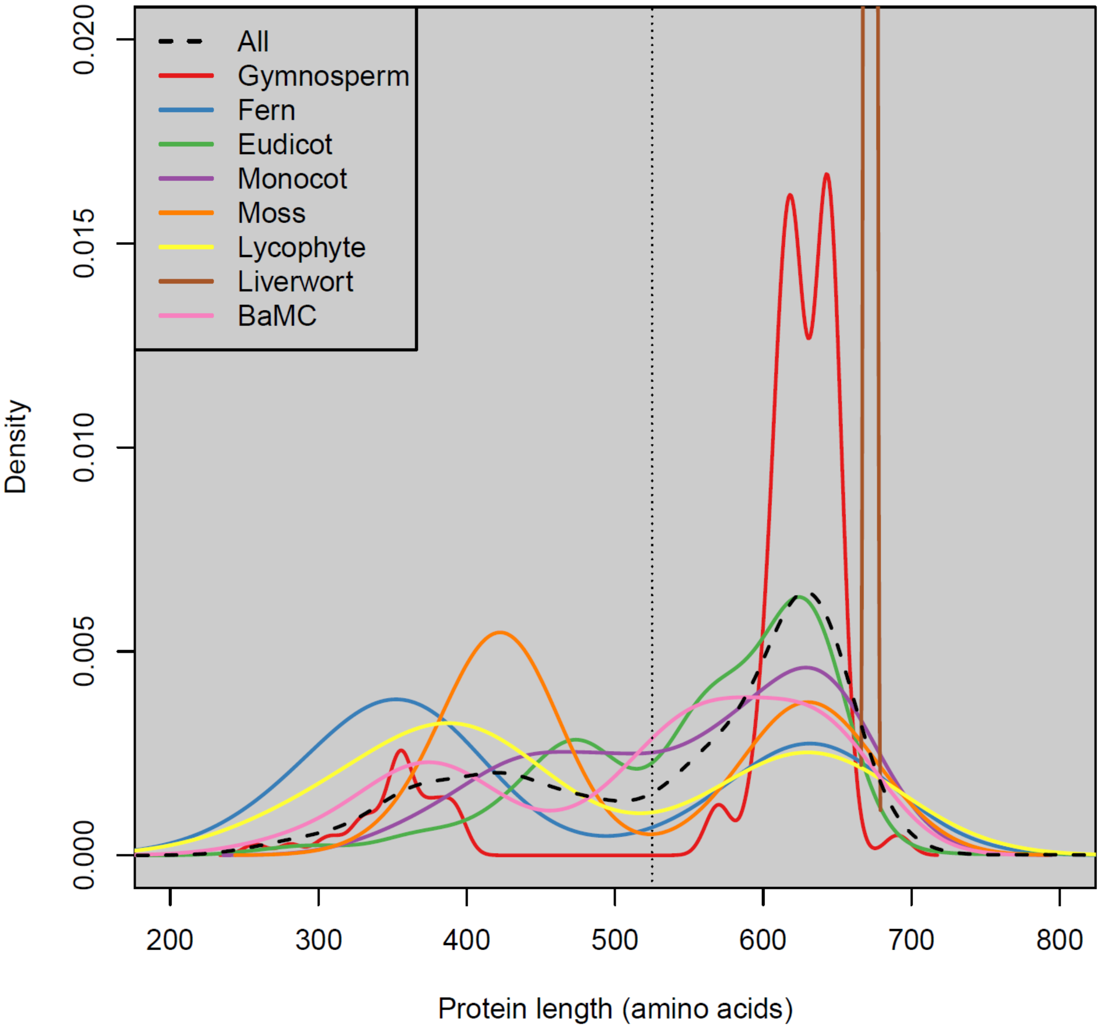


Fig T2S1. The size distributions of the CBP60 sequences of the land plant groups. BaMC: Basal Angiosperms, Magnoliids, and Chloranthales. Only three sequences of very similar sizes are in the Liverwort group. The size of 525 amino acids is shown by the vertical dotted line.

There were 658 sequences longer than 524 amino acids among the 1024 sequences. An ML tree was inferred using 60% site coverage (Fig T2S2). The reason a lower site coverage was chosen was that with the selection of sequences including the C-terminal region, many of the gaps in the alignment of the C-terminal region likely represent true sequence-discriminating information.

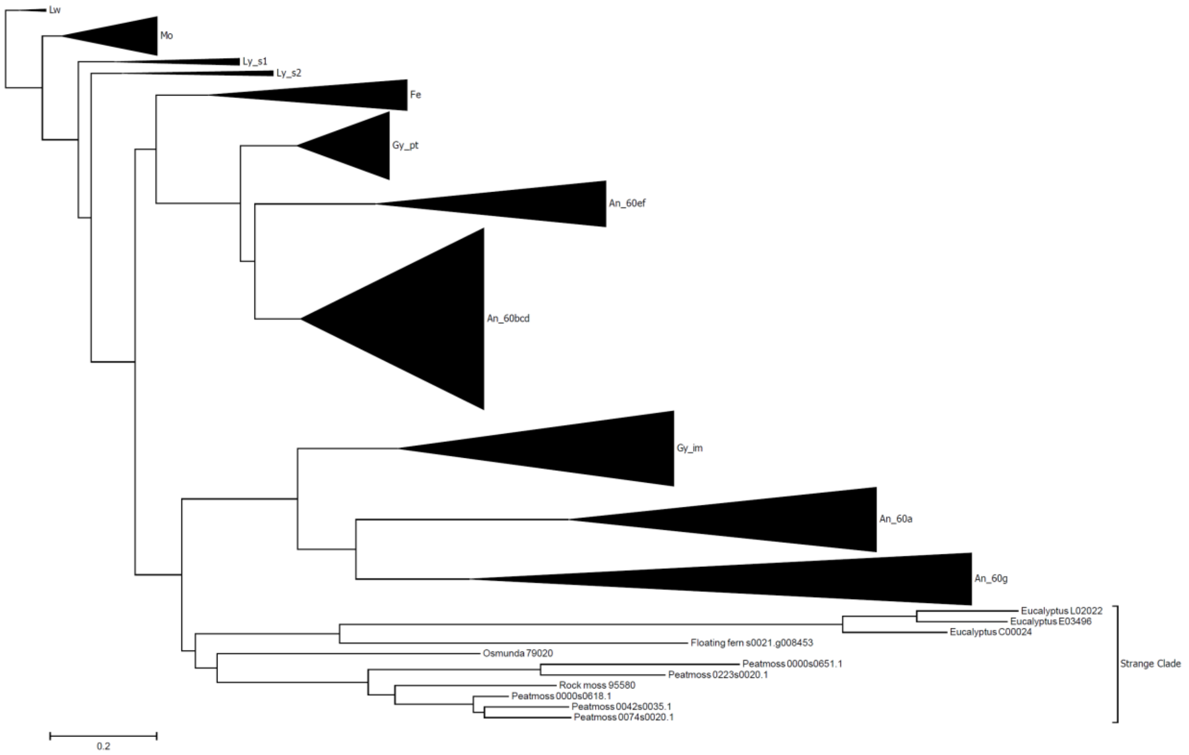


Fig T2S2. An overview tree with 658 sequences longer than 524 amino acids contains a “strange” clade. Only the “strange” clade members are shown at the bottom as individual sequences while other clades are compressed: Liverwort (Lw), Mosses (Mo), Lycophyte subfamily 1 (Ly_s1), Lycophyte subfamily 2 (Ly_s2), Ferns (Fe), Gymnosperm prototypical (Gy_pt), Gymnosperm immune-related (Gy_im), Angiosperm CBP60ef subfamily (An_60ef), Angiosperm CBP60bcd subfamily (An_60bcd), Angiosperm CBP60a subfamily (An_60a), and Angiosperm CBP60g subfamily (An_60g).

The high-level topology of the tree, which describes the relationships among major clades, can be explained by the SPDD rule based on the phylogeny of the major plant groups (Liverwort, Lw; Mosses, Mo; Lycophytes, Ly; Ferns, Fe; Gymnosperms, Gy; and Angiosperms, An), except for one “strange” clade consisting of 11 sequences. This “strange” clade consisted of Peatmoss_0074s0020.1, Peatmoss_0042s0035.1, Peatmoss_0000s0618.1, Rock_moss_95580, Peatmoss_0223s0020.1, Peatmoss_0000s0651.1, Osmunda_79020, Floating_fern_s0021.g008453, Eucalyptus_C00024, Eucalyptus_E03496, and Eucalyptus_L02022. The place of this clade in the tree, which was diversified after diversification of the immune-related clade from the prototypical group and before diversification of the Gymnosperm immune-related clade, could have suggested horizontal transfer of an ancestral immune-related gene to these plant species. However, from the tree topology, this potential horizontal gene transfer would have occurred just once to multiple diverse plant species. The “strange” clade members originated from 2 Moss species, 2 Fern species, and 1 Angiosperm species. In addition, the branch lengths within the “strange” clade were extremely long. We concluded that the “strange” clade is an artifact of tree inference. It was probably due to the tree inference procedure grouping sequences very different from any other sequences even though the sequences were not closely related to each other.

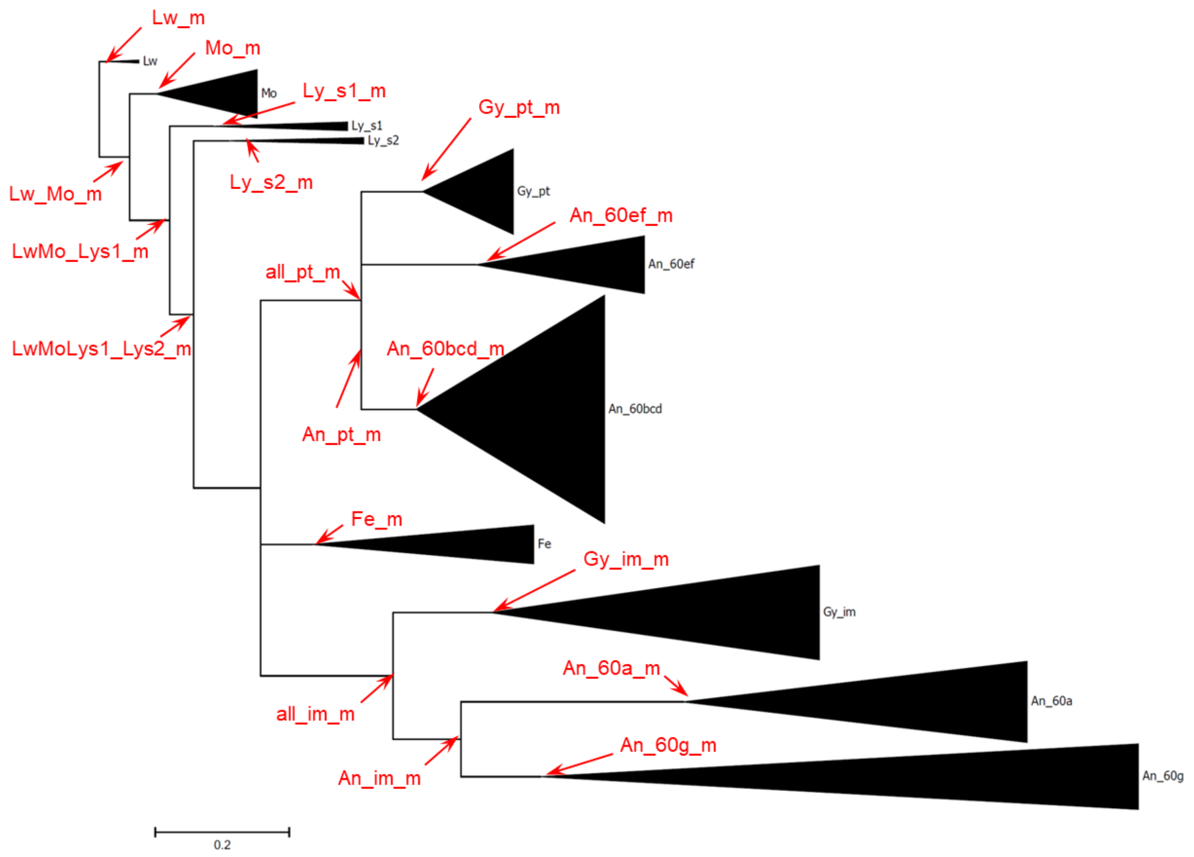


Fig T2S3. The final overview tree with the 647 sequences and the MRCAs inferred (red). The major clades are compressed.

We inferred an overview tree consisting of 647 long sequences (omitting the 11 “strange” clade sequences) using 60% site coverage. In the topology of the obtained tree, two minor modifications were manually made: “Takakia_11547” diversified from the main CBP60 linage directly immediately before divergence of all other Moss sequences, so it was moved closest to the root within the Moss clade; “Coontail_12565” was closest to the MRCA of the Monocot and Eudicot CBP60bcd subfamily members, so it was moved closest to the MRCA of Eudicot CBP60bcd. Then this manually modified tree topology was used as the guide tree for the 647 sequences in ClustalW. The obtained alignment was used for inferring the branch lengths for the manually modified tree topology by ML with 80% site coverage (Fig T2S3). This tree was designated the final overview tree and used to infer the most probable sequences for the MRCAs of the following clades (MRCA labels used are in the parentheses): Lw (Lw_m), Lw and Mo (Lw_Mo_m), Mo (Mo_m), Mo and Ly subfamily 1 (LwMo_Lys1_m), Ly subfamily 1 (Ly_s1_m), Ly subfamily 1 and Ly subfamily 2 (LwMoLys1_Lys2_m), Ly subfamily 2 (Ly_s2_m), Fe (Fe_m), Gy prototypical and An prototypical (all_pt_m), Gy prototypical

(Gy_pt_m), Gy immune-related and An immune regulators (all_im_m), Gy immune-related (Gy_im_m), An prototypical (An_pt_m),An CBP60ef (An_60ef_m), An CBP60bcd (An_60bcd_m), An immune regulators (An_im_m), An CBP60a (An_60a_m), and An CBP60g (An_60g_m). See below for two subfamilies of Ly.

Although short CBP60 sequences were not included, the final overview tree showed: (1) Seed Plants have members of the prototypical group and the immune-related clade; (2) Fe members that are longer than 524 amino acids were monophyletic; (3) the divergence point for the Fe clade was very close to the point where the immune-related clade diversified from the prototypical group; (4) the immune regulator CBP60a and CBP60g subfamilies diversified within the immune-related clade at a time near the divergence of Angiosperms; (5) CBP60 was duplicated on the main CBP60 lineage before divergence of Lycophytes, and both subfamilies were maintained in the Ly group (Ly subfamilies 1 and 2); (6) Only the ancestor of Ly subfamily 2 was maintained in the subsequent main CBP60 lineage.

In the second step of MACCTI, clade-by-clade subtrees were inferred using all sequences belonging to the clade while their flanking MRCAs on the main lineage were used as outgroups to “anchor” the subtrees. Generally, a subtree for a clade was anchored by two inferred MRCA sequences in the main lineage, one was immediately before the time of the clade diversification and the other was immediately after. If a clade is indeed monophyletic, the tree has the topology that the monophyletic group diversified from the part of the main CBP60 lineage delineated by the two MRCAs. If the clade is not monophyletic, we would see it in relation to the two MRCAs.

We inferred an Fe subtree and its divergence point on the main CBP60 lineage. The divergence point of the Fe clade was very close to the point of diversification of the immune-related clade from the prototypical group in the overview tree. We used all 91 Fe CBP60 sequences from the QC-ed set, LyMoLys1_Lys2_m, all_pt_m, and all_im_m. The tree structure clearly showed that: (1) all Fe sequences are monophyletic; (2) the Fe clade diversified from the main CBP60 lineage before diversification of the immune-related clade from the prototypical group; (3) two major subfamilies diversified during very early evolution of Mo; (4) one of the two subfamilies lost the C-terminal region (Fe subfamily 1); (5) Two Fe members of the “strange” clade were placed in the Fe subtree in a way explainable by the SPDD rule.

Next, we inferred Ly subtrees. The overview tree showed that the CBP60 sequence was duplicated in the main CBP60 lineage before divergence of Ly, and one of the duplicated sequences was deleted in the main lineage after divergence of Ly while the Ly lineage maintained both subfamilies originated from the duplication (Figs T2S3 and T2S4). We named the two subfamilies Ly subfamily 1 and Ly subfamily 2 (Ly s1 and Ly s2). The ancestor of Ly s1 was deleted in the main CBP60 linage, and the ancestor of Ly s2 was maintained in the main linage and became the ancestor of all CBP60 sequences evolved subsequently. The two-subfamily topology was clear with the long Ly sequences in the overview tree and was consistent with the SPDD rule. However, once short Ly sequences were added (total 34 sequences), it became difficult to obtain a subtree topology consistent with the SPDD rule using the flanking MRCAs, Lw_Mo_m and Fe_m. Therefore, we considered the two-subfamily structure to be true even when short sequences were included and used LwMo_Lys1_m and LwMoLys1_Lys2_m as the anchors. The obtained topology of subtrees for Ly s1 and Ly s2 was consistent with the SPDD rule. Thus, we decided to use these subtrees as the final subtrees for Ly. Ly subfamily 2 was diversified into two sub-subfamilies (s2.1 and s2.2) in a very early ancestor of Ly (Fig T2S4). Thus, we cannot exclude the possibility that the two sub-subfamilies were diversified on the main lineage before divergence of Ly and that one of the sub-subfamilies was lost while the ancestor of the other sub-subfamily was maintained on the main CBP60 lineage.

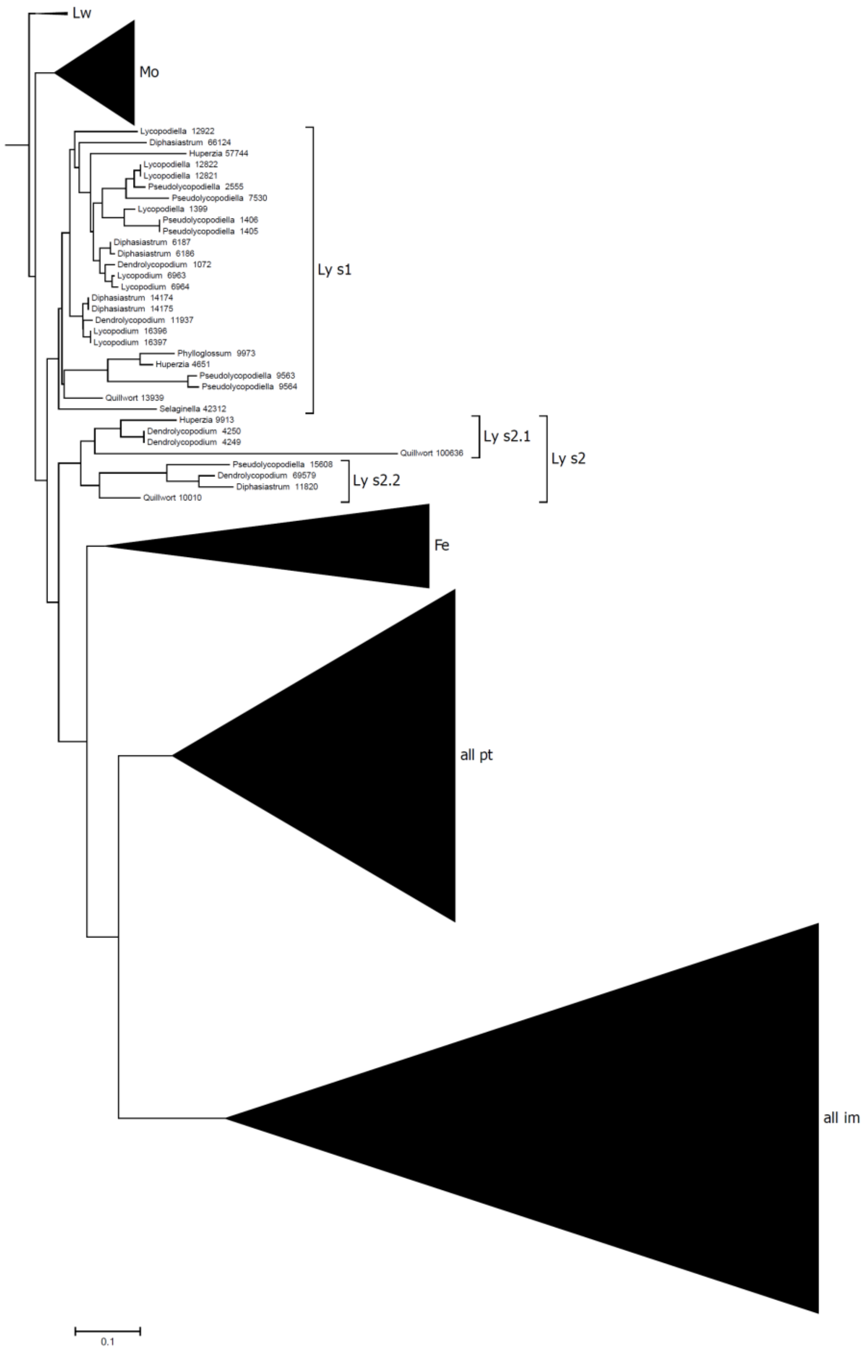


Fig T2S4 Duplication on the main lineage resulted in Ly subfamilies 1 and 2 (Ly s1 and Ly s2). Only the ancestor of Ly s2 was inherited in the main CBP60 lineage. Ly s2 diversified into two sub-subfamilies (Ly s2.1 and Ly s2.2). The clades other than those for Ly are compressed.

Three Lw sequences and Lw_Mo_m were used to infer an Lw subtree. 114 Mo sequences, Lw_Mo_m, and LwMo_Lys1_m were used to infer an Mo subtree. The Mo sequences were monophyletic even though the members of the “strange” clade and the related short sequences were included. The Mo subtree topology can be explained by an early duplication that led to a subfamily containing these “strange” sequences in Peat moss and Rockmoss and by subsequent loss of this subfamily.

189 Gy sequences and Fe_m, all_pt_m, all_im_m, An_pt_m, and An_im_m were used to infer subtrees for the Gy prototypical and Gy immune-related subfamilies. The Gy immune-related subfamily was diversified into two sub-subfamilies early in evolution of the Gy group.

Inference of subtrees for An sequences was conducted in a complex manner since there are no MRCAs to anchor the subtrees after emergence of An, since the prototypical group and the immune-related clade diversified into two and three subfamilies, respectively, and since the SARD1 subfamily was not included in the final overview tree. An sequences were divided into two groups: 207 BaMCM (Basal Angiosperms, Magnoliids, Chloranthales, and Monocots) sequences and 386 Ed (Eudicots) sequences. Note that BaMCM is not monophyletic in respect to the land plant lineage. The BaMCM sequences were classified into CBP60ef, CBP60bcd, CBP60a, CBP60g, and SARD1 subfamilies as follows. (1) The 207 BaMCM sequences were classified into the CBP60ef subfamily (BaMCM_60ef), the CBP60bcd subfamily (BaMCM_60bcd), and the collective immune regulator subfamilies (BaMCM_im), based on a tree made with the BaMCM sequences, Fe_m, Gy_Pt_m, and Gy_im_m. Using the same tree, the most probable MRCA sequence of Gy and the BaMCM prototypical group (Gy_BaMCM_pt_m) was inferred. (2) The 111 BaMCM collective immune regulator sequences were classified into the CBP60a subfamily (BaMCM_60a) and the collective CBP60g/SARD1 subfamily, based on a subtree made with the BaMCM_im sequences and An_im_m. In this subtree the division between the CBP60g and SARD1 subfamilies was not clear when compared to the species phylogeny. (3) The 71 collective BaMCM CBP60g/SARD1 sequences were classified into the CBP60g subfamily (BaMCM_60g) and the SARD1 subfamily (BaMCM_SARD1), based on the tree with the collective BaMCM CBP60g/SARD1 sequences and An_im_m. Since the SARD1 subfamily members have the C-terminal region deleted, 40% site coverage was used.

Since BaMCM was not monophyletic, the MRCAs of the Magnoliids/Chloranthales sequences (Mn_xx_m) and the MRCAs of the Monocot sequences (Mc_xx_m), each of which is monophyletic, were inferred from each BaMCM subfamily subtree made above. This yielded Mn_60a_m, Mc_60a_m, Mn_60g_m, Mc_60g_m, Mn_SARD1_m, and Mc_SARD1_m. In BaMCM_SARD1, diversification of sub-subfamilies (such as Sss1 in Text S1) occurred. The clades for the true Mn and Mc SARD1 subfamilies with high confidences were used for Mn_SARD1_m and Mc_SARD1_m.

A tree with the 76 BaMCM prototypical sequences and Gy_BaMCM_pt_m was used to infer the MRCAs for BaMCM CBP60bcd (BaMCM_60bcd_m), for Magnoliids/Chloranthales (Mn_60bcd_m), and for Monocot CBP60bcd (Mc_60bcd_m). Since most Basal Angiosperm and Monocot species did not have any CBP60ef subfamily sequences, similar MRCA inference (Mn_xx_m, Mc_xx_m) was not performed with the BaMCM CBP60ef sequences.

All 386 Ed (Eudicot) sequences were classified into the CBP60ef, CBP60bcd, CBP60a, CBP60g, and SARD1 subfamilies (Ed_60ef, Ed_60bcd, Ed_60a, Ed_60g, and Ed_SARD1), based on a tree with the Ed sequences, Fe_m, An_60ef_m, An_60bcd_m, An_pt_m, An_im_m, Mc_60a_m, Mn_60a_m, Mc_60g_m, Mn_60g_m, Mc_SARD1_m, and Mn_SARD1_m. Then for each subfamily, an Ed subtree was inferred. Specifically, 74 Ed SARD1 sequences, Mc_SARD1_m, and Mn_SARD1_m were used to infer the final Ed SARD1 tree and a most probable MRCA of the Ed SARD1 sequences (Ed_SARD1_m); 67 Ed CBP60g sequences, Mc_60g_m, and Mn_60g_m for the final Ed CBP60g tree and Ed_60g_m; 58 Ed CBP60a sequences, Mc_60a_m, and Mn_60a_m for the final Ed CBP60a tree and Ed_60a_m; 44 Ed CBP60ef sequences and An_60ef_m for the final Ed CBP60ef tree and Ed_60ef_m; 143 Ed CBP60bcd sequences, BaMCM_60bcd_m, Mc_60bcd_m, and Mn_60bcd_m for the final Ed CBP60bcd tree and Ed_60bcd_m.

The CBP60bcd subfamily diversified into two sub-subfamilies among Core Eudicots after divergence of Basal Eudicots, such as Blue Columbine and Sacred Lotus. While most Core Eudicot species have both sub-subfamilies of CBP60 bcd, the Brassicaceae family, which includes Arabidopsis, has lost one of the sub-subfamilies. Thus, all three CBP60bcd members of Arabidopsis, AtCBP60b, AtCBP60c, and AtCBP60d, belong to the other sub-subfamily. Although the “strange” clade Eucalyptus sequences and related short Eucalyptus sequences belonged to the sub-subfamily that included AtCBP60bcd members, this sequence group consisting of 10 Eucalyptus sequences had very long branch lengths and was placed close to the root of the sub-subfamily. Therefore, these 10 “strange” Eucalyptus sequences do not strictly follow the SPDD rule. We think that this group of sequences has been evolving very fast recently, and its phylogenetic relationships to other sequences inferred based on standard evolution models are not very accurate.

With Ed_xx_m MRCA sequences, the final BaMCM subfamily trees were inferred: 30 BaMCM SARD1 sequences, An_im_m, Ed_SARD1_m, Ed_60g_m, Ed_60a_m, Mc_60a_m, Mn_60a_m, Mc_60g_m, and Mn_60g_m were used to infer the final BaMCM SARD1 tree (containing Ed_SARD1_m) and the MRCA for all An SARD1 sequences (An_SARD1_bu_m); 41 BaMCM CBP60g sequences, An_im_m, Ed_SARD1_m, Ed_60g_m, Ed_60a_m, Mc_60a_m, Mn_60a_m, Mc_SARD1_m, and Mn_SARD1_m for the final BaMCM CBP60g tree (containing Ed_CBP60g_m) and An_60g_bu_m; 40 BaMCM CBP60a sequences, An_im_m, Ed_SARD1_m, Ed_60g_m, Ed_60a_m, Mc_60g_m, Mn_60g_m, Mc_SARD1_m, and Mn_SARD1_m for the final BaMCM CBP60a tree (containing Ed_CBP60a_m) and An_60a_bu_m; 15 BaMCM CBP60ef sequences, An_pt_m, and Ed_60ef_m for the final BaMCM CBP60ef tree (containing Ed_60ef_m) and An_60ef_bu_m; 81 BaMCM CBP60bcd sequences, An_pt_m, Ed_60bcd_m, Ed_60ef_m, and An_60ef_bu_m for the final BaMCM CBP60bcd tree (containing Ed_CBP60bcd_m) and An_60bcd_bu_m.

In the final step of MACCTI, the trees that were inferred for relationships within the clades were combined to reconstruct the entire tree with all of the 1024 CBP60 sequences. A backbone tree was inferred with the MRCAs for the clades, Lw_m, Mo_m, Ly_s1_m, Ly_s2_m, Fe_m,

Gy_pt_m, Gy_im_pt, An_60bcd_bu_m, An_60ef_bu_m, An_60a_bu_m, An_60g_bu_m, and An_SARD1_bu_m (Fig T2S5).

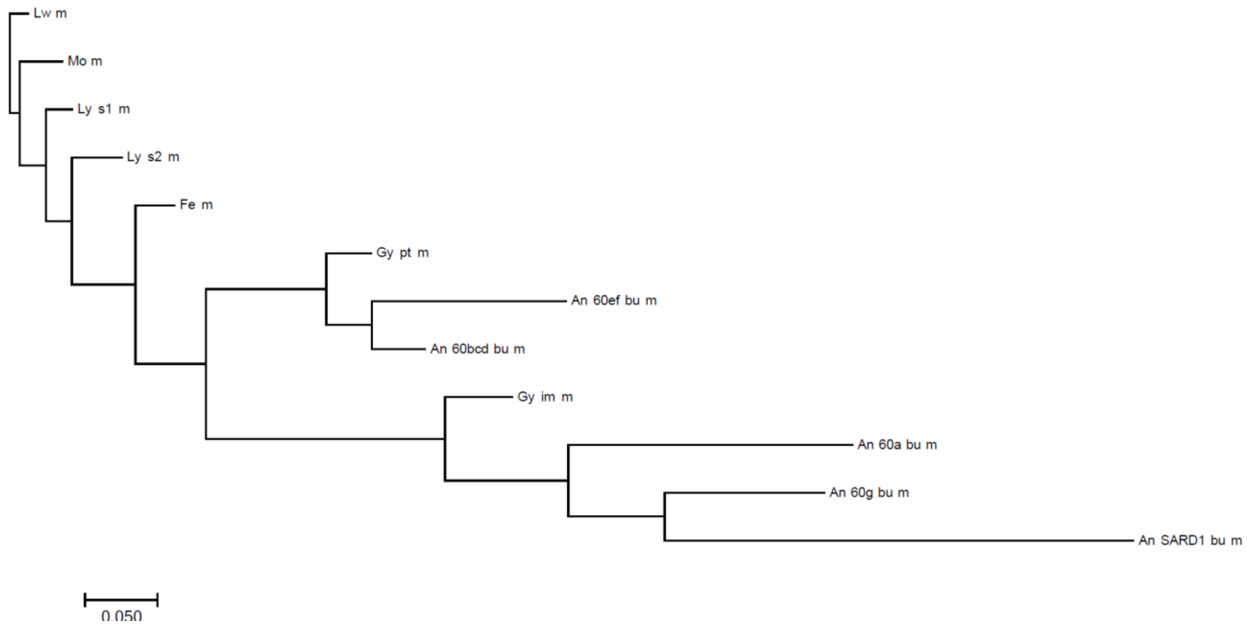


Fig T2S5. The backbone tree among the MRCAs.

The final tree for each of the SARD1, CBP60g, CBP60a, CBP60ef, and CBP60bcd subfamilies in An was reconstructed by replacing the Ed_xx_m in the final BaMCM subfamily trees with the corresponding final Ed subfamily trees: the final An SARD1 tree was reconstructed from the Ed SARD1 tree and the BaMCM SARD1 tree; the final An CBP60g tree from the Ed CBP60g and BaMCM CBP60g trees; the final An CBP60a tree from the Ed CBP60a and BaMCM CBP60a trees; the final An CBP60ef tree from the Ed CBP60ef and BaMCM CBP60ef trees; the final An CBP60bcd tree from the Ed CBP60bcd and BaMCM CBP60bcd trees.

The MRCAs in the backbone tree (Fig T2S5) were replaced with the corresponding clade trees to obtain the entire CBP60 tree consisting of the 1024 QC-ed sequences.

### Reordering of the sequence leaves in the CBP60 sequence tree

The sequence leaves of the MACCTI-obtained CBP60 tree were flipped at the branch points to order the prototypical group and the immune-related clade first, and to order the An subfamilies in the order of the non-Angiosperm clades, the CBP60ef, CBP60bcd, CBP60a, CBP60g, and SARD1 subfamilies second. Then within these groups, clades, and subfamilies, the correlation of the order of the species sources of the sequences to the order in the species phylogeny was maximized by flipping at the branch points. The tree obtained after the reordering procedure was designated the final tree of the CBP60 sequence phylogeny and shown in Fig 2 and Fig S1. The purpose of the reordering of the sequence leaves was to make evaluation of the tree topology according to the SPDD rule easy. For example, diversification into two subfamilies within each of the Fern sequence clade and the Gymnosperm immune-related clade can be visually easily recognized in a low-resolution tree without sequence names (Fig 2).

### Power of the MACCTI approach

It should be noted that the MACCTI approach resulted in placing of the “strange” clade sequences in the tree generally consistent with SPDD rule. Success in the placement of the “strange” Eucalyptus sequences was limited as the sequences were highly diverse from any other CBP60 sequences. The “strange” Eucalyptus sequences were still placed within one of the two Eudicot CBP60bcd sub-subfamilies, which suggests that their ancestral sequence originated from the CBP60bcd sub-subfamily.

Major challenges with phylogenetic inference using the full-length CBP60 sequences were that the large C-terminal region can strongly bias the inference due to poor alignability across different clades, including multiple independent clades lacking the C-terminal region and at the same time the phylogenetic information carried by the well-alignable CBP60-conserved domain is limited and inaccurate. By focusing on a single clade and very closely related MRCAs on the main CBP60 lineage, MACCTI allows a longer and more accurate multiple sequence alignment in general. In addition, the types of variation in the sequences in a clade are limited, which reduces the probability that different types of highly variable sequences artifactually get grouped into a single clade (e.g., the “strange” clade).

In MACCTI, multiple anchoring MRCAs that define different directions on the main CBP60 lineage from the divergence point of the clade in question were used, which gave sensitivity in inferring the topology with respect to the MRCAs. For example, for the Fe sequences, LyMoLys1_Lys2_m, all_pt_m, and all_im_m were used as the anchoring MRCAs. In this way, the questions of whether the Fe sequences were monophyletic and whether the Fe clade diversified before or after diversification of the immune-related clade from the prototypical group were tested with high sensitivity. Since the MRCAs were anchored in the backbone tree, the local features of the clade-by-clade trees can be translated into the context of the entire tree.

We also noticed that it was often important to include MRCAs that represent recent diversifications in order to obtain a more accurate tree. For example, to accurately infer the BaMCM CBP60bcd tree, it was important to include Ed_60ef_m and An_60ef_bu_m. We sometimes tried a sequence set including different MRCAs to find a good set of anchoring MRCAs for more accurate tree inference judged by the SPDD rule.

### Fast evolution of some specific CBP60 clades

In addition to the immune regulator CBP60a, CBP60g, and SARD1 subfamilies, some small CBP60 clades or individual sequences have been evolving very fast. The “strange” sequences and short sequences related to them in the final CBP60 sequence phylogeny are clear examples.

In Text S1, we also described as another fast evolving clade the “distant homolog” clade in Angiosperm species, which appeared to share an ancestor with the CBP60g subfamily. The “distant homolog” sequences were mostly omitted from our CBP60 sequence set. Some such sequences, such as “Floating_fern_s0021.g008453”, may be encoded by pseudogenes or pseudogenizing genes as they are single sequences in single species. However, other sequences had multiple sequences in multiple species and are not likely to be encoded by pseudogenes or pseudogenizing genes, e.g., a Moss subfamily consisting of the “strange” and related sequences in Peat moss and Rockmoss and the “distant homolog” clade sequences. AtCBP60g and AtSARD1 are sequence-specific DNA-binding proteins [25,27]. It is likely that other CBP60 proteins are also sequence-specific DNA-binding proteins. It is possible that these fast evolving sequences may have acquired different sequence-specificity and target genes, i.e., neofunctionalization.

**Distance-based ortholog inference is hampered by unbalanced evolution rates across clades**

Unbalanced evolution rates across the subfamilies, such as prototypical group and immune regulator subfamilies in the CBP60 family, presents practical challenges in assigning sequences to subfamilies based on sequence similarity. For example, Fig T2S6A shows a heatmap of the relative BLASTP bit scores among Arabidopsis CBP60 members when they were queried with each of the Rice CBP60 members. When Rice CBP60a or CBP60g was used as a query, some of the prototypical Arabidopsis CBP60 members, AtCBP60b-f, have clearly higher bit scores than the immune regulators AtCBP60a, g, or AtSARD1. Therefore, the orthologous relationships between Arabidopsis and Rice CBP60a or CBP60g cannot be discovered solely based on the sequence similarity between two species. This problematic situation between, for example, Arabidopsis and Rice CBP60g orthologs is explained as follows (Fig T2S6B): Since CBP60g has been evolving very fast, the dissimilarity between Arabidopsis CBP60g and the MRCA of Arabidopsis and Rice CBP60g ($D_{gAM}$) and the dissimilarity between Rice CBP60g and the MRCA ($D_{gRM}$) are very high; On the other hand, the dissimilarity between the MRCA and some of the prototypical Arabidopsis CBP60 ($D_{gMpA}$) is relatively low; Consequently, the dissimilarity between Arabidopsis and Rice CBP60g ($D_{gAM}$ + $D_{gRM}$) is higher than the dissimilarity between Rice CBP60g and an Arabidopsis prototypical CBP60 ($D_{gRM}$ + $D_{gMpA}$). It is a common practice to use the reciprocal BLAST top hits between gene sequences in two species for the purpose of ortholog identification at the genomic scale [29]. It has been pointed out that unbalanced evolution rates within a subfamily could cause a problem in the similarity-based orthology identification [30]. The CBP60 case showed that unbalanced evolution rates across the subfamilies also cause a problem.

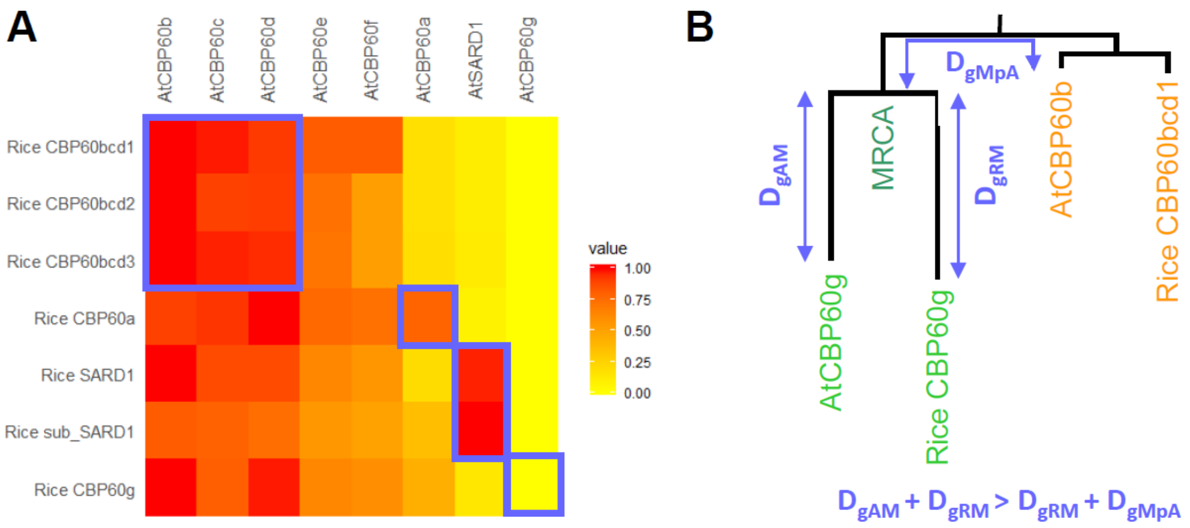


Fig T2S6. Sequence similarity alone cannot be used to identify the subfamily memberships in the CBP60 family. (A) The relative bit score in BLASTP using each of the Rice CBP60 members (rows) as a query and the Arabidopsis CBP60 members (columns) as the subjects. The bit scores were linearly scaled from 0 to 1 (yellow to red) for each query (row). Blue boxes indicate the corresponding subfamily members between Rice and Arabidopsis. Note that with Rice CBP60a and CBP60g as queries, the highest bit scores do not correspond to the same subfamily members in Arabidopsis. (B) A depiction of how unbalanced evolution rates affect subfamily membership identification based on a dissimilarity measure. It shows that Rice CBP60g is more similar to AtCBP60b than to AtCBP60g.

### Inference of the SARD1 divergence point

The topology of our CBP60 sequence phylogenetic tree suggested that the three immune regulator CBP60a, CBP60g, and SARD1 subfamilies diversified at or immediately before divergence of Angiosperms. While the CBP60a and CBP60g subfamilies had Basal Angiosperm sequences, including Amborella sequences, the SARD1 subfamily did not have any Basal Angiosperm sequences. To exclude the possibility that our BLASTP bit score cutoff of FMS was too high to detect Basal Angiosperm SARD1 sequences, we manually scrutinized the BLASTP hit results for much lower bit scores but did not find any SARD1 sequences. We also searched the Amborella transcriptome from OneKP [5] but did not find any SARD1 sequences. There are two possible explanations for the observations of no Basal Angiosperm SARD1 sequences: (1) the inferred tree topology is incorrect, and the SARD1 subfamily diversified from the CBP60g subfamily after divergence of Basal Angiosperms; (2) the inferred tree topology is correct - the SARD1 lineage was deleted in Basal Angiosperms and/or SARD1 subfamily members are present in Basal Angiosperms but were missed in the Basal Angiosperm databases. There are two issues regarding explanation (2). First, the six Basal Angiosperm species we had for our sequence databases (Amborella, Austrobaileya, Anisetree, Carib, Pond_lily, and Water_lily) are not monophyletic with respect to the land plant lineage [28]. Loss of the SARD1 lineage in all five Basal Angiosperm species would require at least three independent gene deletion events.

Second, on the other hand, the sequence databases for all Basal Angiosperm species except Amborella were based on transcriptome sequencing. Particular sequences existing in the genomes could be missed in transcriptome databases if the transcript levels of the sequences in question were low in the tissues used for the transcriptome libraries. Thus, we thought that explanation (2) is possible considering both issues.

We compared the likelihoods of the tree topologies for the two explanations. To obtain a better resolution around the divergence points of immune subfamilies, we compiled a sequence list of all Basal Angiosperm, Magnoliids and Chlorantales (BaMCh) CBP60 sequences and used Liverwort_0018s0017 as an outgroup. The sequences were aligned using PRANK [31], a probabilistic multiple alignment program. PRANK aims to reduce over-estimation of deletion events in favor of insertion events, which could produce a more accurate tree. The resulting alignment was used in RAxML [24] (default parameter value set) to infer a ML-based phylogenetic tree. The resulting tree topology still showed that the SARD1 subfamily diversified at or before divergence of Angiosperms, supporting explanation (2). We created an alternative tree topology in agreement with explanation (1) for this dataset using TreeGraph [32]. The two tree structures were compared in a likelihood ratio test using RAxML, and the tree structure for explanation (2) was favored ($p < 0.01$; Figs S2A and S2B).

Recently, a large-scale waterlily genome research paper was published [28]. We downloaded protein sequences from 19 species in the order Nymphaeales. One of those species, *Nymphaea colorata*, was genome sequenced. We searched with BLASTP using the Liverwort CBP60 conserved domain as a query against these newly sequenced genomes. The same QC criteria were used as described above. Proteins that passed QC were added to our Basal Angiosperm/Magnoliids/Chlorantales dataset, with Liverwort_0018s0017 as an outgroup. Sequences were aligned by PRANK [31], a tree was inferred by RaXML [24] with default parameters and visualized by iTol [20]. Using this new dataset we have identified multiple SARD1 sequences in Nymphaeales, strongly supporting explanation (2). Fig S2C shows a phylogenetic tree of the BaMCh sequence dataset supplemented with CBP60 sequences of two waterlily species *Nuphar advena* and *Nymphaea colorata*.

### Calmodulin binding prediction

Predicting calmodulin binding sites in proteins is notoriously challenging. Although there are many known calmodulin binding motifs, calmodulin binding is not predictable based on the motifs. Secondary structure seems to be important, for example most calmodulin binding domains seem to contain a basic amphiphathic alpha helix [33]. Recognizing these difficulties, we have tried different calmodulin binding prediction algorithms and decided to use CaMELS [26] to analyze protein sequences in our final dataset. CaMELS is a machine-learning based algorithm and was the only calmodulin binding prediction algorithm among those we tested that accurately identified C-CaMBD and N-CaMBD in AtCBP60a and AtCBP60g, respectively, which have been previously experimentally identified [34,35]. The output of this prediction is a calmodulin binding score, which shows the propensity of proteins to interact with CaM on a scale from 0 to 100. CaMELS also identifies the position in the sequence (in a sliding window manner) where the CaM-binding prediction score is maximal. Besides AtCBP60g, only one other protein in our dataset had a predicted N-terminal binding site with a high CaM-binding prediction score (>75.0); Cacao_011988t1 from the CBP60a subfamily. We did not assess the accuracy of CaM binding site predictions by CaMELS beyond those previously investigated experimentally.

We have further used the DECIPHER package in R [36] to predict alpha-helical propensity of CBP60 protein C-termini. Additionally, A Python script, hydrophobic_moment.py (https://gist.github.com/JoaoRodrigues/568c845915aea3efa3578babfd72423c), was used to calculate hydrophobic moment at C-termini, as a measure of amphipathicity.

#### Twelve Core Eudicot Species and their CBP60 sequences

First, the CBP60 sequences for Eudicot species were extracted from the QC-ed full set of CBP60 sequences. Second, they were classified into the CBP60bcd, CBP60a, CBP60g, and SARD1 subfamilies according to the CBP60 sequence phylogenetic analysis results shown in Fig 2. The CBP60ef subfamily members were removed because many Superasterid species did not have CBP60ef subfamily members. Third, the CBP60-conserved domain of Liverwort_ Mapoly0018s0017.1 (sites 88-380; 293 aa) was used as the query in BLASTP against the database consisting of the Eudicot CBP60 sequences, and the sequences with query alignment length < 225 aa were removed. If the part N-terminal to the alignment in a subject was longer than 100 aa, the N-terminus of the sequence was trimmed down to 100 aa. If the part C-terminal to the alignment in a subject was longer than 350 aa, the C-terminus of the sequence was trimmed down to 350 aa. The resulting sequence set was divided into each of the 4 subfamilies. Fourth, the CBP60-conserved domain sequences for Eudicot CBP60 members in each subfamily were defined based on the multiple sequence alignment together with the CBP60-conserved domain of Liverwort_ Mapoly0018s0017.1 by ClustalW (default parameter values). In this way the Eudicot CBP60-conserved domain sequence set for four subfamilies was made. Note that multiple sequence alignment gives a better domain definition particularly near the boundaries of the domain than pairwise sequence alignment. Fifth, the CBP60-conserved domain sequences for each subfamily for twelve Core Eudicot species Quinoa, Coffee, Tomato, Monkeyflower, Carrot, Sunflower, Poplar, Soybean, Peach, Orange, Cotton, and Arabidopsis were extracted (total 100 sequences). Each of the species represents a different taxonomic order, and the former 6 species and the latter 6 species belong to Superasterids and Superrosids, respectively (Fig S3). Sixth, the extracted 100 sequences were used to infer a phylogenetic tree, using ClustalW and ML in MEGA 7 with default parameter values. There were two sub-subfamilies within the CBP60bcd subfamily. One of them contained AtCBP60bcd while the other did not contain any Arabidopsis CBP60. The sub-subfamily with AtCBP60bcd was chosen to represent the CBP60bcd subfamily. When a species had more than one CBP60 sequence for a subfamily, the sequence that had the shortest branch length in the above tree was selected to make a sequence set for four subfamilies for the 12 Core Eudicot species used in the PEAES-PDR analysis (set 1; total 48 sequences). An alternative set was made by selecting the sequence that had the longest branch length, which is called an “alternative set” in the text (set 2; total 48 sequences). The sequence names for each set are as follows.

Set 1

CBP60a: Quinoa_62031038; Coffee_04g13430, Tomato_03g113920.2.1, Monkeyflower_D01792, Carrot_026115, Sunflower_9g00013686, Poplar_015G071800.1, Soybean_05G034600, Peach_5G178300, Orange_1g008453m, Cotton_003G109800, Arabidopsis_AT5G62570.2

CBP60g: Quinoa_62000695, Coffee_02g34690, Tomato_01g100240.2.1, Monkeyflower_B00990, Carrot_025268, Sunflower_16g00021305, Poplar_013G010700.1, Soybean_10G148700, Peach_6G315700, Orange_1g008593m, Cotton_013G128100, Arabidopsis_AT5G26920.1

SARD1: Quinoa_62033803, Coffee_04g06970, Tomato_03g119250.2.1, Monkeyflower_I00405, Carrot_023217, Sunflower_16g00022942, Poplar_015G045300.1, Soybean_09G182400, Peach_5G223600, Orange_1g011961m, Cotton_006G059900, Arabidopsis_AT1G73805.1

CBP60bcd: Quinoa_62040307, Coffee_06g13450, Tomato_07g006830.2.1, Monkeyflower_K00253, Carrot_009913, Sunflower_5g00005047, Poplar_018G095300.1, Soybean_17G065500, Peach_1G089100, Orange_1g006542m, Cotton_013G246400, Arabidopsis_AT4G25800.1

Set 2

CBP60a: Quinoa_62008278, Coffee_04g13430, Tomato_03g113920.2.1, Monkeyflower_I00050, Carrot_016504, Sunflower_2g00026594, Poplar_012G077000.1, Soybean_17G092700, Peach_5G178300, Orange_1g008453m, Cotton_008G297800, Arabidopsis_AT5G62570.2

CBP60g: Quinoa_62005263, Coffee_02g34690, Tomato_01g100240.2.1, Monkeyflower_N02112, Carrot_025268, Sunflower_16g00021849, Poplar_005G020200.1, Soybean_03G232400, Peach_6G315700, Orange_1g008593m, Cotton_004G237500, Arabidopsis_AT5G26920.1

SARD1: Quinoa_62029901, Coffee_04g06970, Tomato_12g036390.1.1, Monkeyflower_I00405, Carrot_023217, Sunflower_17g00009729, Poplar_012G054900.1, Soybean_08G044400, Peach_5G223600, Orange_1g011961m, Cotton_008G287500, Arabidopsis_AT1G73805.1

CBP60bcd: Quinoa_62040001, Coffee_06g13450, Tomato_07g006830.2.1, Monkeyflower_J01456, Carrot_009913, Sunflower_5g00005047, Poplar_018G095300.1, Soybean_13G094800, Peach_1G089100, Orange_1g006542m, Cotton_004G031000, Arabidopsis_AT2G18750.1

For each of sets 1 and 2, multiple sequence alignment of the CBP60-conserved domains was performed by ClustalW (MEGA 7, default parameter values) with a guide tree whose topology was generated according to the species phylogeny and the subfamily phylogeny. Then we manually edited the multiple sequence alignments. The objectives of manual editing were to remove insertional polymorphisms that are specific to one or two sequences to make the total alignment length across the four subfamilies short (i.e., fewest gap sites) and to make the alignment consistent between the two sets. The following deletions were made in each set. The site position number for each sequence is according to the CBP60-conserved domain sequence, not the original sequence that contained sequences outside the CBP60-conserved domain.

Set 1

i. SARD1_Quinoa_62033803, #N261, #G210-#K214, #Y99
ii. CBP60g_Coffee_02g34690, #K206
iii. CBP60bcd_Sunflower_5g00005047, #N205-#K210
iv. SARD1_Tomato_03g119250.2.1, #I219, #T66
v. SARD1_Coffee_04g06970, #A131-#A132
vi. SARD1_Peach_5G223600, #A137, #S48-#D49, #N39-#Q42
vii. CBP60g_Arabidopsis_AT5G26920.1, #T100
viii. SARD1_Poplar_015G045300.1, #C91
ix. CBP60g_Monkeyflower_B00990, #Q63
x. SARD1_Sunflower_16g00022942, #V60
xi. SARD1_Soybean_09G182400, #D65
xii. SARD1_Orange_1g011961m, #I40

Set 2

i. CBP60g_Coffee_02g34690, #K206
ii. SARD1_Quinoa_62029901, #N258, #E210-#K211, #C99
iii. CBP60bcd_Sunflower_5g00005047, #N205-#K210
iv. CBP60g_Soybean_03G232400, #L161-#K162
v. SARD1_Tomato_12g036390.1.1, #I127, #Y26
vi. SARD1_Coffee_04g06970, #A131-A132
vii. SARD1_Peach_5G223600, #A137, #S48-#D49, #N39-#Q42
viii. CBP60g_Arabidopsis_AT5G26920.1, #T100
ix. SARD1_Poplar_012G054900.1, #C91
x. SARD1_Soybean_08G044400, #Y71-#S72, #A63-#P64, #V40-#A41
xi. SARD1_Orange_1g011961m, #I40

The resulting edited multiple sequence alignments (296 sites) were used in the PEAES-PDR analysis. The FASTA file of the multiple sequence alignments for Sets 1 and 2 are provided as supplemental files.

### PEAES metric

PEAES is a way to describe the physical-chemical characteristics of 20 proteinogenic amino acids and gap in a 6-dimensional Euclidean space. The dissimilarity between two amino acids (or between an amino acid and a gap) is defined by the Euclidean distance between them in the space. The 5-dimensional coordinates for 20 amino acids were according to Venkatarajan and Braun [37], which were obtained by linear dimensionality reduction from 237 physical-chemical property dimensions to 5 dimensions. Major determinants of the 1st, 2nd, and 3rd dimension values are the hydrophilicity, side chain length, and alpha-helix frequency, respectively. We added the 6th dimension, which is a "gap" dimension: value 1 for the gap and the value 0 for all amino acids were assigned. In this way, the distance between an amino acid and a gap is close to the maximum pairwise distance value, while the distances among the amino acids are kept unchanged from those in Venkatarajan and Braun. The 6-dimensional coordinates for the amino acids and gap are in Table S3, and their pairwise distances are in Table S4.

**PEAES-PDR analysis**

The Pairwise Distance Rank (PDR) was calculated using the PEAES metric (Fig 5). The PDR was determined for each site of a multiple sequence alignment in comparison of two protein sequence groups. Specifically, the multiple sequence alignment is that for one of Sets 1 and 2 for 12 Core Eudicot species, and the protein sequence groups are two CBP60 subfamilies in the set. The following procedure was applied to each site of the alignment. First, all pairwise distances were determined between every species sequence in group 1 and every species sequence in group 2 according to the PEAES pairwise distance (Table S4). Second, for each species (species A), the rank of the distance between species A in group 1 and species A in group 2 relative to the distances between species A in group 1 and every permuted species in group 2 and the distance between species A in group 2 and every permuted species in group 1 was determined. This is the rank determined among 23 distance values. The rank is the PDR for species A at the site in comparison of groups 1 and 2. We calculated the PDR for every site in every species in comparisons of (i) CBP60a and CBP60g subfamilies, (ii) CBP60g and SARD1 subfamilies, or (iii) SARD1 and CBP60a subfamilies, as groups 1 and 2. We indicate the PDRs for comparisons of these different group pairs as CBP60a:CBP60g, CBP60g:SARD1, and SARD1:CBP60a, respectively. We used only the sites with (number of variants in group 1) * (number of variants in group 2) $\geq$ 12 in each group pair to ascertain a reasonable level of diversity in the group pair comparison.

The median rank among 23 values is 12, and this is the expected mean rank value if evolution of two subfamily protein sequences is independent. Thus, a significant coevolution interaction can be detected if the PDR value is significantly different from the expected mean rank value. However, we cannot know the null distribution of the PDR value since only certain amino acids are possible at a particular site for each group to maintain the function of the group protein and since how these amino acids would distribute at the site under neutral evolution is unknowable. Furthermore, there could be some effect of phylogenic relationships within the group although we initially assumed and later proved that this effect is relatively small compared to the effect of selection.

We instead tested non-randomness in the distribution of PDRs between two group pairs for each species, at the sites where both group pairs have PDR values. We derived the coevolution interaction hypotheses (Hypotheses 3 and 4) from our coevolution mechanism hypotheses (Hypotheses 1 and 2). Hypotheses 3 and 4 predict that when PDR values between SARD1:CBP60a and CBP60g:SARD1 (SARD1-common comparison) at every site are plotted for each species (Fig S7B) and the plot is divided at the median rank value of 12 on each axis, the site points are enriched in quadrants 1 and 3 over quadrants 2 and 4. The designations of the quadrants are shown in Fig 6A. The hypotheses also predict that when PDR values between CBP60g:SARD1 and CBP60a:CBP60g (CBP60g-common comparison) at every site are plotted for each species (Fig S7A), the site points are enriched in quadrants 1 and 3 over quadrants 2 and 4. On the other hand, the hypotheses do not predict anything about the site-point enrichment in particular quadrants when PDR values between CBP60a:CBP60g and SARD1:CBP60a (CBP60a-common comparison) at every site are plotted for each species (Fig S7C). The *p*-value for the site-point enrichment in quadrants 1 and 3 vs. quadrants 2 and 4 was calculated by applying Fisher's exact test (2-sided) to the 2x2 contingency table of the site-point numbers for the four quadrants. The obtained *p*-values were corrected using the Benjamini-Hochberg FDR for each comparison. When the corrected *p*-value was smaller than 0.05, the first principal component (PC1) was shown in the plot to indicate the significant enrichment in quadrants 1 and 3 over quadrants 2 and 4 (i.e., PC1 has a positive slope).

### REFERENCES


1. Li, F.W., Brouwer, P., Carretero-Paulet, L., Cheng, S., De Vries, J., Delaux, P.M., Eily, A., Koppers, N., Kuo, L.Y., Li, Z., *et al.* (2018). Fern genomes elucidate land plant evolution and cyanobacterial symbioses. Nat. Plants.

2. Yagi, M., Kosugi, S., Hirakawa, H., Ohmiya, A., Tanase, K., Harada, T., Kishimoto, K., Nakayama, M., Ichimura, K., Onozaki, T., *et al.* (2014). Sequence analysis of the genome of carnation (Dianthus caryophyllus L.). DNA Res.

3. Al-Dous, E.K., George, B., Al-Mahmoud, M.E., Al-Jaber, M.Y., Wang, H., Salameh, Y.M., Al-Azwani, E.K., Chaluvadi, S., Pontaroli, A.C., Debarry, J., *et al.* (2011). De novo genome sequencing and comparative genomics of date palm (Phoenix dactylifera). Nat. Biotechnol.

4. Goodstein, D.M., Shu, S., Howson, R., Neupane, R., Hayes, R.D., Fazo, J., Mitros, T., Dirks, W., Hellsten, U., Putnam, N., *et al.* (2012). Phytozome: A comparative platform for green plant genomics. Nucleic Acids Res.

5. Leebens-Mack, J.H., Barker, M.S., Carpenter, E.J., Deyholos, M.K., Gitzendanner, M.A., Graham, S.W., Grosse, I., Li, Z., Melkonian, M., Mirarab, S., *et al.* (2019). One thousand plant transcriptomes and the phylogenomics of green plants. Nature.

6. Zheng, Y., Wu, S., Bai, Y., Sun, H., Jiao, C., Guo, S., Zhao, K., Blanca, J., Zhang, Z., Huang, S., *et al.* (2019). Cucurbit Genomics Database (CuGenDB): A central portal for comparative and functional genomics of cucurbit crops. Nucleic Acids Res.

7. Proost, S., Bel, M. Van, Vaneechoutte, D., Van De Peer, Y., Inzé, D., Mueller-Roeber, B., and Vandepoele, K. (2015). PLAZA 3.0: An access point for plant comparative genomics.

Nucleic Acids Res.

8. Gonzales, M.D., Archuleta, E., Farmer, A., Gajendran, K., Grant, D., Shoemaker, R., Beavis, W.D., and Waugh, M.E. (2005). The Legume Information System (LIS): An integrated information resource for comparative legume biology. Nucleic Acids Res.

9. Sato, S., Hirakawa, H., Isobe, S., Fukai, E., Watanabe, A., Kato, M., Kawashima, K., Minami, C., Muraki, A., Nakazaki, N., *et al.* (2011). Sequence analysis of the genome of an oil-bearing tree, jatropha curcas L. DNA Res.

10. Sato, S., Nakamura, Y., Kaneko, T., Asamizu, E., Kato, T., Nakao, M., Sasamoto, S., Watanabe, A., Ono, A., Kawashima, K., *et al.* (2008). Genome structure of the legume, Lotus japonicus. DNA Res.

11. Hiroyasu, K., Feng, L., Hideki, H., Takahiro, K., Zhongwei, Z., Yoichi, H., Kaoru, T., Sachiko, S., Aki, F., Shuji, Y., *et al.* (2014). Draft Sequences of the Radish (Raphanus sativus L.) Genome. DNA Res.

12. Bombarely, A., Rosli, H.G., Vrebalov, J., Moffett, P., Mueller, L.A., and Martin, G.B. (2012). A draft genome sequence of Nicotiana benthamiana to enhance molecular plant-microbe biology research. Mol. Plant-Microbe Interact.

13. van Bakel, H., Stout, J.M., Cote, A.G., Tallon, C.M., Sharpe, A.G., Hughes, T.R., and Page, J.E. (2011). The draft genome and transcriptome of Cannabis sativa. Genome Biol.

14. Singh, R., Ong-Abdullah, M., Low, E.T.L., Manaf, M.A.A., Rosli, R., Nookiah, R., Ooi, L.C.L., Ooi, S.E., Chan, K.L., Halim, M.A., *et al.* (2013). Oil palm genome sequence reveals divergence of interfertile species in Old and New worlds. Nature.

15. Ming, R., VanBuren, R., Liu, Y., Yang, M., Han, Y., Li, L.T., Zhang, Q., Kim, M.J., Schatz, M.C., Campbell, M., *et al.* (2013). Genome of the long-living sacred lotus (Nelumbo nucifera Gaertn.). Genome Biol.

16. Unver, T., Wu, Z., Sterck, L., Turktas, M., Lohaus, R., Li, Z., Yang, M., He, L., Deng, T., Escalante, F.J., *et al.* (2017). Genome of wild olive and the evolution of oil biosynthesis. Proc. Natl. Acad. Sci. U. S. A.

17. Eddy, S.R. (2011). Accelerated profile HMM searches. PLoS Comput. Biol.

18. Thompson, J.D., Higgins, D.G., and Gibson, T.J. (1994). CLUSTAL W: Improving the sensitivity of progressive multiple sequence alignment through sequence weighting, position-specific gap penalties and weight matrix choice. Nucleic Acids Res.

19. Guindon, S., and Gascuel, O. (2003). A Simple, Fast, and Accurate Algorithm to Estimate Large Phylogenies by Maximum Likelihood. Syst. Biol.

20. Letunic, I., and Bork, P. (2019). Interactive Tree Of Life (iTOL) v4: recent updates and new developments. Nucleic Acids Res.

21. Kumar, S., Stecher, G., and Tamura, K. (2016). MEGA7: Molecular Evolutionary Genetics Analysis Version 7.0 for Bigger Datasets. Mol. Biol. Evol.

22. Qiu, Y.L., Li, L., Wang, B., Chen, Z., Knoop, V., Groth-Malonek, M., Dombrovska, O.,

Lee, J., Kent, L., Rest, J., *et al.* (2006). The deepest divergences in land plants inferred from phylogenomic evidence. Proc. Natl. Acad. Sci. U. S. A.

23. Chase, M.W., Christenhusz, M.J.M., Fay, M.F., Byng, J.W., Judd, W.S., Soltis, D.E., Mabberley, D.J., Sennikov, A.N., Soltis, P.S., Stevens, P.F., *et al.* (2016). An update of the Angiosperm Phylogeny Group classification for the orders and families of flowering plants: APG IV. Bot. J. Linn. Soc.

24. Stamatakis, A. (2014). RAxML version 8: A tool for phylogenetic analysis and post-analysis of large phylogenies. Bioinformatics.

25. Zhang, Y., Xu, S., Ding, P., Wang, D., Cheng, Y.T., He, J., Gao, M., Xu, F., Li, Y., Zhu, Z., *et al.* (2010). Control of salicylic acid synthesis and systemic acquired resistance by two members of a plant-specific family of transcription factors. Proc. Natl. Acad. Sci. U. S. A.

26. Abbasi, W.A., Asif, A., Andleeb, S., and Minhas, F. ul A.A. (2017). CaMELS: In silico prediction of calmodulin binding proteins and their binding sites. Proteins Struct. Funct. Bioinforma.

27. Qin, J., Wang, K., Sun, L., Xing, H., Wang, S., Li, L., Chen, S., Guo, H.S., and Zhang, J. (2018). The plant-specific transcription factors CBP60G and SARD1 are targeted by a verticillium secretory protein VDSCP41 to modulate immunity. Elife.

28. Zhang, L., Chen, F., Zhang, X., Li, Z., Zhao, Y., Lohaus, R., Chang, X., Dong, W., Ho, S.Y.W., Liu, X., *et al.* (2020). The water lily genome and the early evolution of flowering plants. Nature.

29. Moreno-Hagelsieb, G., and Latimer, K. (2008). Choosing BLAST options for better detection of orthologs as reciprocal best hits. Bioinformatics.

30. Eisen, J.A., and Fraser, C.M. (2003). Phylogenomics: Intersection of evolution and genomics. Science (80-. ).

31. Löytynoja, A., and Goldman, N. (2010). WebPRANK: A phylogeny-aware multiple sequence aligner with interactive alignment browser. BMC Bioinformatics.

32. Stöver, B.C., and Müller, K.F. (2010). TreeGraph 2: Combining and visualizing evidence from different phylogenetic analyses. BMC Bioinformatics.

33. Yap, K.L., Kim, J., Truong, K., Sherman, M., Yuan, T., and Ikura, M. (2000). Calmodulin target database. J. Struct. Funct. Genomics.

34. Truman, W., Sreekanta, S., Lu, Y., Bethke, G., Tsuda, K., Katagiri, F., and Glazebrook, J. (2013). The CALMODULIN-BINDING PROTEIN60 family includes both negative and positive regulators of plant immunity. Plant Physiol. *163*, 1741–1751.

35. Wang, L., Tsuda, K., Sato, M., Cohen, J.D., Katagiri, F., and Glazebrook, J. (2009). Arabidopsis CaM binding protein CBP60g contributes to MAMP-induced SA accumulation and is involved in disease resistance against Pseudomonas syringae. PLoS Pathog. *5*, e1000301. Available at: https://dx.plos.org/10.1371/journal.ppat.1000301.

36. Wright, E.S. (2016). Using DECIPHER v2.0 to analyze big biological sequence data in R. R J.

37. Venkatarajan, M., and Braun, W. (2001). New quantitative descriptors of amino acids based on multidimensional scaling of a large number of physical-chemical properties. J. Mol. Model.
