## Supplementary figures and images for "Pathogen-driven coevolution across CBP60 plant immune regulator subfamilies confers resilience on the regulator module"

### FigS1.pdf

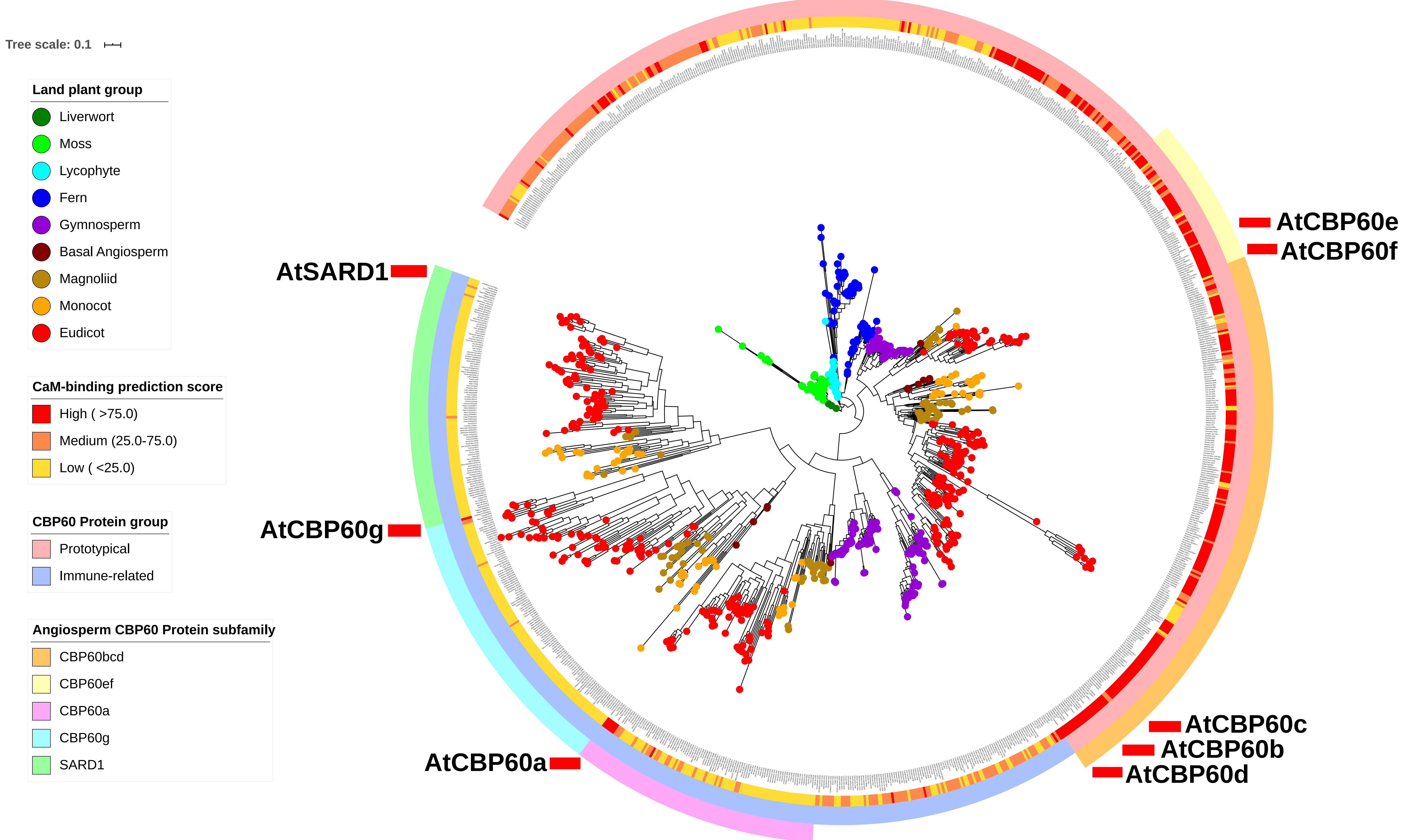

### FigS2.pdf

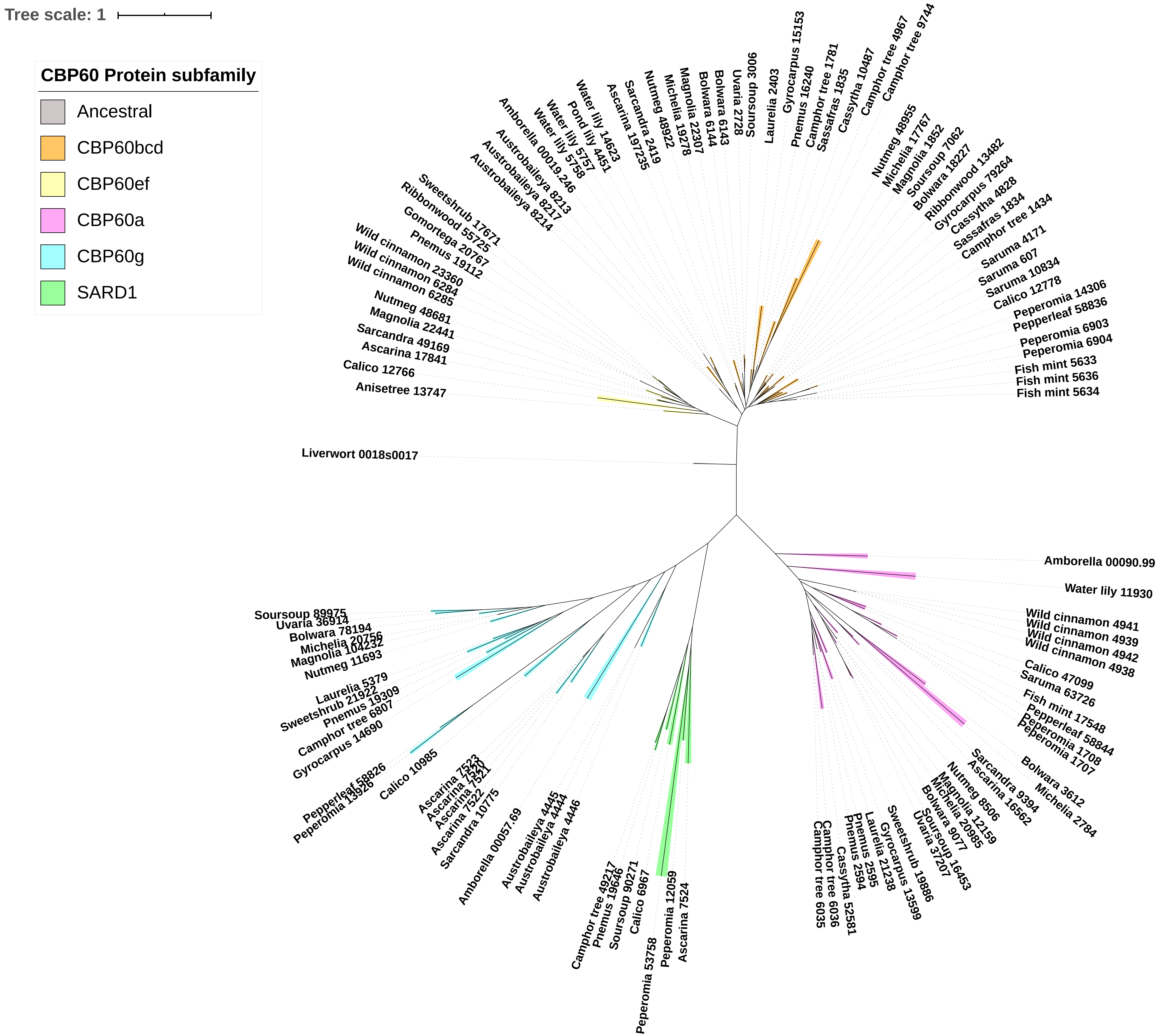

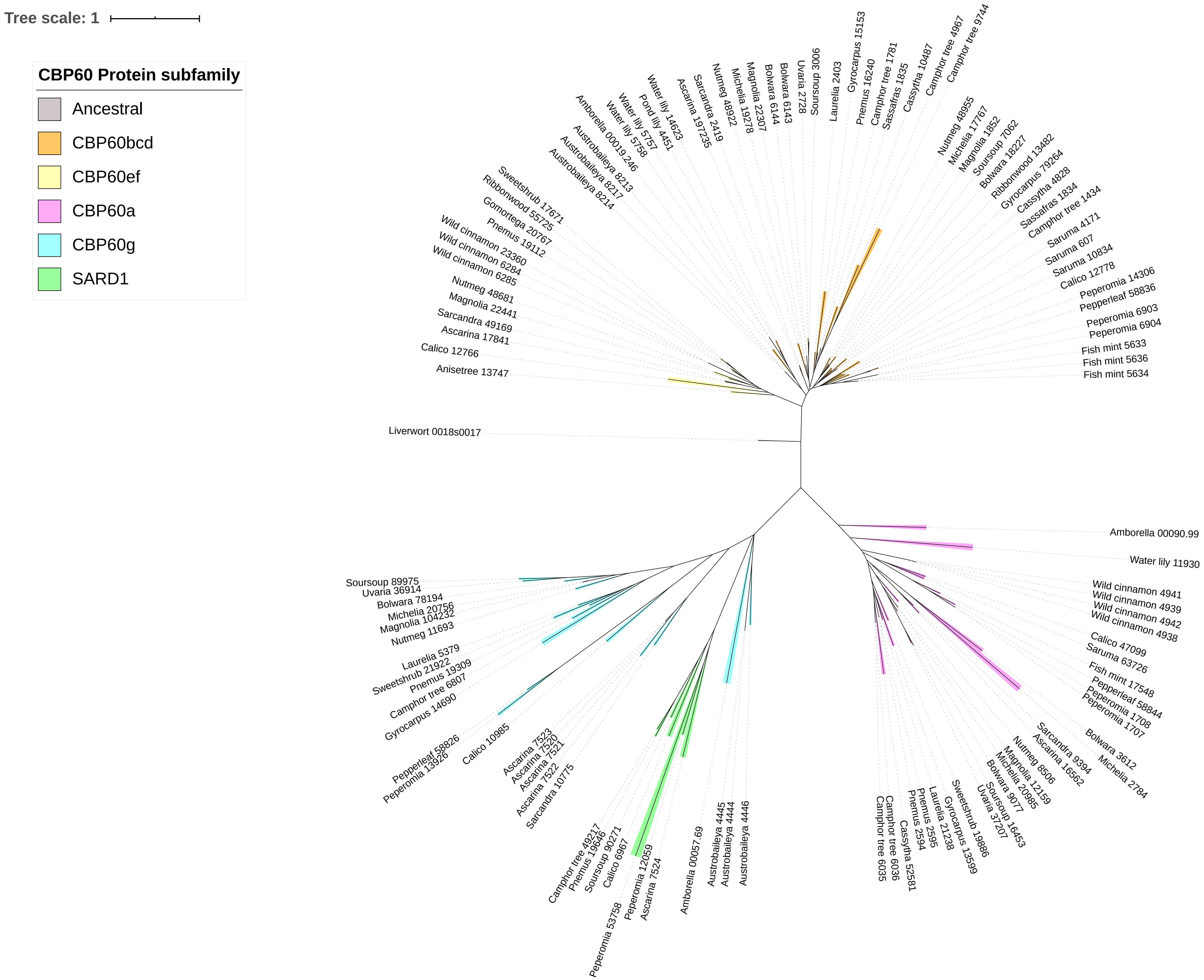

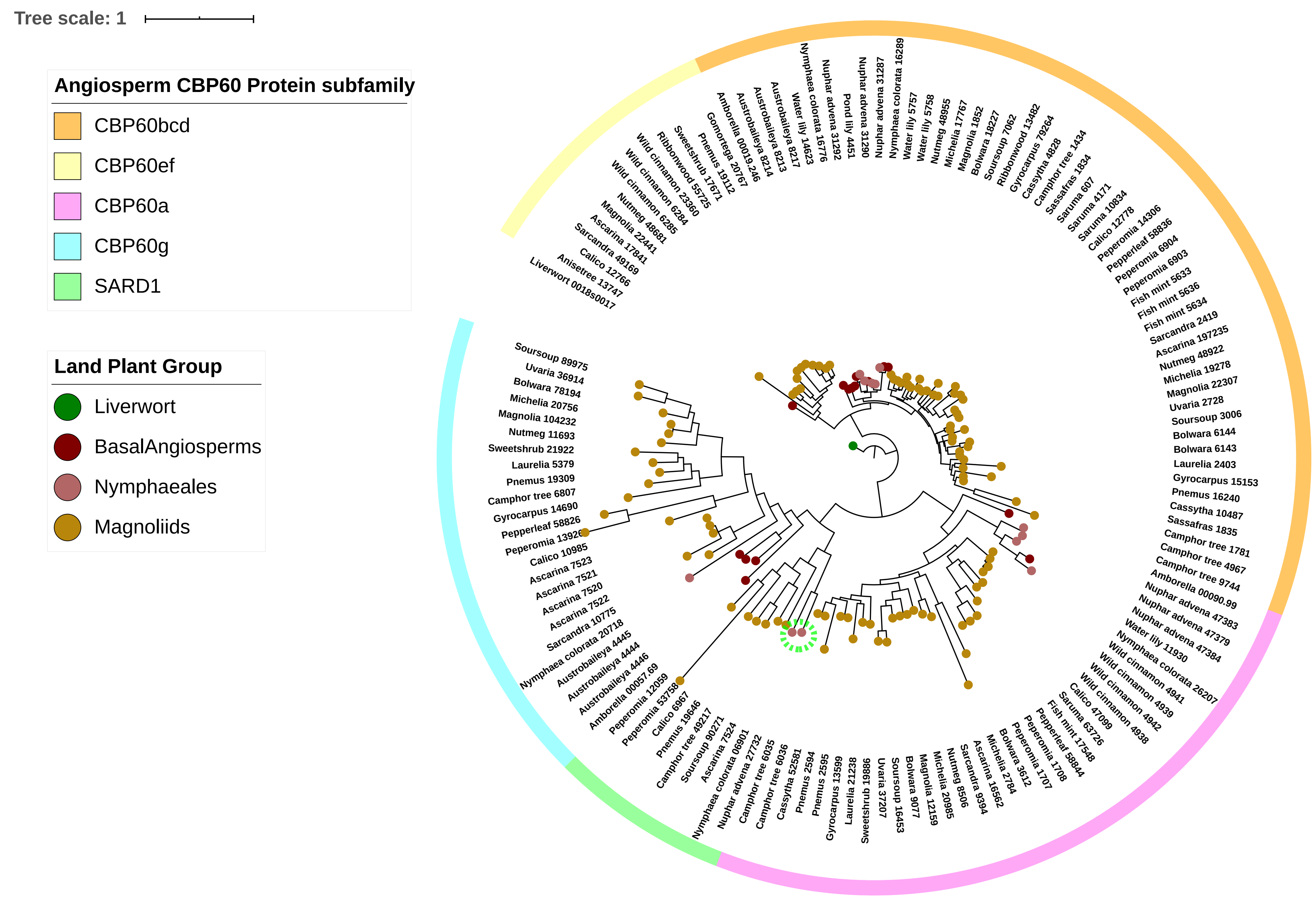

### FigS3.pdf

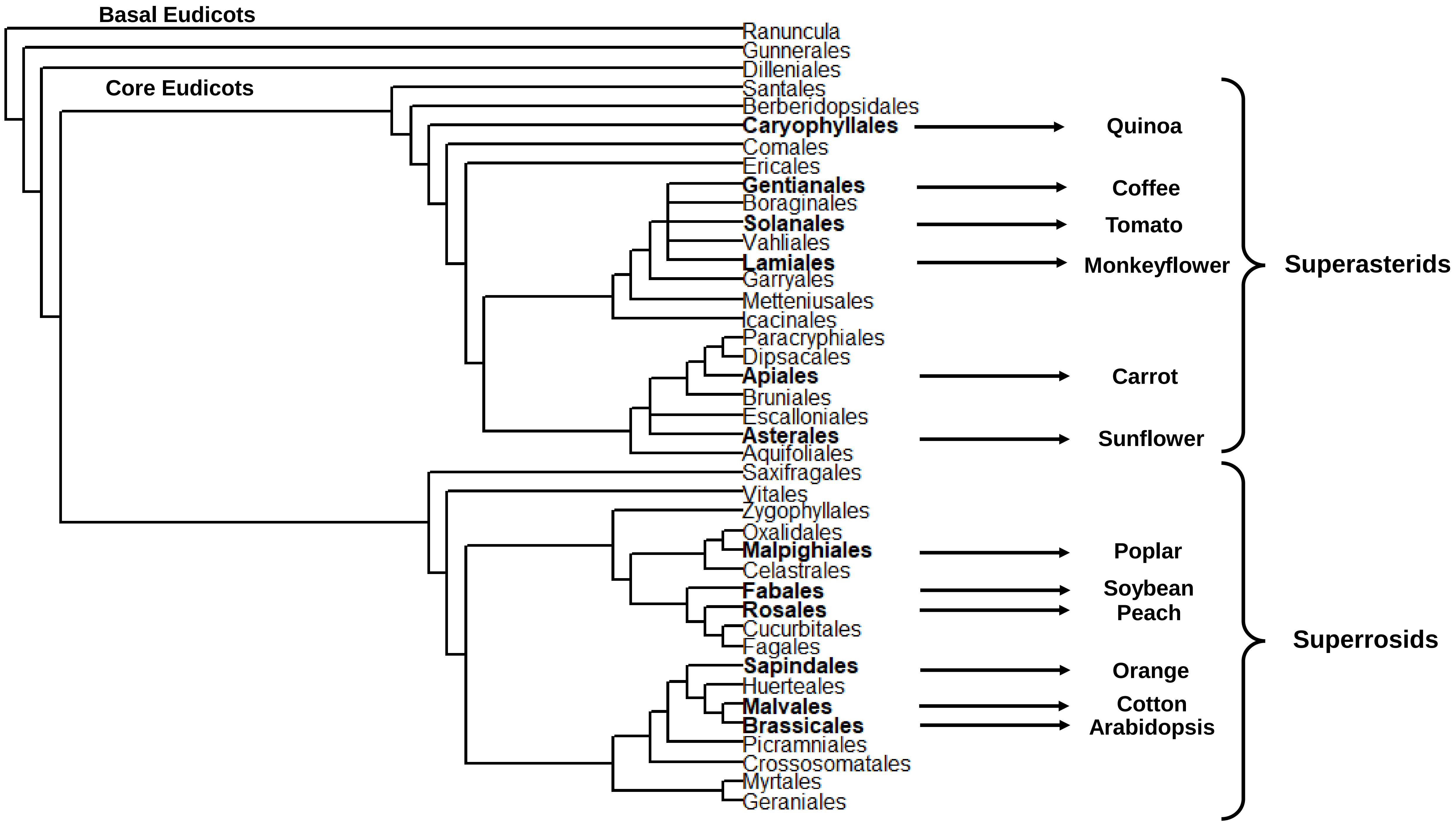

### FigS4.v10.pdf

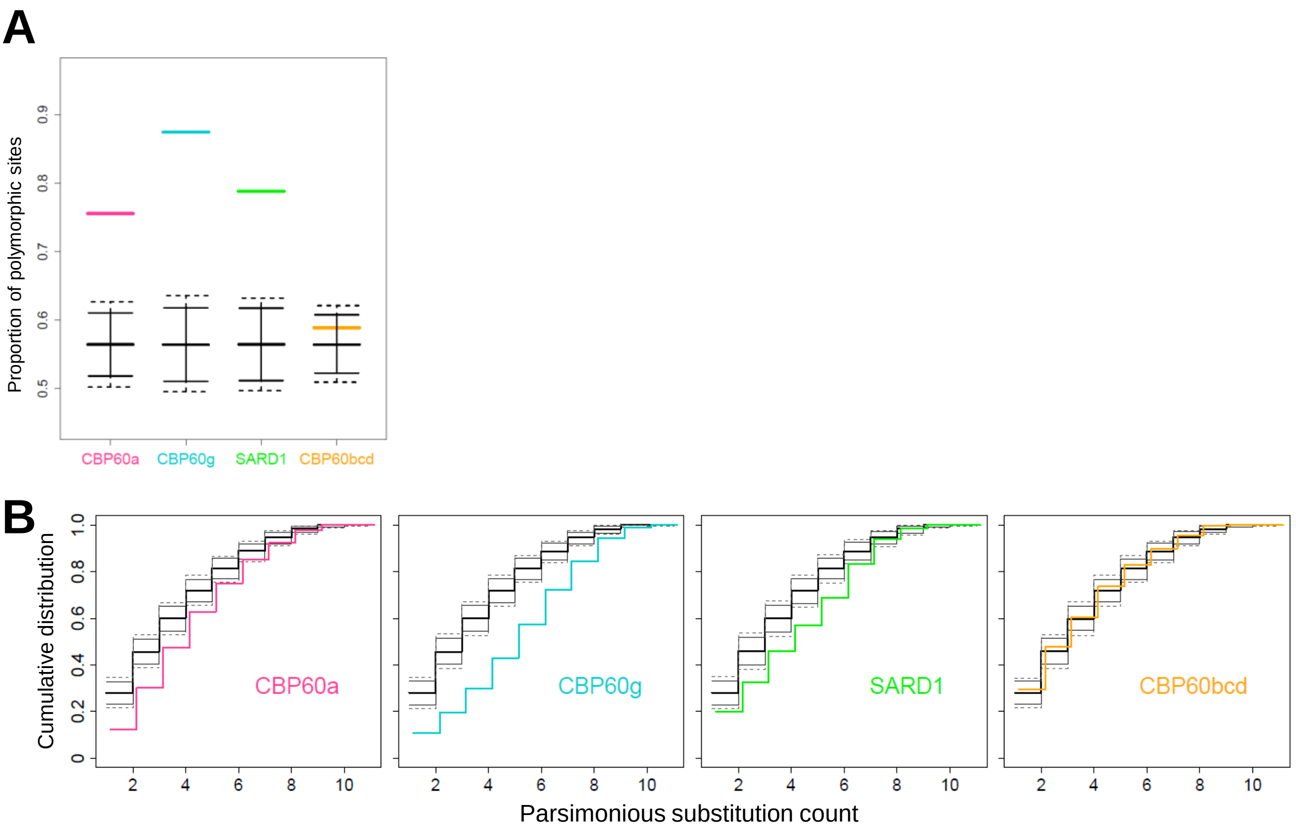

### FigS5.pdf

A. N-terminal to the CBP60-conserved domain

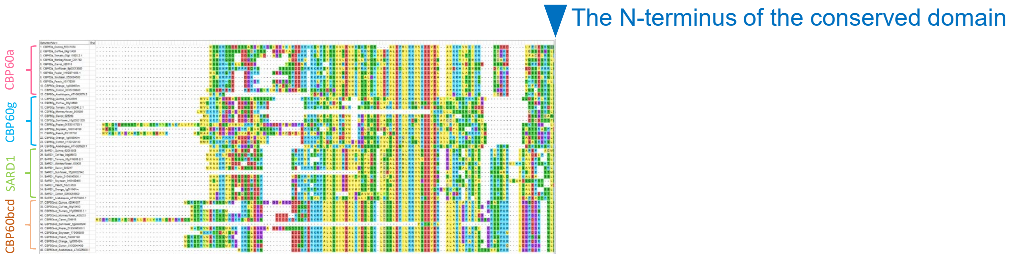


B. C-terminal to the CBP60-conserved domain

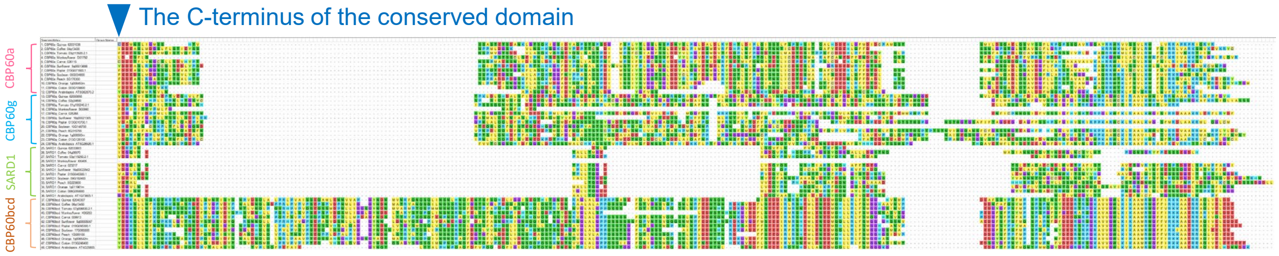

### FigS6.pdf

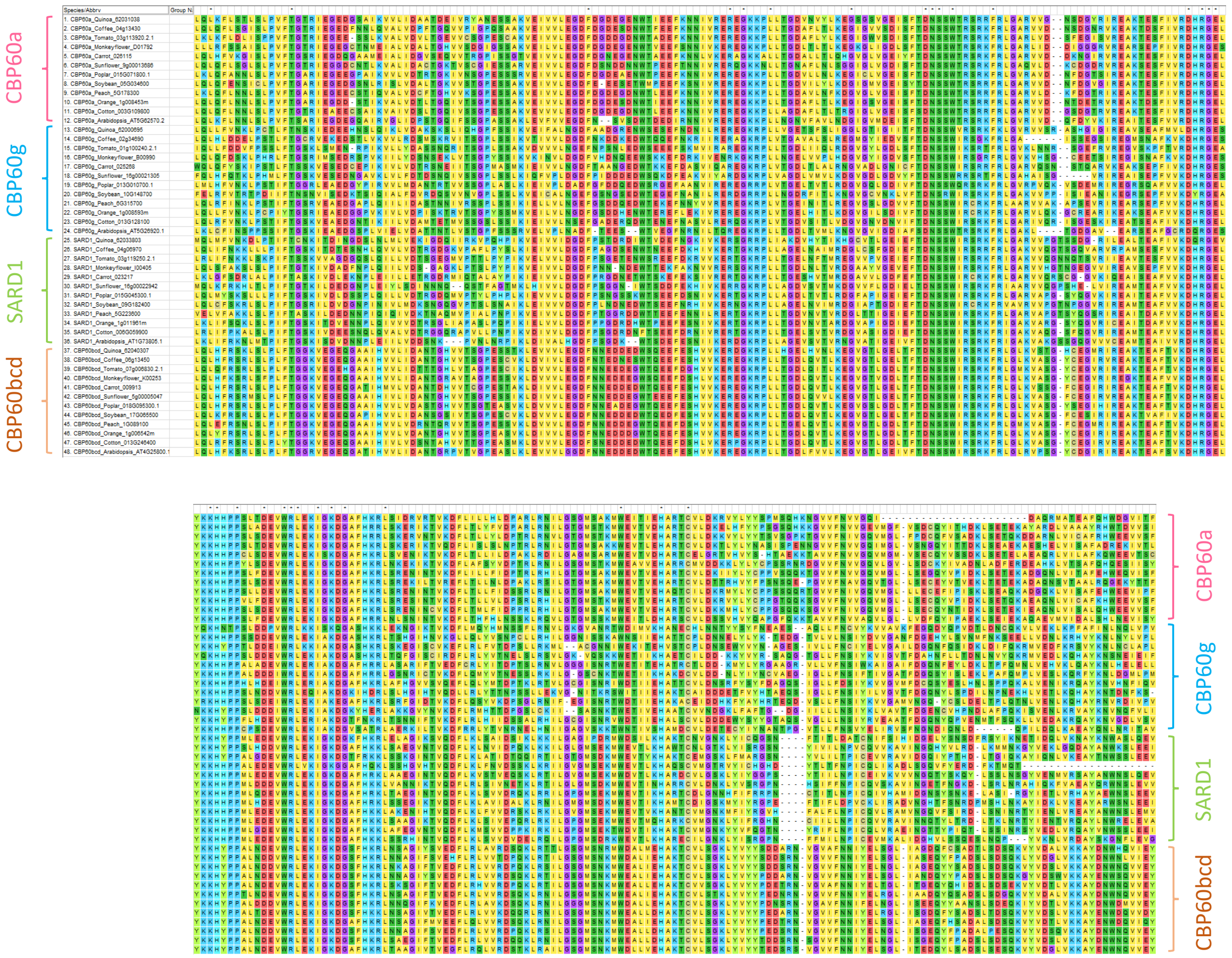

### FigS8.pdf

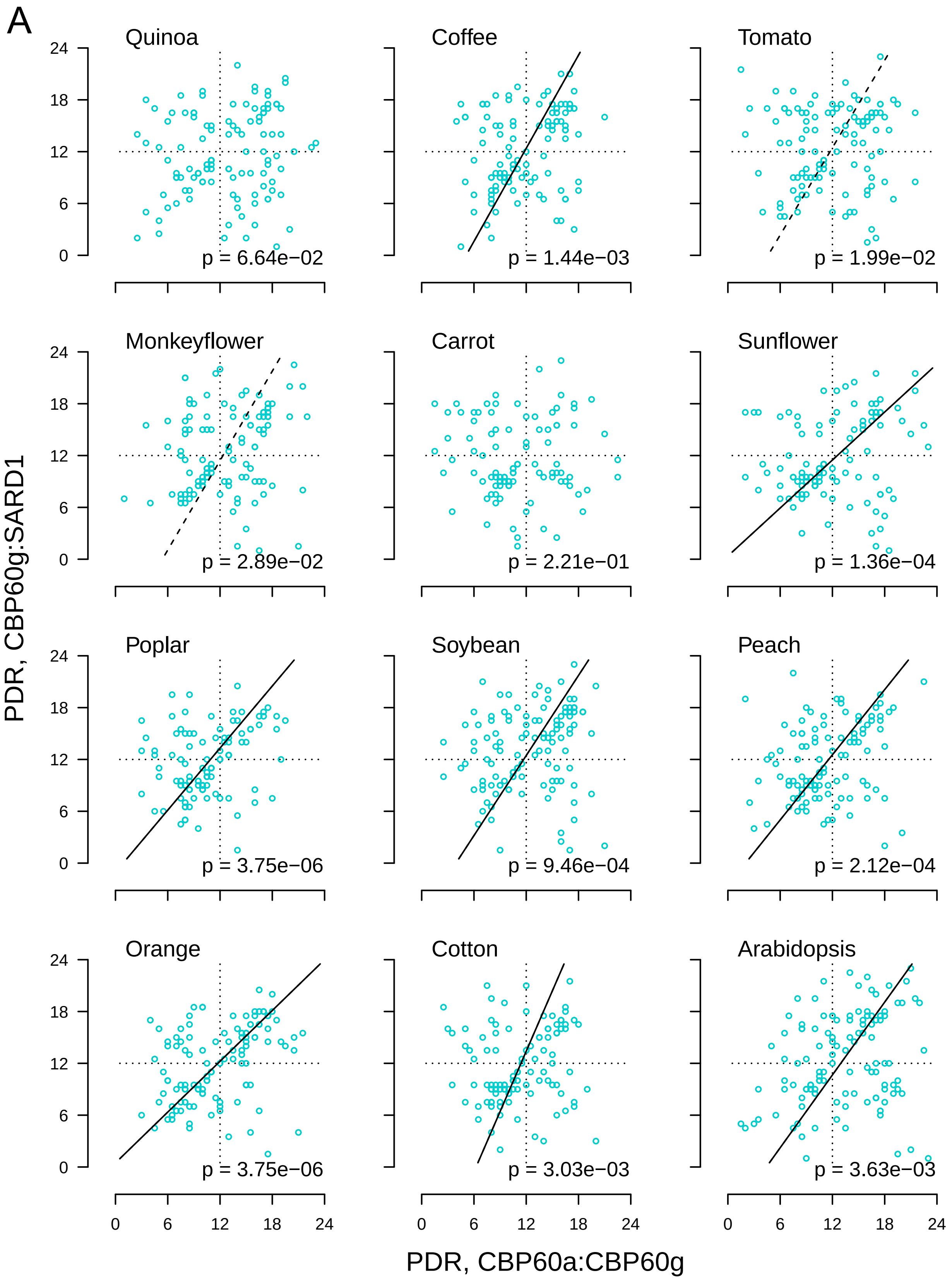

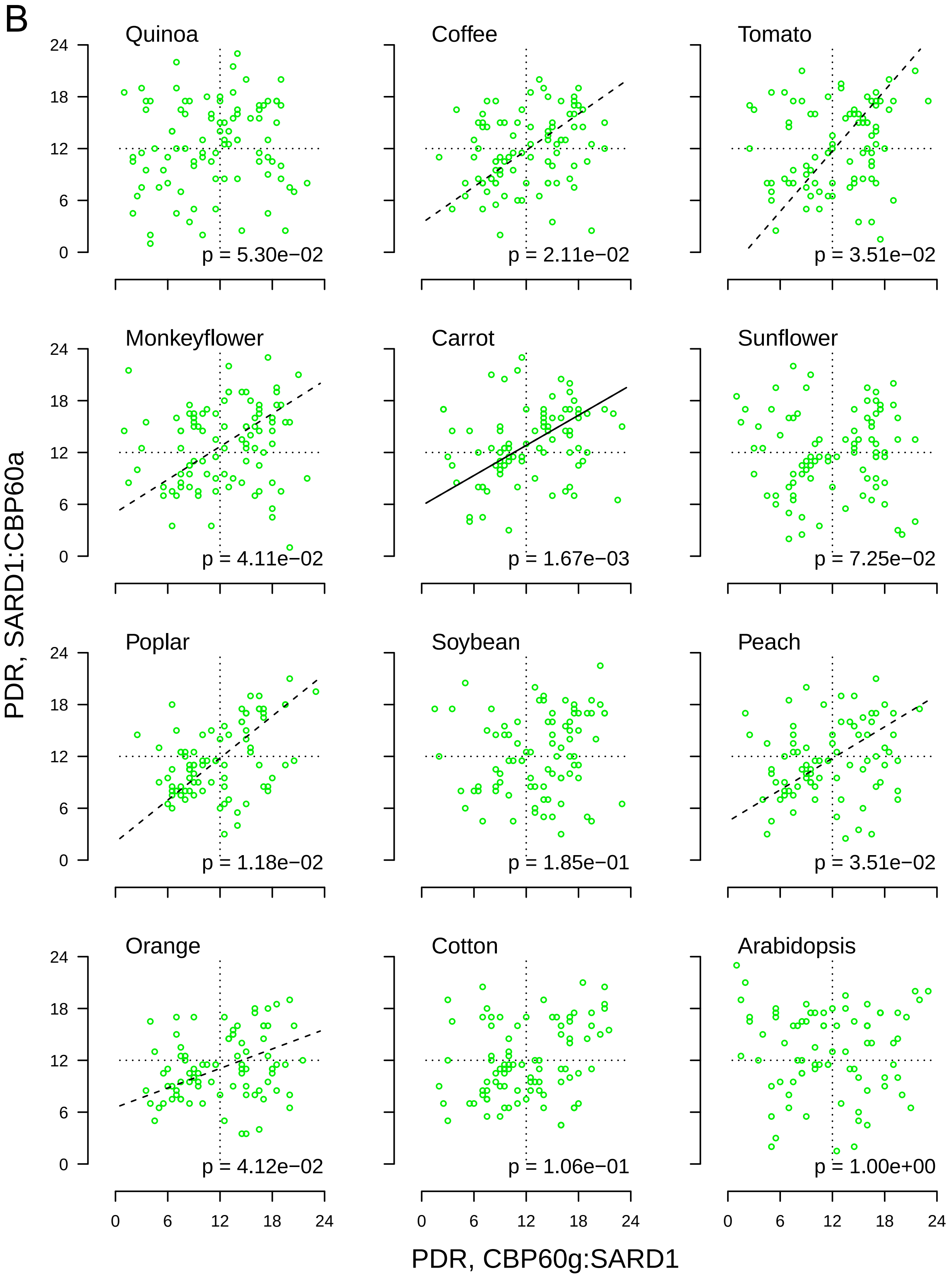

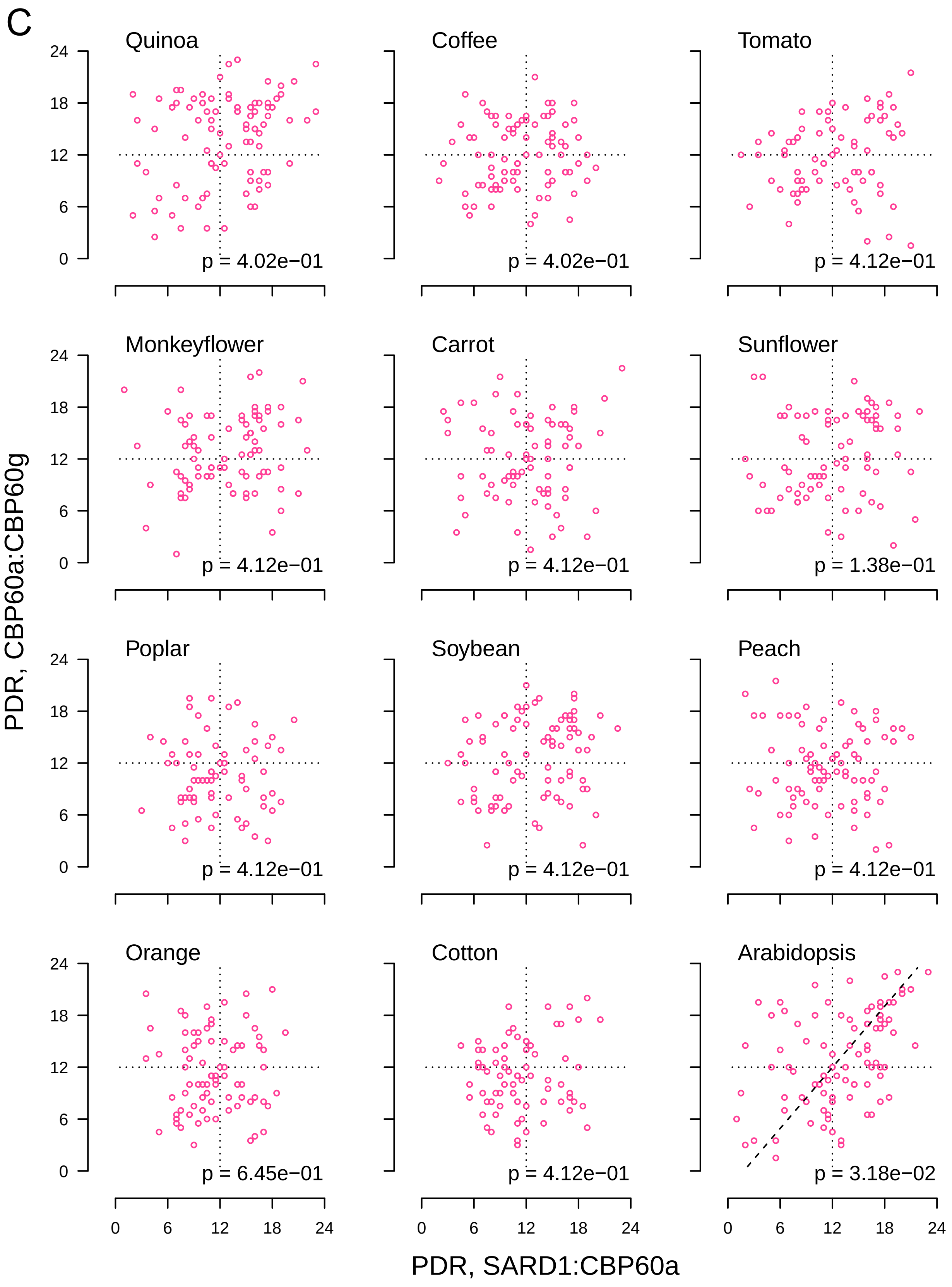

### FigS9.pdf

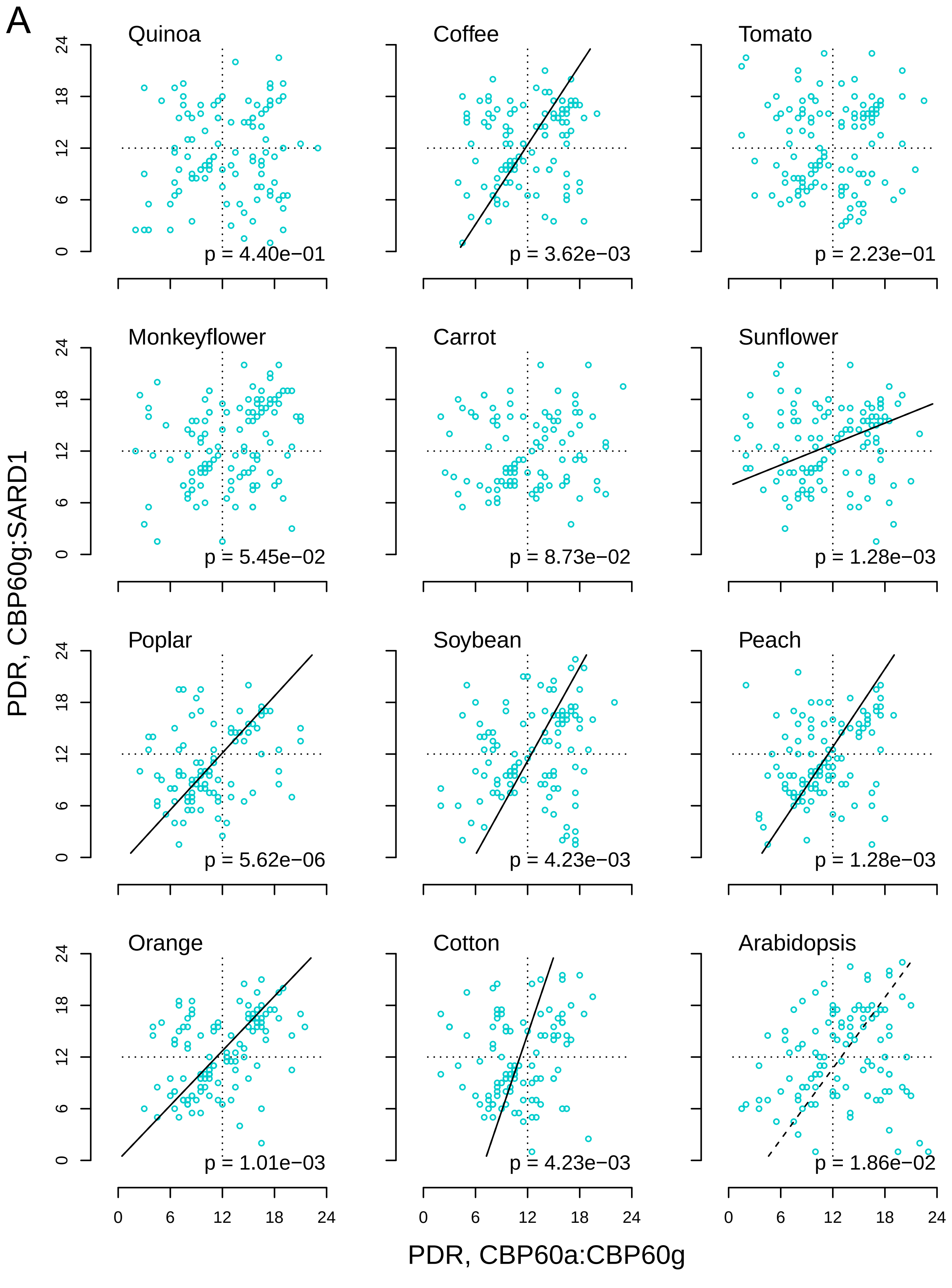

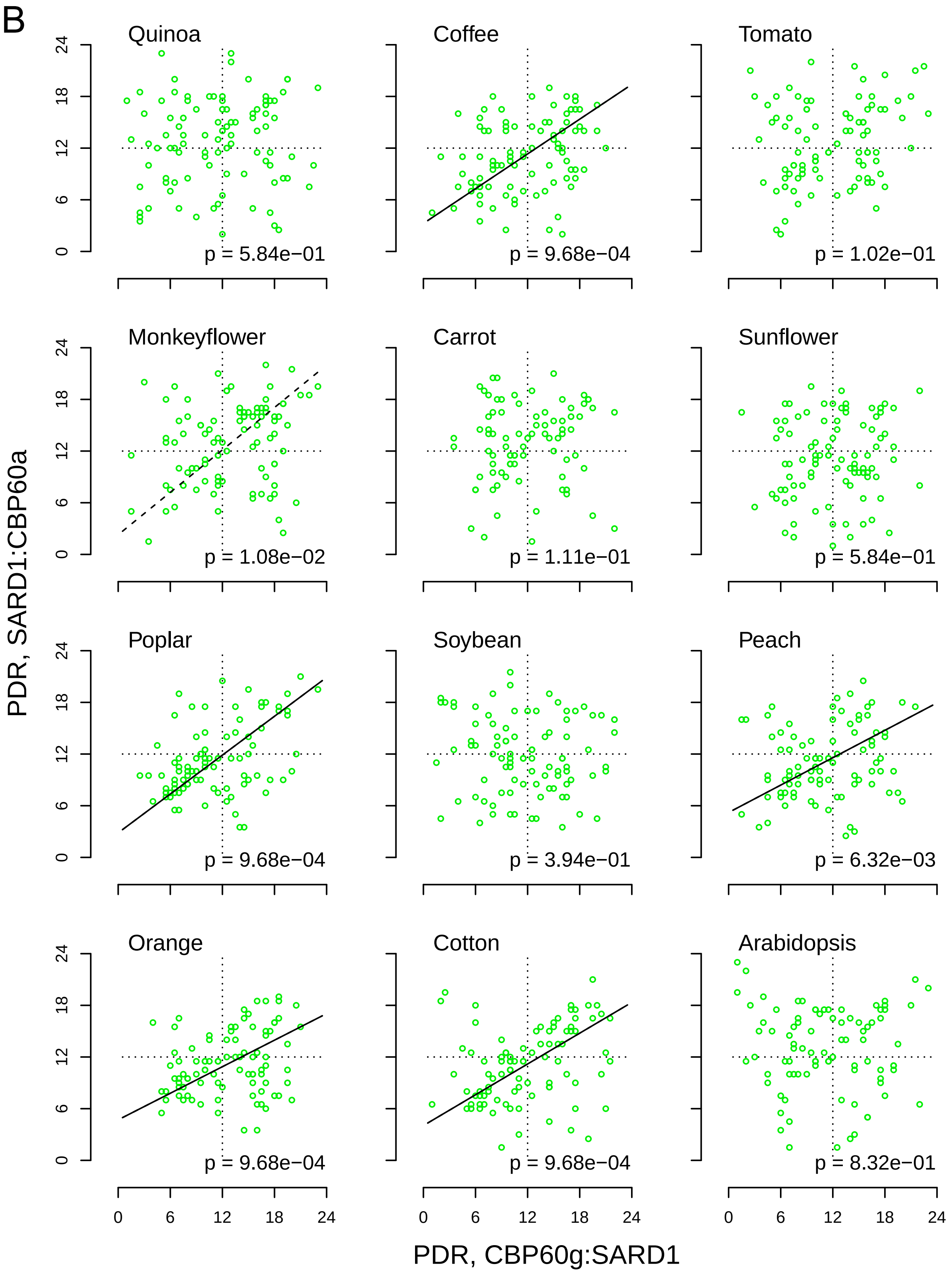

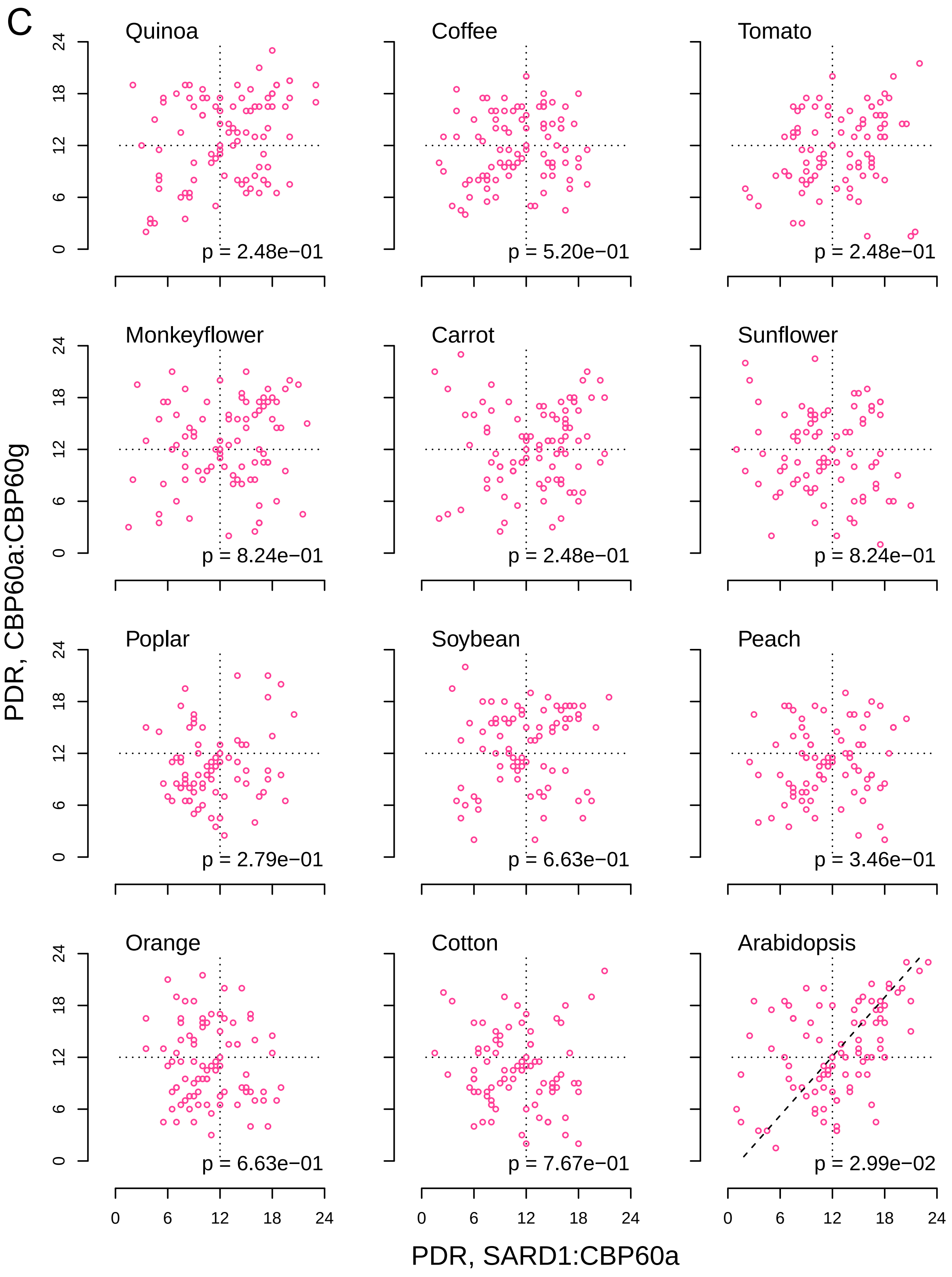

### FigS10.pdf

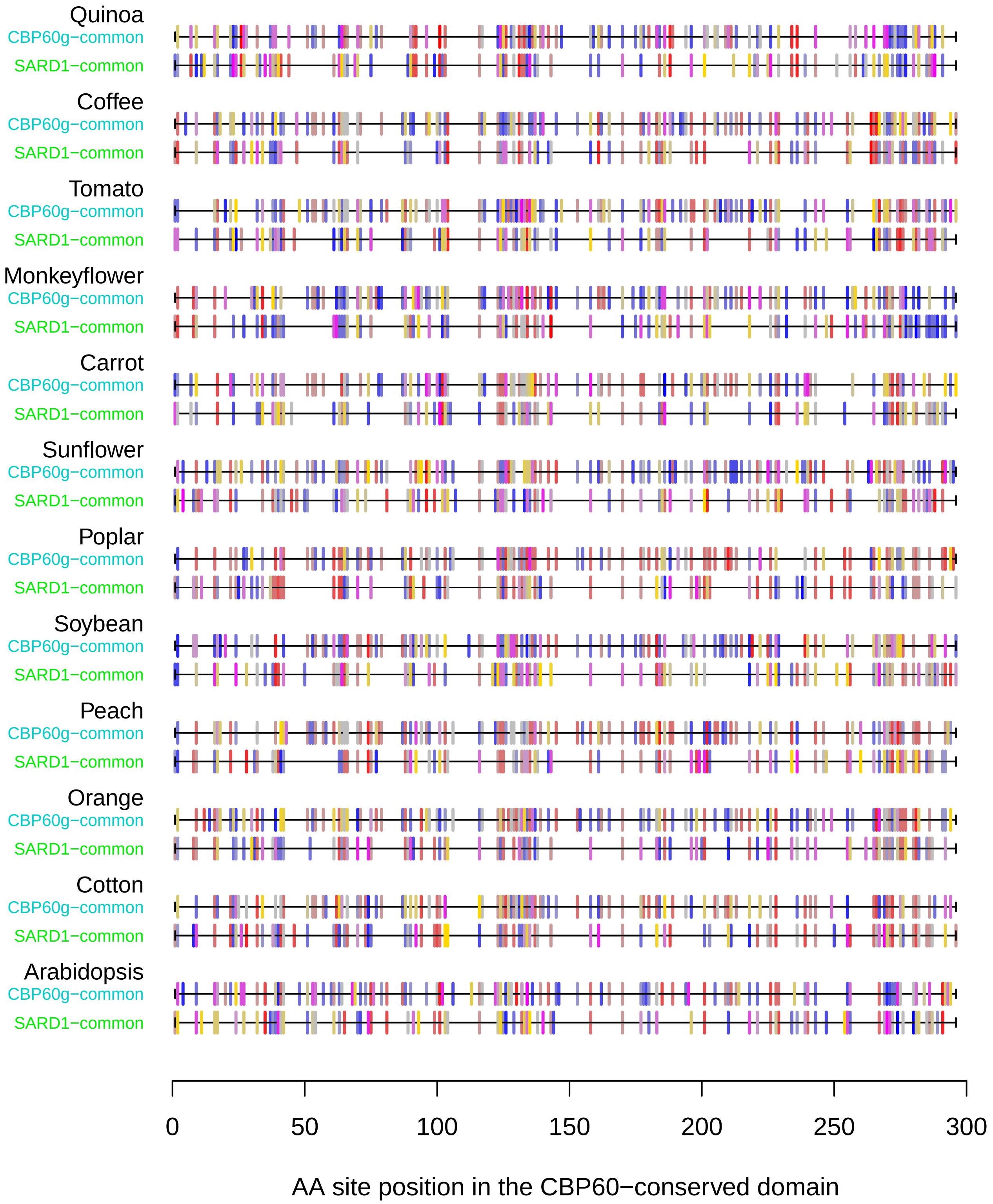

### FigS11.pdf

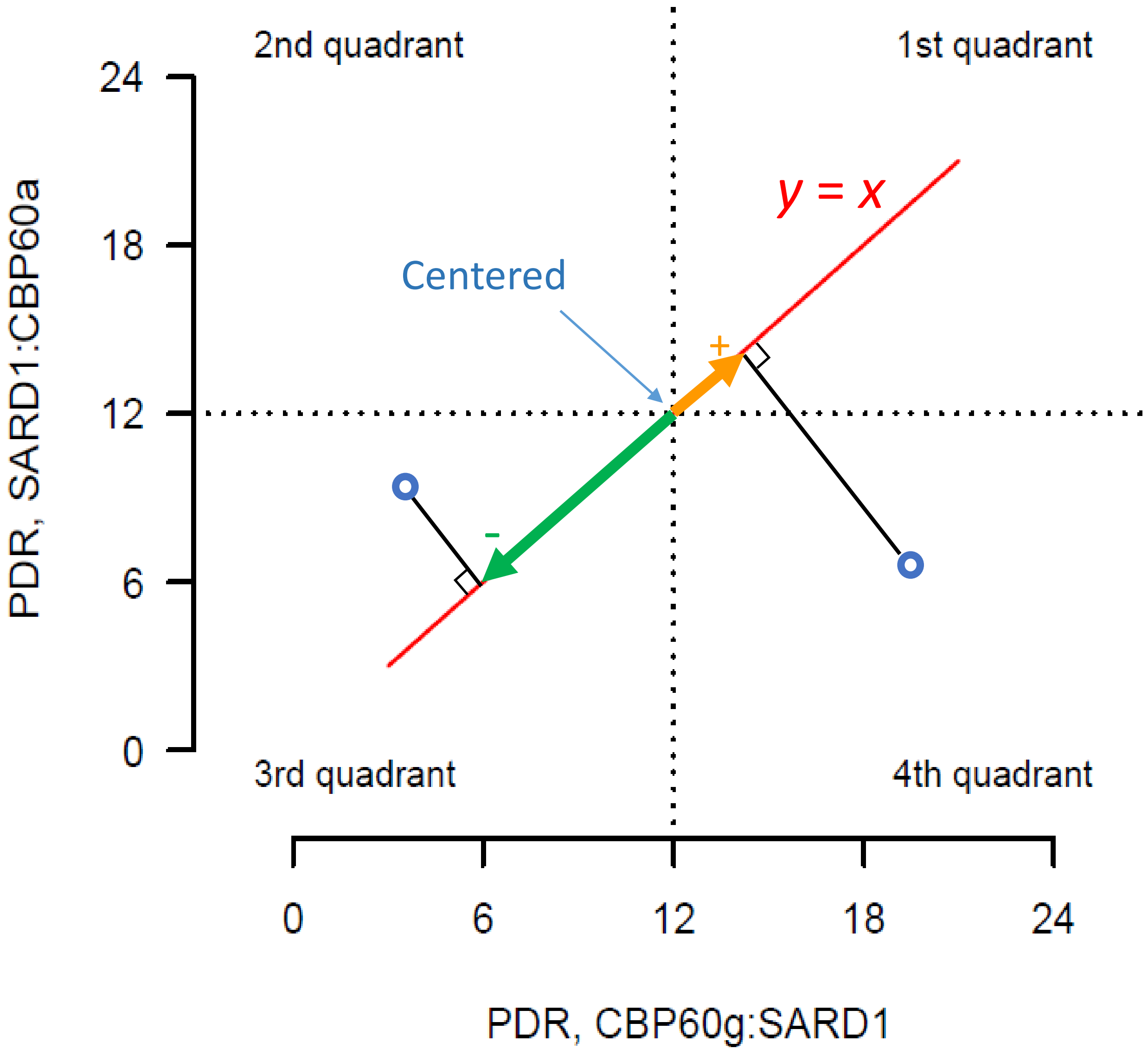

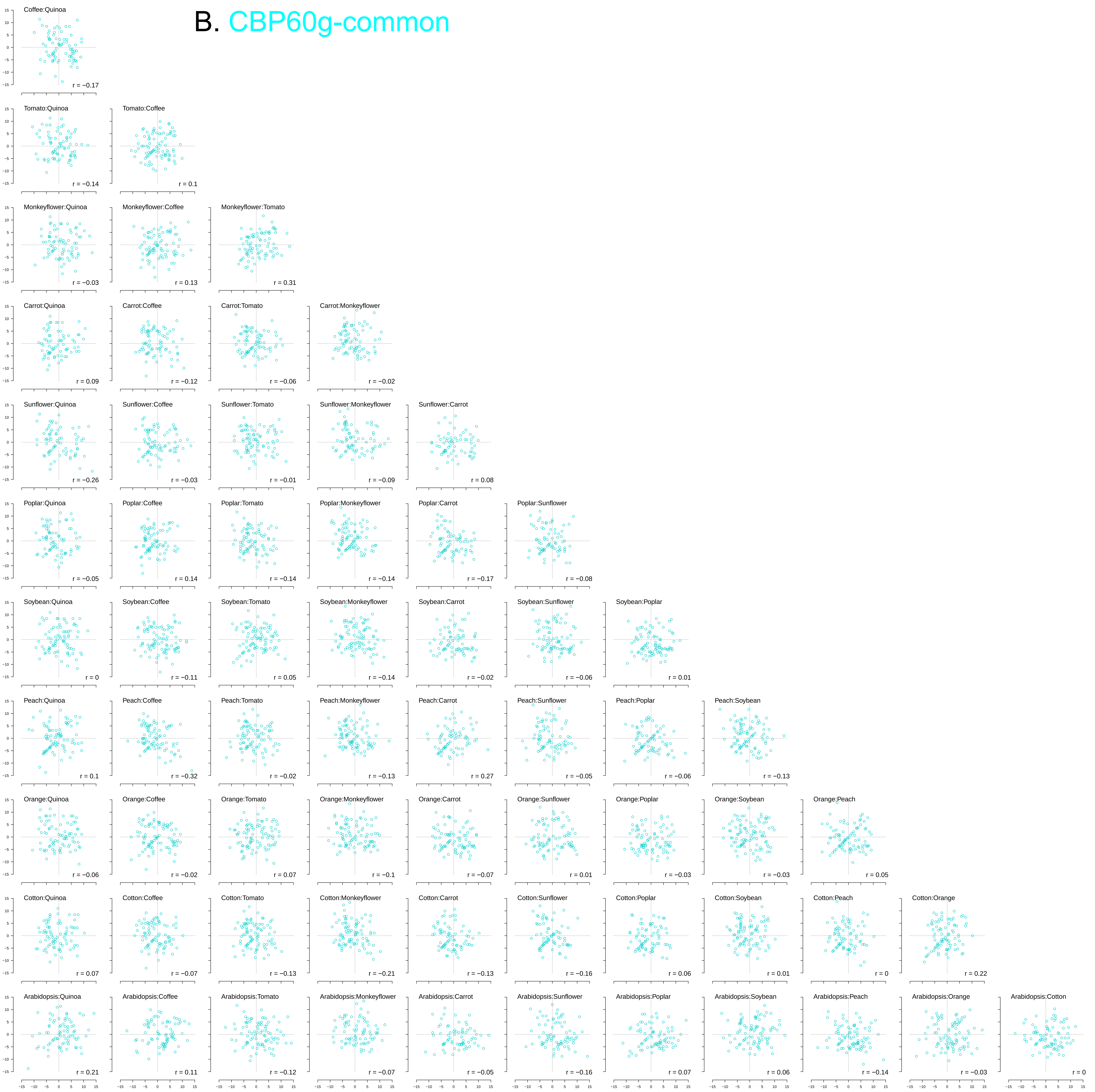

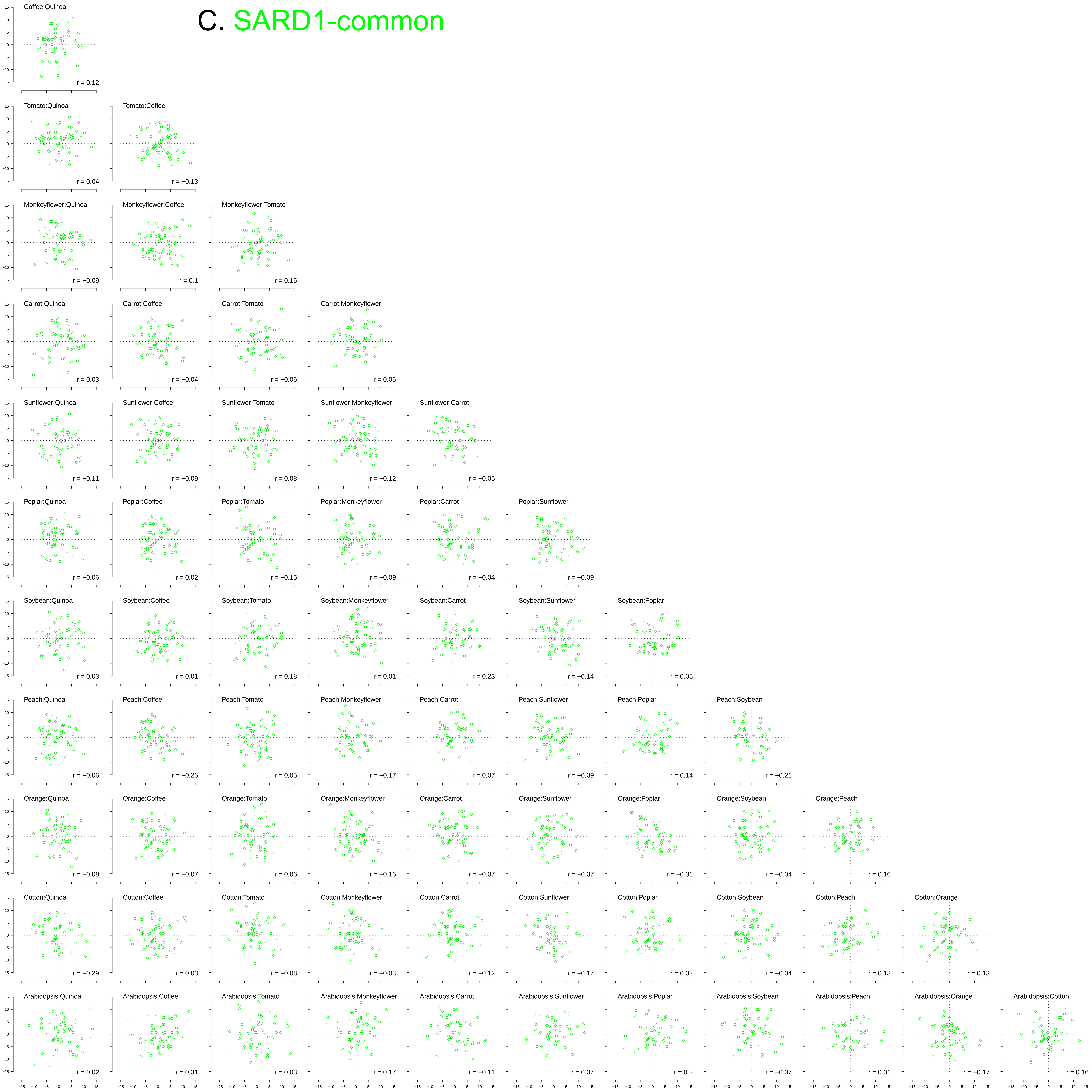

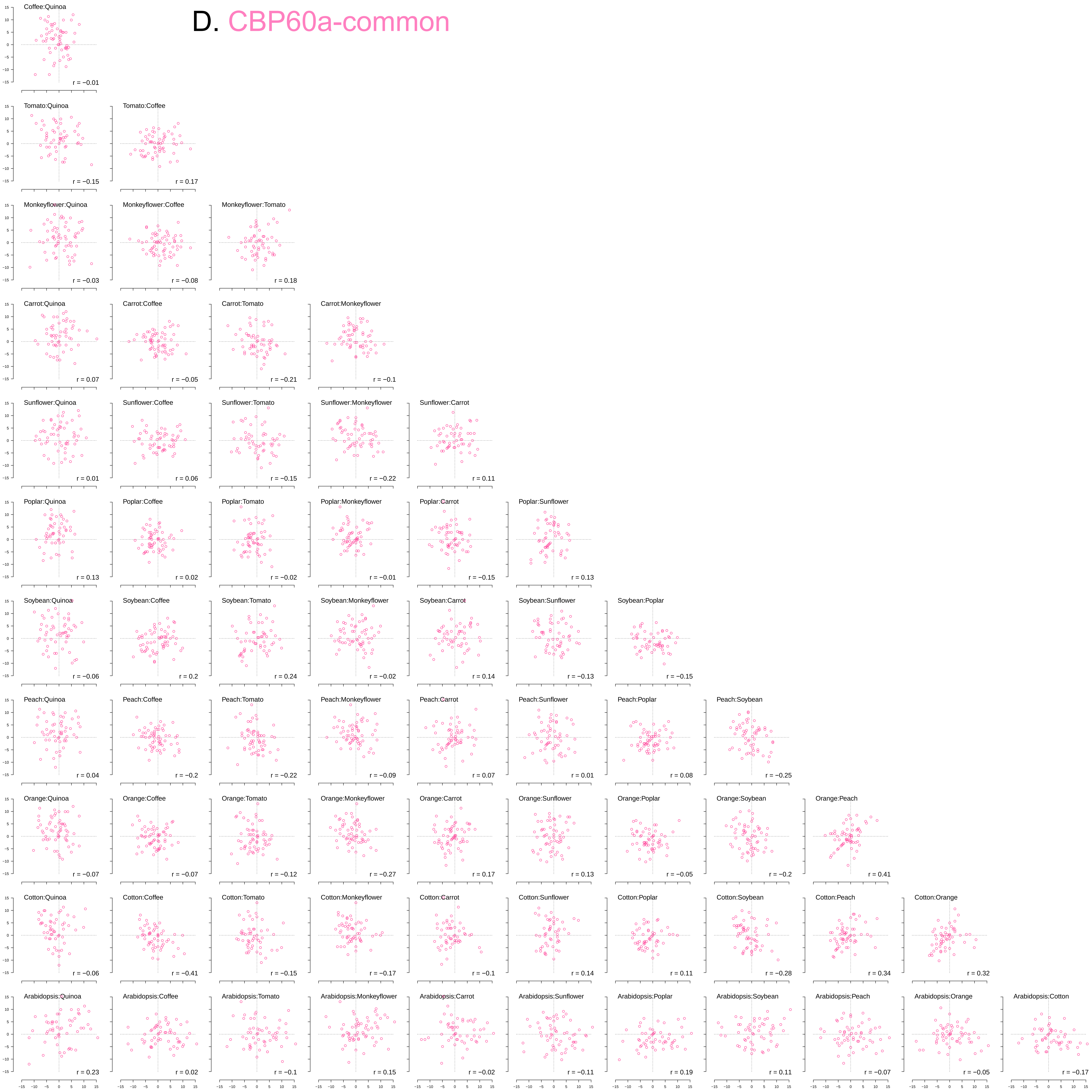

### FigS12.pdf

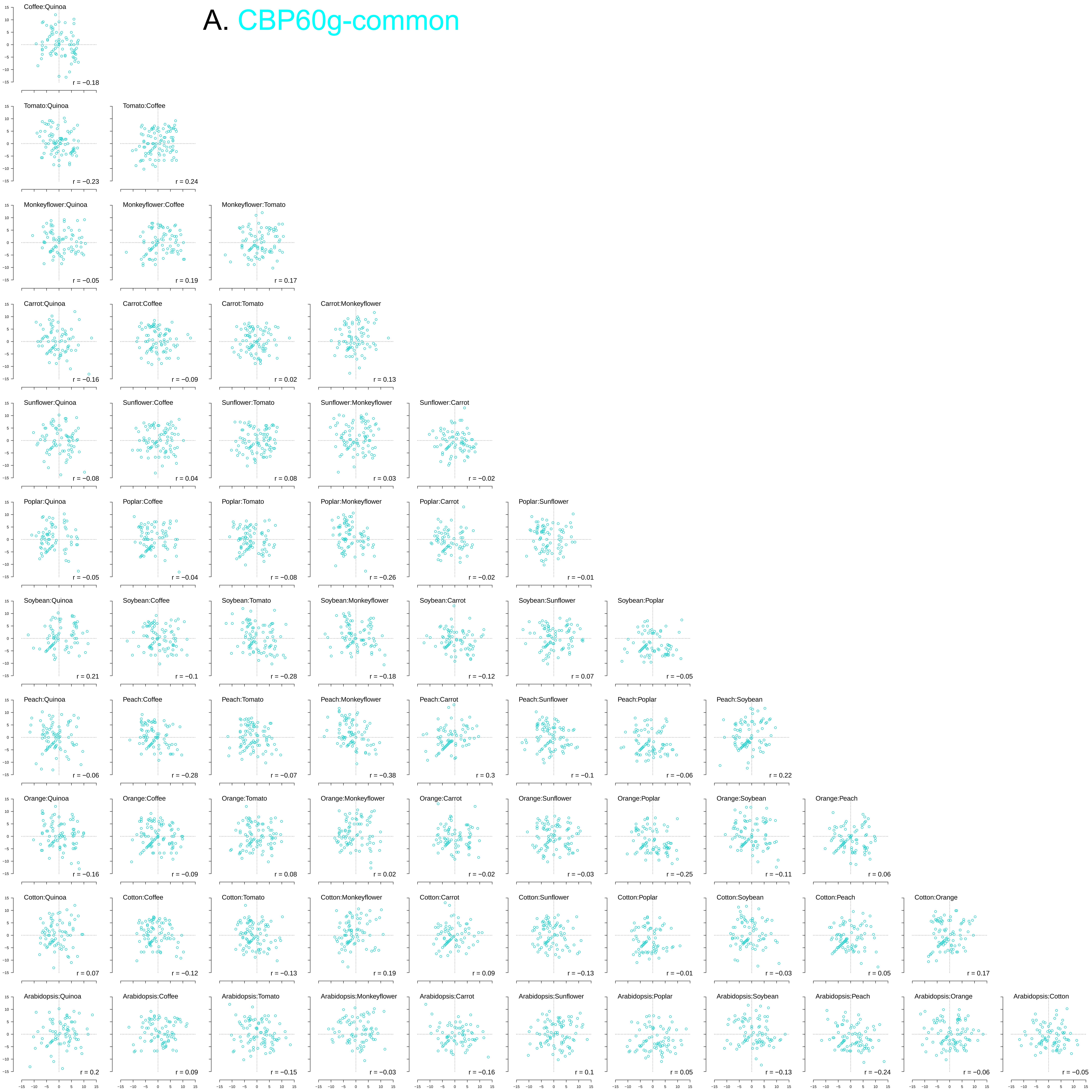

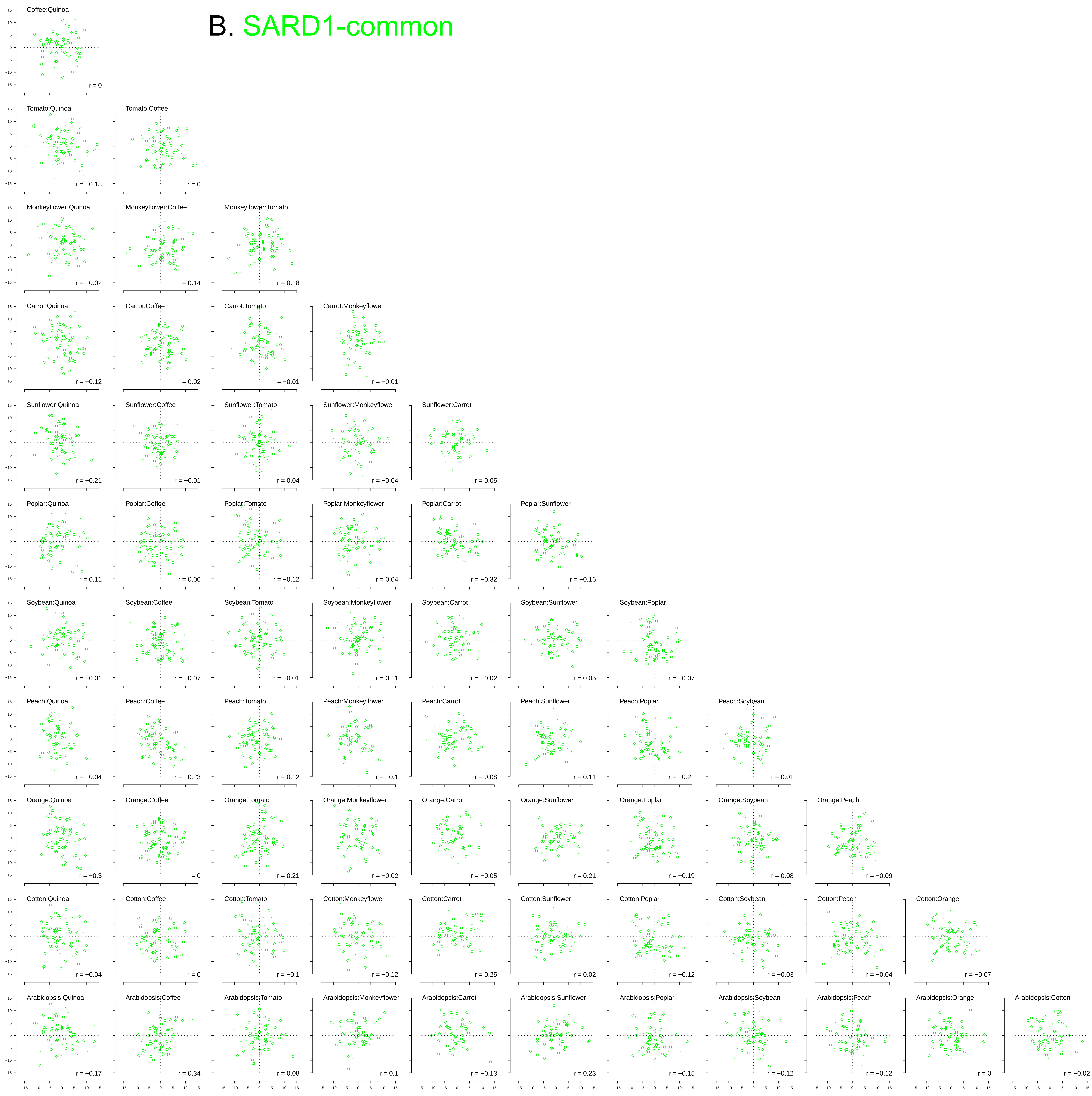

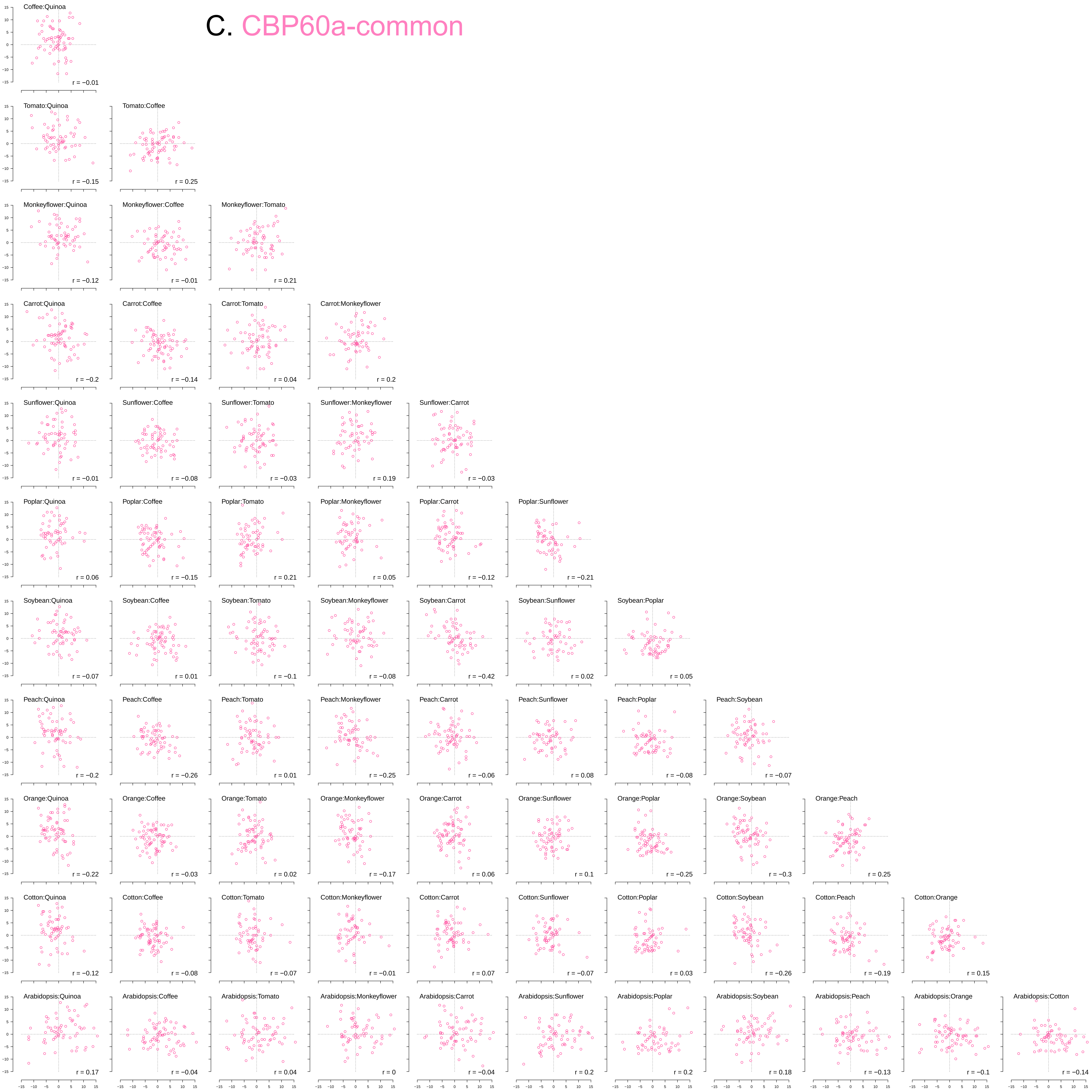

### FigS13.v10.pdf

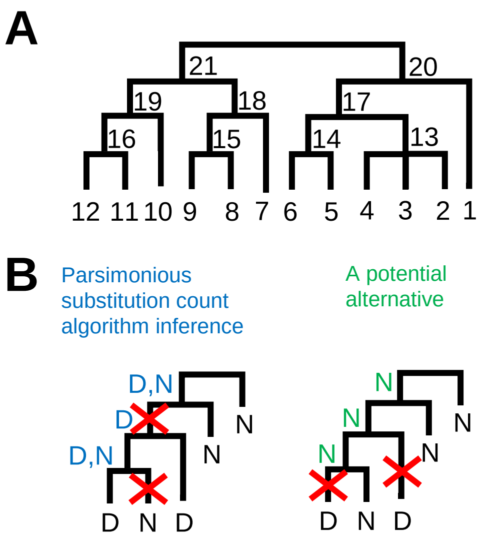
